# Phage DNA polymerase prevents deleterious on-target DNA damage and enhances precise CRISPR/Cas9 editing

**DOI:** 10.1101/2023.01.10.523496

**Authors:** Qiaoyan Yang, Jonathan S. Abebe, Michelle Mai, Gabriella Rudy, Sang Y. Kim, Orrin Devinsky, Chengzu Long

**Affiliations:** NYU Cardiovascular Research Center, Leon H. Charney Division of Cardiology, Department of Medicine, NYU Langone Health, New York, NY, USA; Department of Pathology, NYU Langone Health, New York, NY, USA; New York University Langone Comprehensive Epilepsy Center, NYU Langone Health, New York, NY, USA

## Abstract

Common unintended chromosomal alterations induced by CRISPR/Cas9 in mammalian cells, particularly on-target large deletions and chromosomal translocations present a safety challenge for genome editing. Base editing and prime editing that can precisely introduce desired edits without double-stranded breaks and exogenous DNA templates face their own challenges. Thus, there is still an unmet need to develop safer and more efficient editing tools. We screened diverse DNA polymerases of distinct origins and identified T4 DNA polymerase derived from phage T4 that greatly prevents undesired on-target large deletions and chromosomal translocations while increasing the proportion of precise 1- to 2-base-pair insertions generated during CRISPR/Cas9 editing (termed CasPlus). CasPlus induced substantially fewer on-target large deletions while increasing the efficiency to correct common frameshift mutations in *DMD* (exon 52 deletion) and restored higher level of dystrophin expression than Cas9-alone in human induced pluripotent stem cell-derived cardiomyocytes. Moreover, CasPlus can greatly reduce the frequency of on-target large deletions in mouse germline editing. In multiplexed guide RNAs mediating gene editing, CasPlus represses chromosomal translocations while maintaining gene disruption efficiency that is higher or comparable to Cas9 in primary human T cells. Therefore, CasPlus offers a safer and more efficient gene editing strategy to treat pathogenic variants or to introduce genetic modifications in human applications.

## Introduction

The engineered clustered regularly interspaced short palindromic repeats /CRISPR-associated protein 9 (CRISPR/Cas9)-mediated genome editing has revolutionized genetics. (Cong, Ran et al., 2013, Jinek, Chylinski et al., 2012, Jinek, East et al., 2013, Mali, Yang et al., 2013). However, Cas9 can generate undesired on-target large deletions(Kosicki, Tomberg et al., 2018, Nahmad, Reuveni et al., 2022a), chromosomal translocations(Stadtmauer, Fraietta et al., 2020), chromothripsis(Leibowitz, Papathanasiou et al., 2021), and other complex chromosomal rearrangements,(Boutin, Cappellen et al., 2022) as well as off-target effect. Although numerous strategies have been developed to minimize CRISPR/Cas9-mediated off-target effects(Uddin, Rudin et al., 2020), few approaches can mitigate collateral on-target DNA damage(Bothmer, Gareau et al., 2020, Xin, Yin et al., 2022, Yin, Lu et al., 2022, Yoo, Yadav et al., 2022). Base editing and prime editing are CRISPR/Cas-based gene editing technologies which can efficiently and precisely edit genes, but both have limitations. Base editing is not designed to mediate indels and has off-target effects on DNA and RNA. Prime editing has lower editing efficiency than Cas9 when targeting identical genome sites(Anzalone, Koblan et al., 2020, Anzalone, Randolph et al., 2019, Gaudelli, Komor et al., 2017, Kim, Yu et al., 2021, Komor, Kim et al., 2016). Prime editing efficiency and fidelity are inhibited by DNA mismatch repair (MMR), which is a highly conserved DNA repair pathway and maintains genomic stability(Ferreira da Silva, Oliveira et al., 2022). Clinical trials with CRISPR/Cas9 therapies are underway(2023), but new tools to efficiently introduce precise edits with minimal byproducts and on-target damage would further advance the genome editing field.

Cas9 cleaves target DNA producing double-strand breaks (DSBs) with blunt ends or staggered ends with 5′ overhangs(Shi, Shou et al., 2019). Repairing these ends without exogenous templates usually occurs via non-homologous end joining (cNHEJ) or microhomology-mediated end joining (MMEJ)(Chang, Pannunzio et al., 2017). The specific repair pathway determines Cas9 editing outcomes. MMEJ repair requires an end resection and microhomology sequences, and often results in deletions. MMEJ is associated with Cas9-induced on-target large deletions, chromosome translocations and rearrangement(Kosicki, Allen et al., 2022, Owens, Caulder et al., 2019, Sfeir & Symington, 2015). cNHEJ repair directly joins two compatible ends without or with minor end resection, leading to small insertions or deletions (indels)(Mao, Bozzella et al., 2008, Sfeir & Symington, 2015). Comprehensive analyses on Cas9 on-target edits reveal small insertions produced by exogenous template-free Cas9 editing are precise and predictable(Allen, Crepaldi et al., 2018, Leenay, Aghazadeh et al., 2019, Shen, Arbab et al., 2018). The frequency and pattern of Cas9-generated small insertions depend on the local sequences surrounding the Cas9 cut site (templated insertions)(Shi et al., 2019). Current tools cannot fully control the outcomes of 1- to 2-base-pair (bp) insertions, which can be used to reframe or knockout genes(Bermudez-Cabrera, Culbertson et al., 2021). In budding yeast, Cas9-induced 1–3-bp insertions are attributed to DNA polymerase 4 (Pol 4)(Lemos, Kaplan et al., 2018). Thus, systematic screening gap-filling DNA polymerases may identify enzymes that favor cNHEJ-mediated small insertions during Cas9 editing in mammalian cells. The competition between fill-in and end resection processes in DSB with staggered ends(Cejka & Symington, 2021, Kosicki et al., 2022) suggests that gap-filling DNA polymerases may also inhibit MMEJ-mediated on-target large deletions and chromosome translocations.

Duchenne muscular dystrophy (DMD) is characterized by degeneration of cardiac and skeletal muscles(O’Brien & Kunkel, 2001) and results from mutations in the X-linked dystrophin gene (*DMD*)(Muntoni, Torelli et al., 2003). Deletion of one or more exons is the most common mutation type(Duan, Goemans et al., 2021). Dilated cardiomyopathy (DCM) is a common and lethal feature(Adorisio, Mencarelli et al., 2020). We used CRISPR/Cas9 to correct or bypass the *DMD* mutations in cultured human cells and *mdx* mice(Amoasii, Long et al., 2017, Long, Amoasii et al., 2016, Long, Li et al., 2018, Long, McAnally et al., 2014, Olson, 2021, Zhang, Li et al., 2020, Zhang, Li et al., 2022). However, complex rearrangements can arise during *DMD* editing due to the repetitive elements, stem-loop structures, and preexisting mutations in this unstable genomic region(Nelson, Wu et al., 2019, Oshima, Magner et al., 2009). Undesired on-target DNA damage at edited *DMD* sites, a safety concern in human therapy, were not addressed. CRISPR/Cas9 engineered Chimeric antigen receptor (CAR) T-cell therapy may overcome conventional CAR-T therapy problems, like Graft-versus-host disease (GvHD), CAR-T cell exhaustion and limited patient T cell numbers(Depil, Duchateau et al., 2020, Khan & Sarkar, 2022, Labanieh & Mackall, 2023, Sheridan, 2022b). However, unintended chromosomal abnormalities such as translocations, pose a safety risk with multiplex gene edited patient T cells(Bothmer et al., 2020, Poirot, Philip et al., 2015, Stadtmauer et al., 2020).

Here, we identified a phage DNA polymerase that markedly reduces the on-target large deletions and chromosomal translocations associated with regular Cas9 editing while maintaining an equal or higher editing efficiency of desired products. Our results demonstrate that our Cas9 and DNA Polymerase, termed CasPlus, can more safely and efficiently correct *DMD* mutation in cardiomyocytes and repress chromosomal translocations in human T cells.

## Results

### T4 and RB69 DNA polymerases favor small insertions over deletions during Cas9 editing

First, we aimed to identify DNA polymerases that can enhance the fill-in step, leading to a ligation process favoring small insertions over deletions during Cas9 editing in mammalian cells (**Fig 1A**). We established a stable HEK293T reporter cell line expressing a mutant tdTomato gene with a 1-bp deletion of adenine (A) at position 151 (tdTomato-del151A) (**Fig 1B**). Next, we investigated the effects of DNA polymerases on Cas9 editing. We constructed MCP (MS2 bacteriophage coat protein)-tagged vectors expressing human codon optimized DNA polymerases which can fill in 5’ overhang, including 1) X-family polymerases Pol λ, Pol μ, and Pol β of human origin, 2) Pol 4 from yeast(Lemos et al., 2018, Ramsden & Asagoshi, 2012), 3) family A polymerases Pol I and Klenow fragment (KF)(Bebenek, Joyce et al., 1990) from bacteria and 4) B-family polymerases T4 DNA pol from bacteriophage T4 (Liu, Tao et al., 2015) (**Fig 1C**). We transfected tdTomato-d151A reporter cells with one vector containing Cas9, green fluorescent protein (GFP), and a tdTomato-targeted single guide RNA (tdTomato-sgRNA) alone or combined with second vector with the DNA polymerases. We sorted transfected cells into populations expressing only GFP (tdTomato^-^GFP^+^) or tdTomato and GFP (tdTomato^+^GFP^+^) and amplified the target DNA by Polymerase chain reaction (PCR) for high-throughput sequencing (HTS) (**Fig 1D and appendix Fig S1A**). Majority of edits observed in tdTomato^+^GFP^+^ populations are reframed 3*n*+1 indels and in tdTomato^-^GFP+ populations are 3*n* or 3*n*+2 indels (**Fig 1E and appendix Fig S1B)**. Very few 3n+2 indels were also observed in tdTomato^+^GFP+ populations, indicating that the tdTomato-d151A reporter cell line might contain multiple copies of reporter gene (**Appendix Fig S1B**). HTS results for the tdTomato^-^GFP^+^ populations showed co-expression of T4 DNA polymerase with Cas9 increased the frequency of 2-bp insertions by ∼6-fold, primarily at the expense of deletions. No other DNA polymerase we tested changed indel profiles compared to Cas9-only control (CTR) (**Fig 1E)**. We obtained similar results by co-expressing RB69 DNA polymerase(Hogg, Cooper et al., 2006), a B-family DNA polymerase with 61% amino acid homology to T4 DNA polymerase, but not with the A-family T7 DNA polymerase(Hori, Mark et al., 1979) (**Fig 1F**). All treatments had a similar ratio of templated to non-templated 2-bp insertions (**Fig 1G**); we found no significant changes among different treatments in tdTomato^+^GFP^+^ populations (**Appendix Fig S1B-E**). Western blot analysis confirmed expression of all DNA polymerases in HEK293T cells (**Appendix Fig S1F**). Experiments then focused on T4 DNA polymerase because it strongly favored small insertions over deletions at the tdTomato-d151A site.

**Figure 1.**
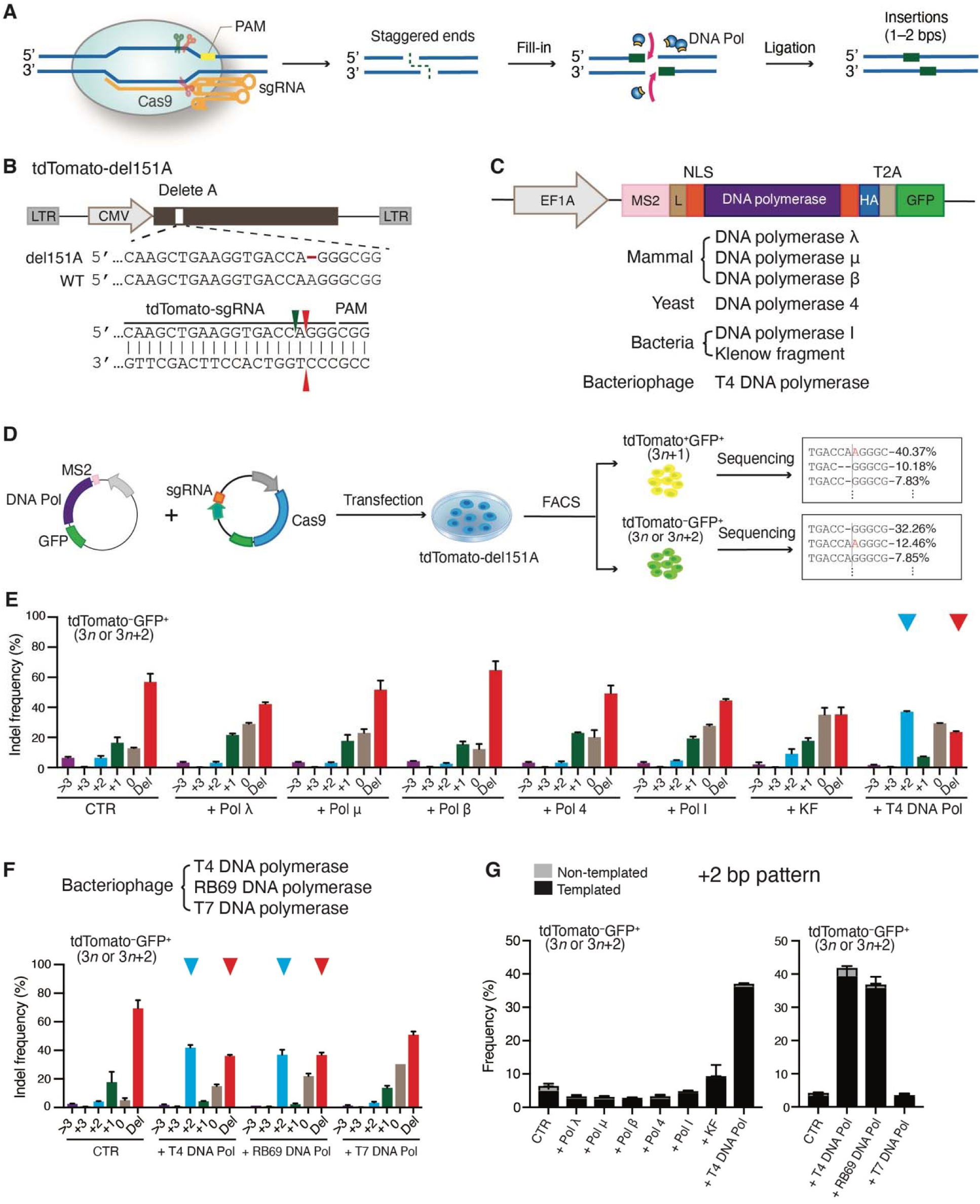
T4 and RB69 DNA polymerases favors small insertions over deletions during Cas9 editing. (**A**) Schematic showing the filling-in of Cas9-induced staggered ends by exogenous DNA polymerase. (**B**) Schematic of lentiviral vector containing tdTomato-d151A (top) and the gRNA used to target it (bottom). Arrowheads, cut sites; red dash, missing nucleotide; LTR, long terminal repeat. (**C**) Architecture of vectors expressing distinct DNA polymerases (see methods). (**D**) Workflow of the DNA polymerase selection process in tdTomato-d151A reporter cells. (**E**-**F**) Frequencies of Cas9-induced indels in tdTomato^-^GFP^+^ populations with or without co-expression of a distinct DNA polymerase. (**G**) Patterns of 2-bp insertions in **E** and **F**.

To test whether tethering of T4 DNA polymerase at Cas9-induced DSBs could further improve 2-bp insertions at the tdTomato-d151A site, we first fused T4 DNA polymerase to the N-terminus or C-terminus of Cas9 via a flexible linker. Unlike co-expression of Cas9 and T4 DNA polymerase, Cas9 fused T4 DNA polymerase proteins (T4-L-SpCas9 and Cas9-L-T4) strongly induced deletions at the protospacer-adjacent motif (PAM) distal region in the tdTomato^-^GFP^+^ and tdTomato^+^GFP^+^ populations (**Appendix Fig S1G-I**). T4 DNA polymerase possesses a polymerase activity that extends the polynucleotide chain, and a 3′-to-5′ proofreading exonuclease activity that excises erroneously incorporated nucleotides (Capson, Peliska et al., 1992). Hence, we hypothesized that, when T4 DNA polymerase fused to Cas9, its exonuclease domain, but not DNA polymerase domain, affects the DSB repair. We deactivated the exonuclease domain of T4 DNA polymerase in T4-L-SpCas9 and Cas9-L-T4 by changing Asp-219 reside to Ala (T4-D219A) or inactivated the T4 DNA polymerase domain in T4-L-SpCas9 and Cas9-L-T4 by substituting the Asn-214 reside to Ser (T4-D214S)(Abdus Sattar, Lin et al., 1996). HTS results demonstrated that the mutation of T4-D219A, but not T4-D214S, eliminated the increased frequency of deletions at the PAM distal region induced by T4-L-SpCas9 and Cas9-L-T4 relative to Cas9-only (**Appendix Fig S1J-K**), confirming our hypothesis. We next engineered tdTomato MS2-sgRNA by incorporating MS2 stem-loops into the classical sgRNA scaffold region (Konermann, Brigham et al., 2015). We compared the editing profiles at the tdTomato-d151A site produced by co-expression of Cas9 and MCP-tagged T4 DNA polymerase with tdTomato sgRNA to that with tdTomato MS2-sgRNA. We confirmed that the untethered and MS2-tethered T4 DNA polymerase was comparable in favoring 2-bp insertions at the tdTomato-d151A site (**Appendix Fig S1L**), potentially because of the saturated expression of T4 DNA polymerase when delivered as vectors. Subsequent in-vitro experiments all used the untethered T4 DNA polymerase.

We next assessed the impact of T4 DNA polymerase on Cas9 editing on the endogenous loci. We designed 26 guide RNAs (gRNAs) targeting multiple pathogenic genes with different chromosomal locations (TS1–TS26; **Appendix Fig S1M**). Compared to the Cas9-only treatment control (CTR), T4 DNA polymerase selectively increased 1–2-bp insertions over deletions (≥ 1-bp) or increased 1–2-bp deletions over ≥ 3-bp deletions, depending on the gRNA context (**Appendix Fig S1N-Q**); Among the 16 guide RNAs that favored 1-bp insertions with Cas9 and T4 DNA polymerase, 11 (68.8%) contained an adenine (A) or thymine (T) while 5 (31.2%) contained an guanine (G) or cytosine (C) at position -4 (counting the NGG PAM sequences as nucleotides 0–2). Among the ten guide RNAs that biased for 1-bp deletions, six (60%) had an identical nucleotide at positions -4 and -3 while 4 (40%) had a repeating nucleotide at positions -5 and -4. All four guide RNAs that contained a repeating nucleotide at positions -4 and -2 or -5 and -3 led to deletions of the repeating nucleotide and intervening nucleotide (**Appendix Fig S1R**). Collectively, these results reveal that T4 DNA polymerase favors small insertions over deletions, or small deletions (1–2-bp) over ≥ 3-bp deletions depending on the cleavage site sequences. We refer to this novel gene editing platform -- phage T4 DNA polymerase associated with Cas9 -- as CasPlus editing.

### Improved CasPlus editing efficiency via engineered T4 DNA polymerase

We hypothesized that site-directed mutagenesis on the exonuclease or T4 DNA polymerase domains might affect the fill-in step and change the CasPlus editing efficiency (**Fig 2A**). We constructed seven T4 DNA polymerase mutants with a decreased or increased DNA mutation frequency relative to wild-type T4 DNA polymerase (T4-WT), and one N-terminal truncation mutant lacking the exonuclease domain (Δ1-377)(Abdus Sattar et al., 1996, Reha-Krantz, 1988, Reha-Krantz, 1998, Reha-Krantz, Stocki et al., 1991) (**Fig 2B**). Compared to T4-WT, Asp-219 residue to Ala mutant (T4-D219A), had 2.4-fold more 1-bp insertions, at the expense of deletions at TS11 (**Fig 2C**). T4-WT and the T4 DNA polymerase mutants exhibited similar 1-bp insertion patterns at TS11 (**Appendix Fig S2A)**. T4-D219A led to up to 6-fold increases in 1-bp insertions at the expense of deletions across 17 genomic sites tested (**Appendix Fig S2B-D**). Mutations changing the Asp-222 residue to Ala in RB69 DNA polymerase (RB69-D222A) (Hogg et al., 2006) also resulted in a 2-to-4-fold higher frequency of 1-bp insertions at the TS2, TS11, and TS12 than RB69-WT (**Appendix Fig S2E**). Together, these results support that CasPlus editing enhances precise 1-bp insertions, with T4-D219A outperforming T4-WT and RB69-D222A outperforming RB69-WT.

**Figure 2.**
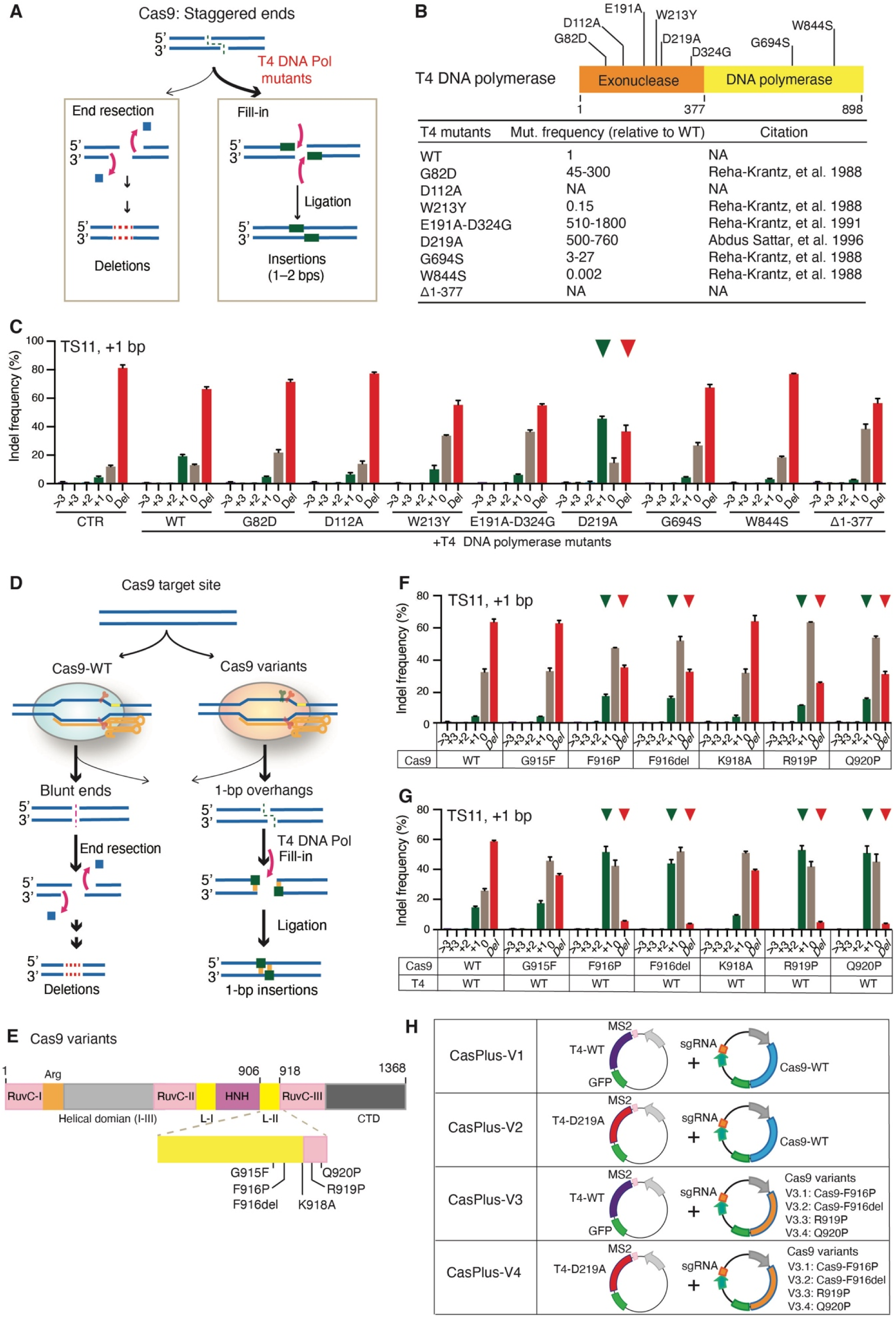
Engineered T4 DNA polymerase mutants and Cas9 variants improve CasPlus editing efficiency. (**A**) Cas9-induced staggered ends can be processed for either end resection that generates deletions, or fill-in that produces insertions. T4 DNA polymerase mutants can enhance the fill-in process, thereby increasing 1–2-bp insertions. (**B**) Schematic showing the amino acid alterations introduced into T4 DNA polymerase; the corresponding relative mutation frequencies are below. NA, non-available. (**C**) Frequency of Cas9-induced indels at TS11 with or without T4 DNA polymerase mutants. (**D**) At target sites where Cas9-WT often generates blunt ends, leading to deletions (left), engineered Cas9 variants can preferentially generate staggered ends with 1-bp overhangs, enabling T4 DNA polymerase to produce 1-bp insertions (right). (**E**) Amino acid alterations introduced within Cas9. (**F**-**G**) Frequency of Cas9-induced indels at TS11 in cells transfected with Cas9-WT or distinct Cas9 variants individually (**F**) or along with T4-WT (**G**). (**H**) Four distinct versions of the T4 DNA polymerase mediated CasPlus system.

We compared CasPlus and prime editing (PE) in generation of 1-bp insertions at position -4 of TS5 and TS10 (**Appendix Fig S2F**). For each target site, we screened four prime editing guide RNAs (pegRNAs) varying in sizes of reverse transcriptase templates (RTT) and prime binding sequences (PBS) and selected the most efficient pegRNA for comparison (**Table EV1**). PE3 resulted in ∼16.1% 1-bp insertions without undesired indels at TS5 and ∼8.2% 1-bp insertions without undesired indels at TS10 while Cas9 combined with T4-D219A induced ∼72.3% 1-bp insertions and ∼4.8% undesired indels at TS5 and ∼48.8% 1-bp insertions and ∼4.4% undesired indels at TS10 (**Appendix Fig S2G**). These results indicated that, at certain defined genome sites, CasPlus editing could be a more efficient approach in introducing precise1-bp insertions than prime editing.

To further investigate the capabilities of T4 DNA polymerase in favoring small insertions with other Cas proteins, we tested Cas12a, an RNA-guided endonuclease that creates 5′ staggered ends with 5–8-bp overhangs(Zetsche, Gootenberg et al., 2015). We reasoned that T4 DNA polymerase could fill in those overhangs, creating 5–8 nucleotides insertions. Because the Cas12a cleaves the target DNA at the distal end of PAM (18-25 bp away), Cas12a can recut the allele until an indel produced upstream of the insertions to prevent recutting (**Appendix Fig S2H**). Thus, we co-expressed T4 DNA polymerase along with Cas12a and guide RNA-Lb1 into cells and analyzed the proportion of alleles containing 5-8 nucleotides insertions. As expected, T4-WT and T4-D219A substantially increased the frequency of alleles carrying 5-8 nucleotides insertions compared to Cas12a-only treatment (**Appendix Fig S2I-J**). Hence, with Cas12a, T4 DNA polymerase was also capable of filling in the 5′ staggered ends with 5–8-bp overhangs, which is consistent with our results in Cas9-mediated editing.

### Combining Cas9 variants with T4 DNA polymerase expands the target range for CasPlus editing

CasPlus editing requires Cas9 to generate DNA ends with 5′ overhangs. Certain Cas9 variants preferentially produce staggered ends with 1-bp overhangs, while wild-type Cas9 (Cas9-WT) mainly generates blunt ends at cleavage sites (Jiang, Taylor et al., 2016, Shou, Li et al., 2018). We hypothesized that combining these Cas9 variants with T4 DNA polymerase would favor 1-bp insertions at target sites (**Fig 2D-E**). At TS11, the Cas9 variants F916P, F916del, R919P, and Q920P produced an average of 15% 1-bp insertions, whereas Cas9-WT produced <5% 1-bp insertions (**Fig 2F**). Combining these Cas9 variants with T4-WT resulted in an average of ∼50% 1-bp insertions and ∼4.5% deletions whereas combining Cas9-WT and T4-WT yielded ∼15% 1-bp insertions and ∼59% deletions (**Fig 2G**). We tested five target sites that rarely induced 1-bp insertions by Cas9-WT with or without T4-WT. Cas9-F916P led to an average 4.3-fold and Cas9-F916del led to an average 5.1-fold increase in 1-bp insertions at the expense of deletions across the five target sites with T4-D219A compared to Cas9-WT or Cas9 variants alone (**Appendix Fig S2K**). Our strategy expanded targetable sites for CasPlus editing for precise insertions. We predicted that Cas9 variants with T4 DNA polymerase could produce longer insertions (2–4-bp) at target sites where Cas9-WT and T4-WT only produce 1–3-bp insertions (**Appendix Fig S2L**). With T4-WT, Cas9 variants F916P, F916del, or Q920P substantially increased 3-bp insertions at tdTomato-d151A site where Cas9-WT produced 2-bp insertions (**Appendix Fig S2M-N**). With T4-WT or T4-D219A, Cas9-F916P and Cas9-F916del promoted 2-bp insertions at TS5, 3-bp insertions at TS17, and 4-bp insertions at TS18 whereas Cas9-WT produced 1-bp insertions at TS5, 2-bp insertions at TS17 and 3-bp insertions at TS18 (**Appendix Fig S2O**).

The four versions of the CasPlus system (**Fig 2H**) generate distinct editing outcomes. At target sites where CasPlus-V1 predominantly favors 1–3-bp insertions, CasPlus-V2 offers even higher efficiency. CasPlus-V3 and -V4 can increase insertion length by 1 bp. At target sites where CasPlus-V1 mainly generates 1- or 2-bp deletions, CasPlus-V2, -V3, and -V4 favor 1-bp insertions (**Appendix Fig S2P**).

### CasPlus enhanced efficiency of *DMD* (exon 52 deletion) correction

CasPlus editing enhances 1–2-bp insertions, which can efficiently correct diverse disease-causing frameshift mutations, such as those found in *DMD* mutations with deleted exons. Precise insertion of 1 bp at the 3′ end of exon 51 or the 5′ end of exon 53 could reframe mutated *DMD* with exon 52 deleted (**Fig 3A**). We generated all gRNAs targeted exon 51 or exon 53 which can potentially reframe *DMD* del exon 52 and determined their editing efficiencies in HEK293T cells (**Fig 3B**). The gRNAs G10 and Ex51-G2 targeting exon 51 and G9 and Ex53-G3 targeting exon 53 had higher editing efficiencies compared to other gRNAs (**Fig 3C**). Consistent with gRNA G10 and G9 (**Appendix Fig S2B**), Ex51-G2 and Ex53-G3, when used with CasPlus-V1 or -V2, resulted in a 1.6-to-2-fold increase in correction efficiency compared to Cas9-only samples, primarily due to the increased 1-bp insertions (**Appendix Fig S3A**). Compared to Cas9, CasPlus editing is more flexible and efficient in correcting *DMD* mutations. Using CasPlus-V1 and -V2, gRNA G10 led to a 4.3-to-6.5-fold increased correction efficiency in exon 51, and gRNA G9 increased correction efficiency by 2.1-to-2.5-fold in exon 53 versus using Cas9-only in a human induced pluripotent stem cell (iPSC) line with DMD exon 52 deletion (DMD-del52). (**Fig 3D and appendix Fig S3B**). We differentiated the pool of edited iPSCs and a single clone (SC) with 1-bp insertion into cardiomyocytes (iCMs). Compared to Cas9 editing, frequencies of in-frame mRNAs (3*n*+1) in iCMs resulting from CasPlus-V1 and -V2 editing were ∼2.5- fold higher in exon 51 and ∼1.5-fold higher in exon 53 (**Fig 3E**). Western blot analysis confirmed that CasPlus-V1 and -V2 editing produced 1.5-to-3-fold higher dystrophin expression than Cas9-only editing (**Fig 3F**).

**Figure 3.**
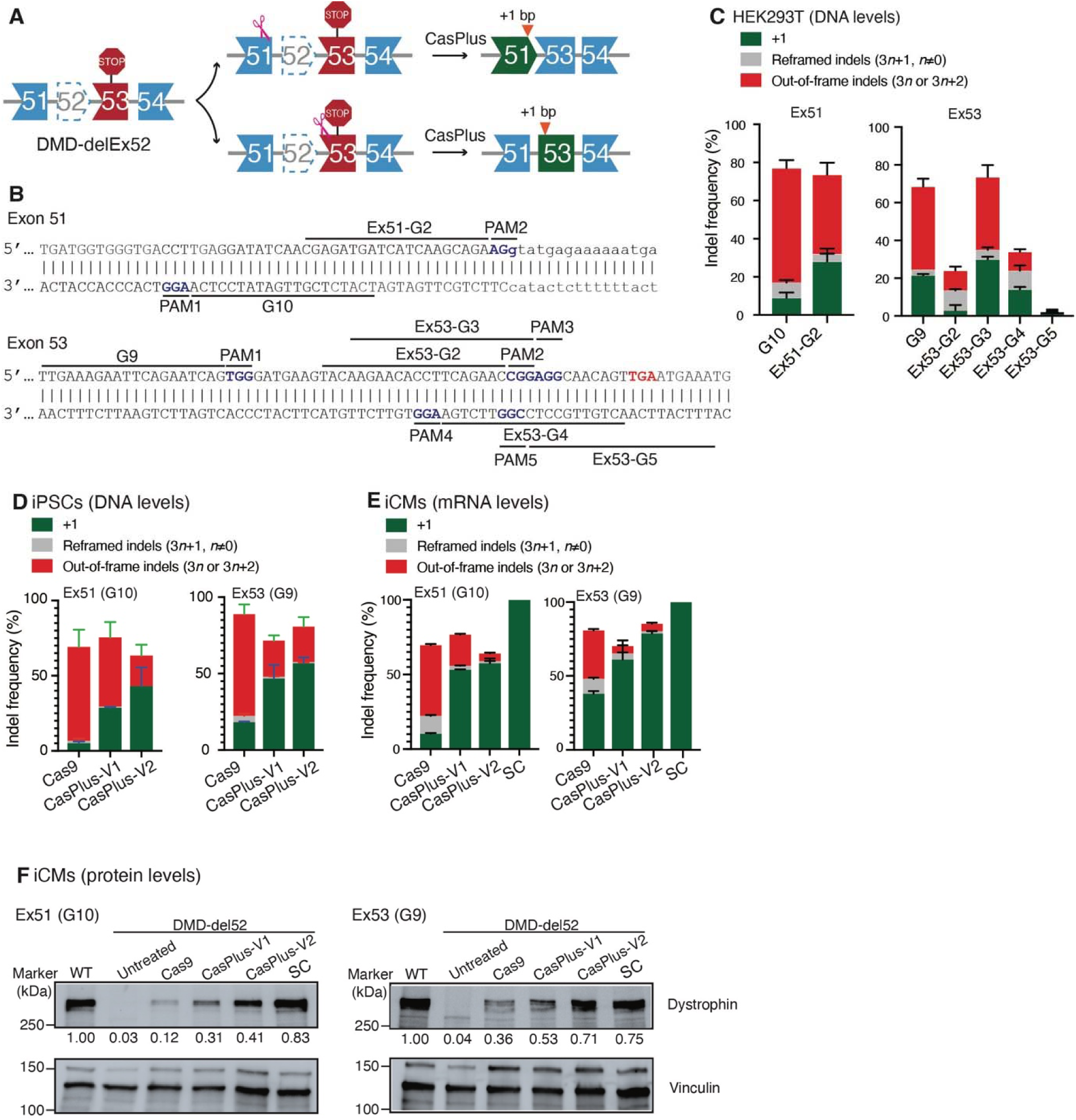
CasPlus enhanced efficiency of *DMD* (exon 52 deletion) correction. (**A**) Deletion of *DMD* exon 52 creates a premature stop codon in exon 53, preventing dystrophin expression. Two CasPlus editing strategies can restore dystrophin expression by generating 1-bp insertions. (**B**) All potential gRNA sequences with an NGG PAM on *DMD* exons 51 and 53 available for restoring the *DMD* exon 52 deletion mutation. Red, premature stop codon. (**C**) Frequencies of in-frame (3*n*+1) and out-of-frame (3*n* or 3*n*+2) indels induced by Cas9 with gRNAs described in **B** in HEK293T cells. Frequencies were calculated based on ICE analysis on Sanger sequencing files. (**D**) Frequencies of in-frame (3*n*+1) and out-of-frame (3*n* or 3*n*+2) indels induced by Cas9, CasPlus-V1, and CasPlus-V2 with gRNA G10 or G9 in DMD-del52 iPSCs. (**E**) Edited DMD-del52 iPSCs described in (**D**) were differentiated into iCMs and then in-frame (3*n*+1) and out-of-frame (3*n* or 3*n*+2) indels on the mRNA levels were analyzed by HTS. (**F**) Western blot analysis of the expression of dystrophin and vinculin in DMD-WT iCMs, untreated DMD-del52 iCMs, DMD-del52 iCMs differentiated from iPSCs with edits at *DMD* exon 51 (left) or 53 (right), or DMD-del52 single clone containing 1-bp insertions. The normalized dystrophin quantitation is shown below the gel image.

To investigate whether co-expression of T4 DNA polymerase and Cas9 increases genome-wide off-target effects, we performed whole-genome sequencing (WGS) in HEK293T cells with or without Cas9 or CasPlus editing using gRNA G10 (**Table EV2**). We observed no apparent differences in insertion/deletion (indel) or single-nucleotide polymorphism (SNP) profiles when comparing CasPlus to Cas9 editing (**Appendix Fig S3C-F**, and **table EV3**). We did not detect gene editing in Cas9- or CasPlus-edited cells at off-target sites predicted by Cas-OFFinder (Bae, Park et al., 2014)(**Appendix Fig S3G-H** and **table EV4**). Phage T4 DNA polymerase is essential for replication of phage genome(De Waard, Paul et al., 1965). To assess the impact of transient co-expression of engineered T4 DNA polymerase and Cas9 on cell cycle progression, we stained the transfected HEK293T cells using propidium iodide (PI) and measured the distribution of cells in three major phases of the cycle (G1, S and G2/M) via flow cytometry. A slight but not significant increase of cells in S phase was observed in cells with CasPlus-V1 but not CasPlus-V2 editing verse Cas9 editing. Collectively, these results indicate that CasPlus potentially did not raise additional safety concerns compared to Cas9 editing (**Appendix Fig S3I**).

### Repression of on-target large deletions by CasPlus editing in iPSCs

Unexpected on-target large deletions can arise from the long-range end resection that occurs during Cas9 editing (Yoo et al., 2022) (**Fig 4A**). In our corrected DMD-del52 iCMs, at the mRNA level, we detected the unexpected skipping of the whole exon 51 using gRNA G10 and exon 53 using gRNA G9 in Cas9-only editing (**Appendix Fig S4A-B**). We speculated that these exons loss resulted from unexpected on-target large deletions which eliminate exon 51 or exon 53. To investigate this, we PCR-amplified a ∼2-kilobase (kb) region in pools of edited cells (**Appendix Fig S4C**). We observed several lower bands representing deletions of ∼0.5 kb distal to gRNA G10 and ∼1.2 kb distal to gRNA G9 in Cas9-only but not CasPlus-edited iPSCs and iCMs (**Fig 4B and appendix Fig S4D-G**). We amplified a ∼5-kb region around the *DMD* exon 51 and 53 target sites from pools of edited iPSCs and sequenced the PCR amplicons using PacBio sequencing technology. Up to 23.0% of the PacBio reads contained deletions of 200-3,000 bp around the exon 51 cleavage site in Cas9-edited cells (**Fig 4C, Appendix Fig S4H**, and **table EV5)**. This on-target effect was not observed in untreated cells or cells edited with CasPlus-V1 or -V2. In untreated cells, we detected ∼3-kb deletions around *DMD* exon 53 in 13.2% of the PacBio reads. This result likely resulted from artifact due to PCR amplification process, as 3-kb deletions of similar scale were observed in all tested cells [Cas9 (11.1%); CasPlus-V1 (9.4%); CasPlus-V2 (14.8%)]. On *DMD* exon 53, reads with deletions of 200-3,500 bp around the cut site were 2.8 to 5.1-fold less frequent in cells edited with CasPlus-V1 (9.5%) and -V2 (17.4%) versus Cas9 (48.9%) (**Fig 4D, appendix Fig S4H, and table EV5**). We genotyped single clones sorted from edited iPSC pools to confirm our results. Three of 33 clones (9.1%) using Cas9 with gRNA G10 and three of 13 clones (23.1%) using Cas9 with gRNA G9 contained deletions of 300 bp to 10 kb. No clones from CasPlus-V1- or CasPlus-V2 edited cells had deletions >150 bp (**Fig 4E and appendix Fig S4I-K**). To investigate whether CasPlus editing could inhibit on-target large deletions on other genomic loci, we used a flow cytometric assay(Kosicki et al., 2022, Kosicki et al., 2018) to detect and isolate cells containing large deletions and more complex rearrangements on the X-linked *PIGA* gene. We transfected male iPSCs with a gRNA targeted the intron 1 and observed a significant decrease in the frequency of PIGA^-^ populations in cells treated with CasPlus verse Cas9 (**Appendix Fig S4L**). Genotyping of single clones from PIGA^-^ populations confirmed that all clones contained large deletions (>265 bp) resulting in partial or entire deletion of exon 2 (**Appendix Fig S4M**). Thus, CasPlus efficiently represses on-target large deletions in mammalian cells.

**Figure 4.**
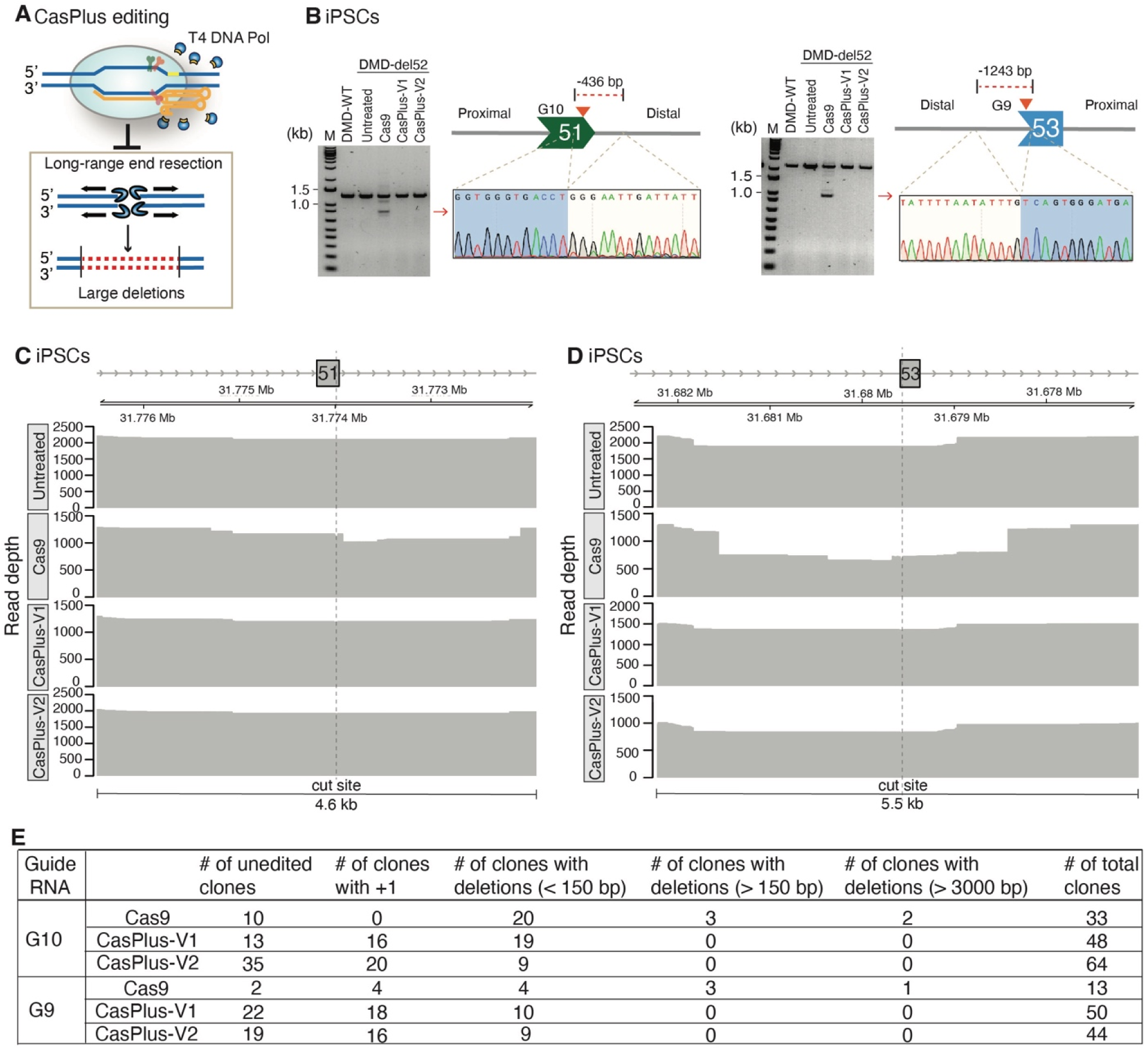
Repression of on-target large deletions by CasPlus editing in iPSCs. (**A**) CasPlus editing represses the generation of large deletions by counteracting the long-term end resection process. (**B**) Representative gel images showing the PCR products amplified from DMD-WT iPSCs, untreated DMD-del52 iPSCs or DMD-del52 iPSCs with edits at *DMD* exon 51 (**left**) or 53 (**right)**. Lower bands (red arrowheads) were purified and sequenced. (**C-D**) Depth of PacBio reads at *DMD* exon 51 (**C**) or 53 (**D**) in untreated, Cas9-, CasPlus-V1-, and CasPlus-V2-edited DMD-del52 iPSCs. (**E**) Table summarizing number of unedited single clones and single clones containing 1-bp insertions or deletions with different sizes. Single clones were isolated from Cas9-, CasPlus-V1-, and CasPlus-V2-edited DMD-del52 iPSCs.

### Repression of on-target large deletions by CasPlus editing in mouse germ line

On-targe large deletions frequently occur during CRISPR/Cas9 mediated mouse embryo-editing(Adikusuma, Piltz et al., 2018, Papathanasiou, Markoulaki et al., 2021). To evaluate the capacity of CasPlus editing in inhibiting on-target large deletions for mouse embryo editing, we microinjected Cas9 mRNA and gRNA against *Mybpc3* alone (Cas9) or combined with T4-WT (CasPlus-V1) or T4-D219A (CasPlus-V2) mRNA into fertilized mouse zygotes. Four days later, we harvest the embryo for future analysis (**Fig 5A**). The editing efficiency of each embryo varied between 80%-100% in both Cas9- and CasPlus-edited group (**Appendix Fig S5A**). To assess the presence of large deletions, we performed multiple PCRs to amplify the proximal, distant, or both side of the PAM. Full-length and lower molecular mass PCR bands were analyzed by Sanger sequencing (**Fig 5B and appendix Fig. S5B**). The sequencing results showed that 8/10 embryos (80%) treated with Cas9, and 3/10 embryos (30%) treated with CasPlus-V1 or V2 harbored > 500 bp large deletions (**Fig 5C-D and appendix Fig. S5C-D**). Collectively, these results indicated that CasPlus-V1- and -V2 editing efficiently repressed Cas9 mediated on-target large deletions in mouse germline editing.

**Figure 5.**
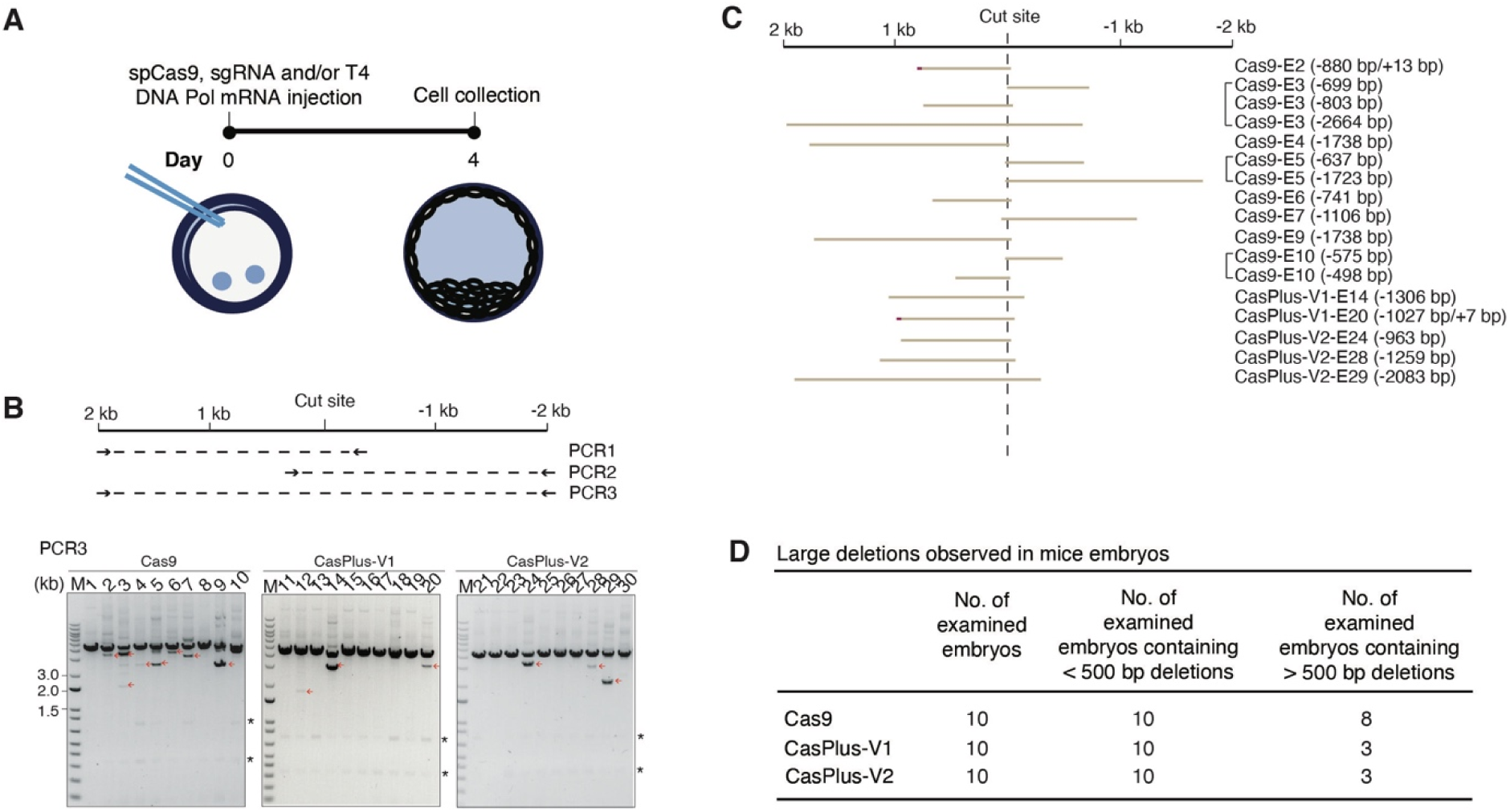
Repression of on-target large deletions by CasPlus editing in mouse germline. (**A**) At day 0, Fertilized mouse zygotes were microinjected with Cas9 mRNA and a sgRNA alone or combined with T4 DNA polymerase mRNA. At day 4, all embryos were collected for analysis. (**B**) Top schematic illustrating the sizes and locations of the three PCR amplicons used to detect large deletions. Gel images showing the PCR result for PCR3. Small molecular mass products (red arrowheads) were sequenced. Asterisk, non-specific bands. (**C**) The sequencing results for the small molecular mass products observed in **B**. (**D**) Table summarizing the total number of embryos tested, the number of tested embryos containing small deletions (<500 bp) and the number of tested embryos containing large deletions (>500 bp) in Cas9, CasPlus-V1- or -V2 edited group.

### Repression of on-target chromosomal translocations during multiplex gene editing by CasPlus editing in primary T cells

Chromosomal translocations can occur when two simultaneous DSBs are present on two chromosomes during Cas9 mediated single gene editing or multiple gene editing(Amit, Iancu et al., 2021). To investigate whether CasPlus editing can reduce chromosomal translocations, we recapitulated translocation events between the genes *CD74* (chromosome 5) and *ROS1* (chromosome 6) in HEK293T cells (Choi & Meyerson, 2014) (**Appendix Fig S6A)**. We PCR-amplified the breakpoint junction regions on the fused chromosomes and assessed translocation efficiencies. The translocation frequencies between *CD74* and *ROS1* were ∼5-fold lower with CasPlus-V1 and ∼2-fold lower with CasPlus-V2 versus Cas9 editing (**Appendix Fig S6B-C**). The frequencies of insertions at *ROS1* and *CD74* individual sites were higher with CasPlus-V1 and -V2 editing compared to Cas9 (**Appendix Fig S6D**). We observed similar trends of repression of chromosomal translocations in iPSCs (**Appendix Fig S6E-G**). Genotyping revealed that three of 95 clones (3.2%) from the Cas9 edited iPSC pool but no clones from the CasPlus edited iPSC pools contained *ROS1-CD74* and *CD74-ROS1* translocations (**Appendix Fig S6H-K**).

In a clinical trial (NCT03399448), chromosomal translocations were frequently observed between targeted chromosomes in Cas9-engineered T cells when delivered with three gRNAs targeting *PDCD1*, *TRBC1/2* and *TRAC* genes on different chromosomes (Stadtmauer et al., 2020). We compared the same three gRNAs with Cas9 or CasPlus to assess translocations in these three chromosomes in HEK293T cells (**Fig 6A and appendix Fig S6L**). PCR demonstrated that CasPlus-V1 led to a 2.5-to-4.5-fold decrease in all translocation types in the three chromosomes (**Fig 6B-C and appendix Fig S6M-N**). TaqMan assays revealed that CasPlus-V1 decreased an average 5.8-fold, and CasPlus-V2 decreased an average 3.9-fold translocations compared to Cas9 editing (**Fig 6D**). CasPlus-V1 editing induced similar gene disruption efficiency to Cas9 editing for four genes (**Fig 6E**). CasPlus-V2 induced a similar gene disruption efficiency but was less effective in repressing chromosomal translocations compared to CasPlus-V1. We recommended CasPlus-V1 to generate CAR-T cells.

**Figure 6.**
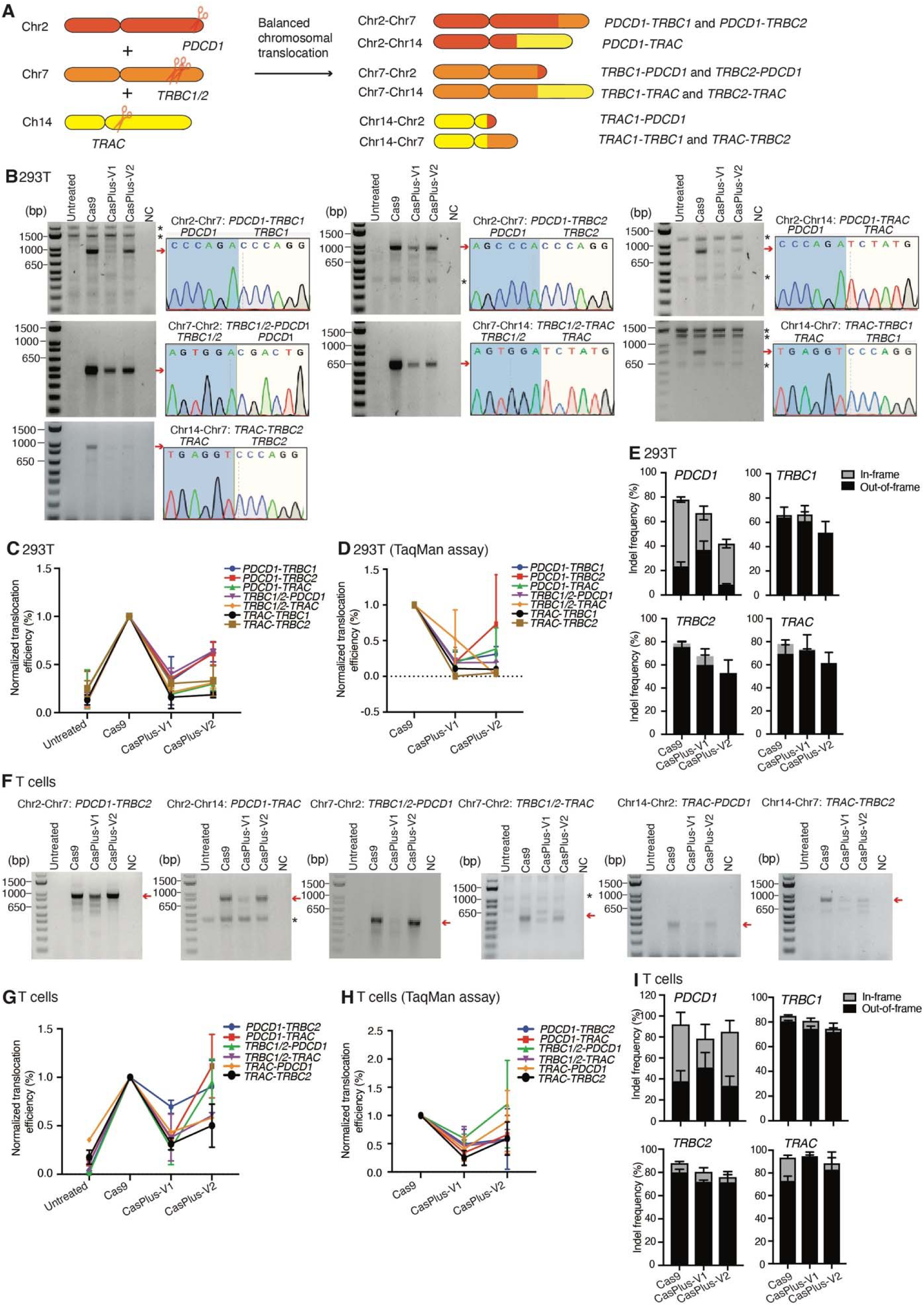
Repression of on-target chromosomal translocations during multiplex gene editing by CasPlus editing in primary T cells. (**A**) Schematic illustrating the balanced translocations among genes *PDCD1*, *TRBC1/2*, and *TRAC*. (**B**) Representative gel images demonstrating the balanced translocations detected in HEK293T cells during Cas9, CasPlus-V1, or CasPlus-V2 editing (see methods). Balanced translocation of Chr14-Chr2: *TRAC*-*PDCD1* was undetectable by PCR. Bands with expected size (red arrowheads) were purified, TA-cloned and sequenced (one of the TA clone results was show for each type of translocation). Asterisk, non-specific bands. (**C**) Normalized quantification of data in **B**. (**D**) Evaluation of balanced chromosomal translocations in HEK293T cells during Cas9, CasPlus-V1, or CasPlus-V2 editing by TaqMan assay (see methods). For each assay, the translocation efficiency in Cas9 edited cells is set as 1. (**E**) Frequencies of out-of-frame and in-frame indels at four individual sites in HEK293T cells during Cas9, CasPlus-V1, or CasPlus-V2 editing. Frequencies were calculated based on ICE analysis on Sanger sequencing files. (**F**) Representative gel images demonstrating the balanced translocations detected in primary human T cells during Cas9, CasPlus-V1, or CasPlus-V2 editing (see methods). Balanced translocation of Chr14-Chr7: *TRAC*-*TRBC1 and* Chr2-Chr7: *PDCD1*-*TRBC1* was undetectable by PCR. (**G)** Normalized quantification of data in **F.** Values and error bars reflect mean ± SEM of n=3 replicates from two donors. (**H**) Evaluation of balanced chromosomal translocations in primary human T cells during Cas9, CasPlus-V1, or CasPlus-V2 editing by TaqMan assay (see methods). Values and error bars reflect mean ± SEM of n=3 replicates from two donors. (**I**) Frequencies of out-of-frame and in-frame indels at four individual sites in primary human T cells during Cas9, CasPlus-V1, or CasPlus-V2 editing. Frequencies were calculated based on ICE analysis on Sanger sequencing files. Values and error bars reflect mean ± SEM of n=3 replicates from two donors.

We investigated CasPlus editing efficacy in suppressing chromosomal translocations among genes *PDCD1*, *TRBC1/2* and *TRAC* in primary human T cells. First, we nucleofected the activated primary human T cells with Cas9 mRNA and *PDCD1*, *TRBC1/2* and *TRAC* sgRNAs alone (Cas9) or in conjunction with T4-WT (CasPlus-V1) or T4-D219A (CasPlus-V2) mRNA. Four days after mRNA introduction, we accessed translocations using PCR and TaqMan assay. CasPlus-V1 induced 1.5-to-11-fold fewer translocations than Cas9 editing in PCR assays (**Fig 6F-G**). TaqMan assays confirmed that CasPlus-V1 reduced translocations by 1.2-to-4.3-fold relative to Cas9 (**Fig 6H**). As in HEK293T cells, CasPlus-V2 also suppressed translocations but was less efficient compared to CasPlus-V1. Analysis of the on-target gene editing through sequencing revealed that CasPlus editing had similar knockout efficiency at *PDCD1, TRBC1/2*, and *TRAC* sites in T cells **(Fig 6I).** Together, our findings indicate that CasPlus editing significantly inhibits Cas9-mediated on-target chromosomal translocations and offers a potentially safer editing strategy to engineer T cell and other ex vivo cell therapies.

## Discussion

Despite the successes achieved with all CRISPR/Cas-based genome editing technologies, including base editing and prime editing, the clinical potential of the technologies is currently constrained by the safety and efficacy concerns in cultured cells and animal models, and likely in humans. Unexpected on-target large deletions and chromosomal translocations from Cas9 editing are an emerging risk factors for *in vitro*, *in vivo* and *ex vivo* applications(Kosicki et al., 2018, Leibowitz et al., 2021, Nahmad, Reuveni et al., 2022b, Stadtmauer et al., 2020). Our results show that in cultured human cells, the CasPlus editing platform can substantially reduce deleterious on-target large deletions and chromosomal translocations while maintaining equal or higher efficiency of desired edits (**Appendix Fig S6O**). Future research will assess the safety and efficacy of CasPlus in animal models using adeno-associated virus and lipid nanoparticles delivery systems.

CasPlus utilizes T4 DNA polymerase to fill in the staggered end created by Cas9 or Cas9 variants, producing small insertions and reducing relatively large deletions. Fusion of DNA polymerase I or KF to Cas9 could increase the frequency of 1-bp deletions over >1 deletions by counteracting the DNA resection process(Yoo et al., 2022). We did not observe similar results when co-expressing DNA polymerase I or KF with Cas9 at the tdTomato-d151A site which may reflect a sequence specific effect. Fusion of T4 DNA polymerase to Cas9 strongly favored relatively large deletions distal to the PAM region. Although the mechanism underlying the different effect of fused T4 DNA polymerase, DNA polymerase I or KF with Cas9 are unclear, it may partly be attributed to the higher 3′-to-5′ exonuclease activity of T4 DNA polymerase than DNA polymerase I or KF(Kucera & Nichols, 2008), which may degrade the released 3’ single-strand DNA (ssDNA) when fused to Cas9.

Our data shows that CasPlus can favor templated 1–2-bp insertions as well as 1–2-bp deletions in a guide RNA context dependent manner, but more comprehensive and systematic analysis is needed to predict or better control the outcomes of CasPlus editing. CasPlus-mediated 1–2-bp insertions could be harnessed to disrupt pathogenic genes or correct frameshift mutations. Base editing and prime editing are versatile approaches to disrupt and correct genes without requiring exogenous templates and DSBs. Base editing can generate single-nucleotide mutations but not indels (Gaudelli et al., 2017, Komor et al., 2016). Prime editing can achieve single-nucleotide and indels editing(Anzalone et al., 2019). However, the efficiency of prime editing varies among different pegRNA constructs in diverse targets in cells and organoids and was averagely less than Cas9 when targeting the identical genomic sites(Geurts, de Poel et al., 2021, Kim et al., 2021). Efficient prime editing can achieved by introducing a second nick on the non-edited stand, but can generate undesired DSBs and large deletions in mouse embryos(Tomomi Aida, 2020). Other methods can reduce translocations, including nuclease combinations(Bothmer et al., 2020) or Cas12f nuclease(Xin et al., 2022) for multiplex gene editing and fusion of Cas9 to an exonuclease to prevent repeat cleavage(Yin et al., 2022). However, unlike CasPlus, these approaches cannot precisely control the editing outcomes. Since CasPlus does not negatively affect the editing efficiency of regular Cas9 and can produces a dominant genotype of 1–2-bp indel with much fewer byproducts and on-target damage, it could be a safer and more efficient method optimized for human applications.

Additionally, CasPlus’ enhanced safety features can expand ex vivo applications, including chimeric antigen receptor T (CAR-T) cells to target cancer or autoimmune diseases. The US Food and Drug Administration placed temporary holds on CRISPR/Cas9-based clinical trials related to allogeneic CAR-T therapies following the detection of a chromosomal abnormality in a patient and requested control data on genomic rearrangement(Sheridan, 2022a). As eukaryotic DNA repair mechanisms are conserved, our data from HEK293T, iPSC, and primary T cells suggest that CasPlus can generate engineered cells with fewer chromosomal abnormalities for cell therapy applications. Hence, CasPlus may improve the safety and efficacy of gene editing to develop novel cellular therapies.

## Material and methods

### Plasmids

The vector pSpCas9(BB)-2A-GFP (PX458) (Addgene plasmid #48138) containing the human-codon-optimized SpCas9 gene with 2A-GFP and the sgRNA backbone, vector pCMV-PE2 (Addgene plasmid # 132775), vector pCMV-BE3 (Addgene plasmids # 73021) were purchased from Addgene. p3xFlag-CMV-10 was a gift from Dr. Xiaodong Wang. pLentiV-SgRNA-tdTomato-P2A-BlasR was a gift from Dr. Lukas Dow. EF1A-CasRx-2A-EGFP was a gift from Dr. Patrick Hsu. Plasmids psPAX2 and pMD2G were gifts from Dr. Lei Bu. pBSU6_FE_Scaffold_rsv_GFP was a gift from Dr. Dirk Grimm. To construct the lentiviral vector expressing tdTomato-d151A, the tdTomato-d151A gene was synthesized by Integrated DNA Technologies (IDT). First, it was cloned into vector p3xFlag-CMV-10, then the CMV-10-tdtomato-d151A was cloned into pLentiv-SgRNA-tdTomato-P2A-BlasR using MluI and BamHI restriction sites. For DNA polymerase cloning, the coding sequences of DNA polymerase 4, DNA polymerase I, Klenow fragment, T4 DNA polymerase, RB69 DNA polymerase, and T7 DNA polymerase were codon-optimized for human cell expression using the Genewiz Codon Optimization tool. For each DNA polymerase, an expression cassette containing the polymerase, a MCP (MS2 bacteriophage coat protein) and a hemagglutinin (HA) tag, two copies of a nuclear localization sequence (NLS), and a flexible linker (L) was synthesized from Genewiz and cloned into EF1A-CasRx-2A-EGFP via Gibson assembly. Mutations of T4 DNA polymerase and RB69 DNA polymerase were introduced into the vectors EF1A-MCP-T4-DNA-Polymerase-2A-EGFP and EF1A-MCP-RB69-DNA-polymerase-2A-EGFP, respectively, via Gibson assembly. To construct Cas9 and T4 DNA polymerase fusion protein, T4 DNA polymerase was amplified from EF1A-MCP-T4-Pol-2A-GFP and cloned into backbone pSpCas9(BB)-2A-GFP (PX458) via Gibson assembly. Mutations of Cas9 were generated in the backbone pSpCas9(BB)-2A-GFP (PX458) via Gibson assembly. To test CasPlus editing, Guide RNAs were cloned into pSpCas9(BB)-2A-GFP (PX458) or Cas9 variants derived from pSpCas9(BB)-2A-GFP (PX458) according to the CRIPSR plasmid instructions from the Feng Zhang Lab(Ran, Hsu et al., 2013). Prime editing guide RNAs and guide RNAs for creating a second nick were cloned into pBSU6_FE_Scaffold_rsv_GFP via Gibson assembly or restriction digestion and ligation. All guide RNA sequences were listed in **table EV7**. All DNA sequences synthesized for vector constructions were listed in **table EV8**.

#### Generation of HEK293T cell lines containing the tdTomato-d151A reporter gene

To generate a stable tdTomato-d151A reporter cell line in HEK293T (ATCC, CRL-3216) cells, we co-transfected pLentiV vector expressing tdTomato-d151A, the lentiviral helper plasmids psPAX2 and pMD2G, and pEGFP (Lonza) into HEK293T cells. Single cells expressing GFP were isolated in 96-well plates 72 hours post-transfection and genotyped 2 weeks later. Positive clones were then stored and expanded for subsequent experiments. Primers for genotyping were shown in **table EV6**.

#### Generation of male iPS cell lines containing the *DMD* exon 52 deletion

Male wild type iPSCs (NCRM-1) were electroporated with vectors expressing Cas9, GFP, and a pair of guide RNAs specific for the deletion (DMD-Ex52-g1 and DMD-Ex52-g2, see **table EV7**). Single cells expressing GFP were isolated in 96-well plates 72 hours post-transfection and genotyped 2 weeks later. Positive clones containing the *DMD* exon 52 deletion were stored and expanded for subsequent experiments. Primers for genotyping were shown in **table EV6**.

#### Transfection and sorting of HEK293T cells

HEK293T cells were transfected using Lipofectamine 2000 Transfection Reagent (ThermoFisher Scientific) according to the manufacturer’s instructions. Cell sorting was performed by the Cytometry & Cell Sorting Laboratory Core Facility at New York University Langone Health. Briefly, HEK293T cells were seeded into 12-well plates. After 20–24 hours, cells were co-transfected with 1 μg vectors expressing Cas9 (with 2A-GFP) and a sgRNA, and 1 μg vectors expressing one of the DNA polymerases. Seventy-two hours post-transfection, cells were dissociated using a trypsin-EDTA solution (Corning) for 2 min at 37°C. Subsequently, 2 ml of warm Dulbecco’s modified Eagle’s medium (DMEM) (Corning) supplemented with 10% fetal bovine serum (FBS) (Gemini Bio-Products) was added. The resuspended cells were transferred into a 15-ml Falcon tube and centrifuged at 1000 rpm for 5 min at room temperature. The medium was then removed, and the cells were resuspended in 0.4–1 ml DMEM. Cells were filtered through the 50-μm-mesh cap of a CellTrix strainer (Sysmex). Cells expressing GFP were sorted by flow cytometry into a 5-ml polypropylene round-bottom tube (Corning) for immediate DNA extraction.

#### Isolation of raw DNA from sorted cells

Protease K (20 mg/ml) (Qiagen) was added to DirectPCR Lysis Reagent (Viagen Biotech Inc.) to a final concentration of 50 μg/ml. Sorted cells (4 × 10^4^–1 × 10^5^) were centrifuged at 4°C at 12000 rpm for 5 min and the supernatant was discarded. Cell pellets were resuspended in 20–50 μl of DirectPCR/protease K solution, incubated at 55°C for >2 hours or until no clumps were observed, incubated at 85°C for 30 min, and then spin down briefly (10 sec). 1–2 μl DNA was used for PCR amplification. All PCR primer sequences were summarized in **table EV6**.

#### Prime editing

HEK293T cells were co-transfected with 1 μg vectors expressing PE2, 1 ug vectors expressing pegRNAs, and 0.5μg vectors expressing a second nicking sgRNA. Cells expressing GFP were sorted, and DNA was extracted from sorted cells as described above. PegRNAs and sgRNAs for prime editing were listed in **table EV1**.

#### Human iPSC maintenance and nucleofection

Human iPSC lines were cultured in Stemflex^TM^ medium (ThermoFisher) and passaged approximately every 3 days (1:8–1:12 split ratio). One hour before nucleofection, iPSCs were treated with 10 μM ROCK inhibitor (Y-27632) and dissociated into single cells using Accutase (Innovative Cell Technologies Inc.). Cells (8 × 10^5^) were mixed with 2 μg of a vector expressing Cas9 (with 2A-GFP), and a guide RNA, as well as 2 μg of a vector encoding a DNA polymerase. This mixture was electroporated into cells using the P3 Primary Cell 4D-Nucleofector X kit (Lonza) according to the manufacturer’s protocol. After nucleofection, iPSCs were cultured in StemFlex^TM^ medium supplemented with CloneR (10×) (STEMCELL Technologies) and antibiotic-antimycotic (100×) (ThermoFisher). Three days after nucleofection, cells expressing GFP were sorted as described above and replated in StemFlex^TM^ medium. Ten to fifteen days after sorting, cells were harvested for DNA isolation.

#### Cardiomyocyte differentiation and purification

Human iPSCs (edited iPSC pools or single clones with 1-bp insertions) were induced for differentiation into cardiomyocytes according to the manufacturer’s instructions using the PSC Cardiomyocyte Differentiation Kit (ThermoFisher Scientific). At 15–20 days after differentiation initiation, cells were purified in RPMI-1640 medium lacking glucose supplemented with B27 (ThermoFisher Scientific). Cells were cultured in this medium for 2–4 days. Cardiomyocytes were used for experiments on day 30–40 after the initiation of differentiation.

#### RNA extraction and cDNA synthesis

RNA from iPSC-derived cardiomyocytes was extracted using TRIzol (ThermoFisher Scientific) according to the manufacturer’s protocol. cDNA was synthesized using the Superscript III First-Strand cDNA Synthesis Kit (ThermoFisher Scientific) according to the manufacturer’s instructions. All RT-PCR primer sequences were summarized in **table EV6**.

#### Cell cycle analysis

HEK293T cells were transfected with Cas9 (T2A-GFP) and gRNA G10, or Cas9 (T2A-GFP), gRNA G10 and T4-WT or T4-D219A. Cells were harvested 24 hours post-transfection, washed in PBS, and fixed using cold 70% ethanal/PBS overnight at 4°C. Cells were washed with PBS and then incubated with PBS containing RNase A (1 mg/ml) at room temperature for one hour. Cells were centrifuged and resuspended in PBS containing RNase A (1 mg/ml) and PI (0.5 μg/ml) at room temperature for 15 min. Following PI staining, cells were transferred to 5-ml polypropylene round-bottom tube for cell cycle analysis in BioRad ZE5 analyzer.

#### Western blotting

HEK293T cells and cardiomyocytes (iCMs) differentiated from iPSCs were harvested, centrifuged, and lysed with RIPA lysis buffer (Santa Cruz Biotechnology) according to the manufacturer’s protocol. Samples were lysed and centrifuged, and the supernatant was incubated at 95°C for 10 minutes in the presence of Laemmli sample buffer (Bio-Rad). Proteins (20 μg per sample) were separated on Mini-PROTEAN TGX 4–15% precast SDS-PAGE gels (Bio-Rad) for 1–3 hours at 100 V and then transferred to PVDF membrane (Bio-Rad) at 40 V for 2–8 hours. Membranes were blocked with 5% non-fat milk (Santa Cruz) for 1–2 hours. Membranes were probed overnight at 4°C with anti-HA antibody (MBL, M180-3) and anti-glyceraldehyde-3-phosphate dehydrogenase antibody (Sigma-Aldrich, G8795) or with anti-dystrophin (Sigma-Aldrich, D8168) and anti-vinculin antibody (Sigma-Aldrich, V9131). Membranes were then washed, probed with a goat anti-mouse or goat anti-rabbit IgG H+ L-HRP conjugated secondary antibody (1:5000-1:10000) (Bio-Rad) for 1 hour, and visualized with Luminol reagent (Santa Cruz) according to the manufacturer’s protocol.

#### Detection of large deletions in iPSCs

DMD-del52 iPSCs were co-electroporated with 2 μg vectors expressing Cas9 (with 2A-GFP), and G10 or G9 and either 2 μg empty vectors or vectors expressing T4-WT or T4-D219A. Cells expressing GFP were then sorted into 5-ml polypropylene round-bottom tubes 72 hours post-electroporation. Bulk sorted cells were replaced and expanded. DNA was isolated from expanded bulk cells using the DNeasy Blood and Tissue Kit (Qiagen) 2 weeks later and subjected to large deletions detection (PCR and PacBio sequencing). Single cells were isolated from bulk edited cells into 96-well plates 2 weeks after electroporation and genotyped 2 weeks after isolation. Single cells containing one insert of G at *DMD* exon 51 or T at *DMD* exon 53 were stored and expanded for subsequent experiments. Bulk edited iPSCs and the single clones containing 1-bp insertions were further differentiated into iCMs. DNA was isolated from iCMs and subjected to large deletions detection. Primers for large deletions detection were summarized in **table EV6**.

#### FLAER staining

Nucleofection in iPSCs with a gRNA against gene *PIGA* was performed as described above. Seventy-two hours after nucleofection, cells expressing GFP were sorted and expanded. FLAER staining was performed one week after expansion. Briefly, cells (3 x 10^5^) were harvested and resuspended in 0.1% Bovine serum albumin (BSA) in PBS containing 1 μg/ml Cederlane FLAER (Alexa 488 proaerolysin variant) (Fisher scientific). Cells were cultured in a rotator for 30 min at room temperature and then washed twice with 0.1% BSA in PBS for 2 minutes. Cells were resuspended with 0.1% BSA in PBS and analyzed in BioRad ZE5 analyzer.

### In vitro transcription of T4 DNA polymerase

The Cas9 mRNA (5meC, Ψ) was purchased from TriLink Biotechnologies (L-6125) and single RNA against genes *PDCD1*, *TRAC*, *TRBC* or *Mybpc3* was synthesized from Synthego (https://ice.synthego.com/#/). To construct the vector for in vitro transcription of T4 DNA polymerase, the T4 DNA polymerase expression cassete (without EGFP) was amplified from EF1A-MCP-T4-DNA-Polymerase-2A-EGFP and cloned into pCMV-BE3 via Gibson assembly. The construct pCMV-T7-T4-DNA-polymerase were then used as template for in vitro transcription using Invitrogen™ mMESSAGE mMACHINE™ T7 ULTRA Transcription Kit (ThermoFisher Scientific). The RNA transcripts were purified by MEGAclear Transcription Clean Up Kit (ThermoFisher Scientific) and eluted with nuclease-free water (Ambion). The concentration of T4 DNA polymerase was measured by a NanoDrop instrument (Thermo Scientific).

#### In vitro fertilization and microinjection

All animal procedures were approved by the Institutional Animal Care and Use Committee at the NYU Langone medical center. hDMDdel52/*mdx* female mice (Gifts from Dr. Nicole Datson) at 4 weeks of age were used as oocyte donors for superovulation, which was performed by intraperitoneal injection of PMSG (5 IU, Sigma-aldrich) and hCG hormone (5 IU, Sigma-Aldrich). hDMDdel52/*mdx* male mice (Gifts from Dr. Nicole Datson) at 6-8 weeks of age were used as sperm donors. Four hours after in vitro fertilization, total volume of 10 μl solutions containing complexes of Cas9 mRNA (100 ng/μl) and Mybpc3 sgRNA (50 ng/μl) alone or together with T4 mRNA (50 ng/μl) diluted with DEPC-treated injection buffer (0.25 mM EDTA, 10 mM Tris, pH7.4) were injected into cytoplasm of the zygotes. After microinjection, embryos were cultured in microdrops of KSOM+AA containing D-glucose and phenol red (Millipore) under mineral oil at 37°C for 4 days in a humidified atmosphere consisting of 5% CO^2^ in air.

#### Detection of chromosomal translocations by PCR in HEK293T cells

HEK293T cells were co-transfected with a vector expressing Cas9 (with 2A-GFP), and guide RNAs targeting either genes *ROS1* and *CD74* or genes *PDCD1*, *TRBC1*/*TRBC2,* and *TRAC* individually or along with an empty vector or vector expressing T4-WT or T4-D219A. Transfected cells expressing GFP were sorted into 5-ml polypropylene round-bottom tubes 72 hours post-transfection and sorted cells (1 × 10^6^) were immediately subjected to DNA extraction. DNA was extracted using the DNeasy Blood and Tissue Kit (Qiagen) and 1 μl (50 ng/μl) DNA was used for each PCR reaction. Chromosomal translocations were detected by PCR using a GoTaq kit with primers specifically recognizing the breakpoint junction region of each fused chromosomes. All the guide RNAs used for translocations detection were summarized in **table EV7**. All the primers used for PCR were summarized at **table EV6**.

#### Detection of chromosomal translocations by TaqMan qPCR assay

A panel of TaqMan qPCR assays were performed to detect the balanced chromosomal translocations among genes *PDCD1*, *TRBC1*/*TRBC2,* and *TRAC* in HEK293T cells and primary T cells. To increase the sensitivity and accuracy of TaqMan assays in HEK293T cells, purified genomic DNA (25 ng) was first amplified according to the manufacturer’s instructions using REPLI-g Single Cell kit (Qiagen). Amplified genomic DNA was then diluted into ∼800 ng/μl and 1 μl DNA was used for each qPCR reaction. In primary T cells, DNA was extracted from unedited or edited cells, diluted into ∼10 ng/μl and 2 μl DNA was used for each qPCR reaction. For each assay, a separate 20x FAM-labelled master mix was prepared with 1.8 μl forward primer (100 μM), 1.8 μl reverse primer (100 μM), 0.5 μl FAM labelled probes (100 μM), and 8.1 μl H_2_O. A qPCR reaction was prepared with 10 μl 2x TaqMan gene expression master mix (Applied Biosystems), 1 μl 20x FAM-labelled master mix, 1 μl amplified DNA (∼800 ng/μl) and 8 μl H_2_O. Assays were run in StepOne Real-time PCR systems (Applied Biosystems) using the following program: 2 min at 50°C, 10 min at 95°C and followed by 40 cycles of 20 sec at 95°C, 1 min at 60°C. The primers and probes used for TaqMan assays were listed at **table EV6**.

#### Primary human T cell isolation and stimulation

Human peripheral blood mononuclear cells (PBMCs) were purchased from Lonza. Frozen PBMCs were thawed and cultured in X-vivo 15 medium (Lonza) with 5% human AB serum (Heat inactivated) (Valley Biomedical) for one day. Primary T cells were isolated from PBMCs using EasySep Human T Cells Isolation Kits (StemCell Technologies) according to the manufacturer’s instructions. Immediately after isolation, primary T cells were activated and stimulated with a 1:1 ratio of anti-human CD3/CD28 magnetic Dynabeads (Thermo Fisher) to cells in X-vivo 15 medium supplemented with 5% human AB serum (Heat inactivated), 5 ng/mL IL-7 (PeproTech), 5 ng/mL IL-15 (PeproTech), and 200 U/mL IL-2 (PeproTech). Two days after stimulation, magnetic beads were removed, and T cells were ready for nucleofection.

#### T cell nucleofection

Nucleofection in T cells were performed using P3 Primary Cell 4D-Nucleofector™ X Kit S (Lonza) according to the manufacturer’s instruction with minor modifications. Briefly, T cells (∼5-7 x 10^5^) were collected and resuspended in 20 μl P3 buffer. 1μg Cas9 mRNA and 3 μg sgRNAs (*TRAC*, *TRBC*, *PDCD1* sgRNA, 1 μg each) alone or together with 2 μg IVT T4 mRNA were added to the cells before nucleofection in a Lonza 4D-Nucleofector with pulse code EO-115. 80 μl X-vivo 15 medium with 5% human AB serum and 200 U/mL IL-2 was added to the nucleofected cells before a 15 min recovery at 37°C. Nucleofected T cells were plated at a density of ∼0.5-1x 10^6^ cells/ml in X-vivo 15 medium supplemented with 5% human AB serum and 200 U/mL IL-2 in 48-well plates and replenished as needed to maintain a density of 10^6^ cells per ml. Four days after nucleofection, edited cells were harvested for analysis.

#### PCR amplicon preparation for high-throughput sequencing

To prepare for high-throughput sequencing, PCR amplicons of ∼300 bp were amplified using a GoTaq kit (Promega), separated on a 2% agarose gel, and purified with the MinElute Gel Extraction Kit (Qiagen). For each sample, gel-purified PCR product was barcoded with the Nextera Flex Prep HT kit according to the manufacturer’s instructions and sequenced using the MiSeq paired-end 150-cycle format by the Genome Technology Center Core Facility at New York University Langone Health.

#### PCR amplicon preparation for whole-genome sequencing

To prepare for whole-genome sequencing (WGS), HEK293T cells were co-transfected with a vector expressing only Cas9, T4-WT, or T4-D219A proteins; Cas9 in combination with G10; or Cas9 and either T4-WT or T4-D219A in combination with G10. All the vectors transfected expressed GFP. Seventy-two hours post-transfection, cells expressing GFP were sorted into 5-ml polypropylene round-bottom tubes for immediate DNA isolation. Genomic DNA was extracted using the DNeasy Blood and Tissue Kit (Qiagen), barcoded with the PCR-free library prep kit according to the manufacturer’s instructions, and sequenced using a S2 300-cycle flow cell v1.5 by the Genome Technology Center Core Facility at New York University Langone Health.

#### PCR amplicon preparation for PacBio sequencing

To prepare samples for PacBio sequencing, genomic DNA was extracted from iPSCs using the DNeasy Blood and Tissue Kit. Barcodes were added to the target region via a two-step PCR reaction. The first-round PCR was performed using LA Taq DNA polymerase (Takara) according to the manufacturer’s instructions. The first-round PCR amplified a 5-kb region around the target site using target-specific primers tailed with universal forward and reverse sequences. The second round of PCR re-amplified and barcoded the first round of PCR products using universal, barcoded forward and reverse primers. The final barcoded PCR products were sequenced using the SMRT Cell (1M v3 LR) platform by the Genome Technology Center Core Facility at New York University Langone Health. All primers used for PacBio sequencing were summarized in **table EV6**.

#### High-throughput sequencing

To detect indels in the high-throughput sequencing data, unmapped paired-end amplicon high-throughput sequencing reads were used as inputs into the CRISPResso2 tool to quantify the frequency of editing events(Pinello, Canver et al., 2016). The tool was run with default parameters (https://github.com/pinellolab/CRISPResso2).

#### PacBio sequencing

Raw PacBio data were demultiplexed with the corresponding barcode using the SMRTlink software to assign barcoded reads to each sample (smrtlink version: 8.0.0.80529, chemistry bundle: 8.0.0.778409, params: 8.0.0). Analysis of demultiplexed data was performed using PacBio tools distributed via Bioconda (https://github.com/PacificBiosciences/pbbioconda). For *DMD* exon 51 and 53 locus pileup, circular consensus sequences were converted to HiFi calls using the pbccs command and filtering for reads with support from at least three full-length subreads. The resulting fastq files were used as inputs to a custom python script that filtered for reads containing specific 50-bp index sequences at both the 5′ and 3′ regions of each read. Resulting filtered reads were mapped to the reference genome using minimap2 (ax splice -- splice-flank=no -u no -G 5000). The genome coverage of the alignment files was calculated using the “bedtools genomecov -d” (v 2.27.1) command with all downstream analyses performed using custom R script (v4.1.1) and visualized with the Gviz1 package(2016, Li, 2018). For *DMD* exon 51, the 5′ index sequence is tttttccaaacgtgcttttcaggaaacagtggtctgcttgttgaagtctg and the 3′ index sequence is aatcctggaccagaggttccattgagctgagatcacaccattgcactcca. For *DMD* exon 53, the 5′ index sequence is ggactatatttttgatttcatgttacaatcactagttttgtggggtcttt and the 3′ index sequence is tgatgtgtattgctgcagattcaatgtaagttcccgatacagataaagat.

#### Genome-wide off-target analysis

FASTQ files were provided by the Genome Technology Center Core Facility. FASTQ files were aligned to human genome reference build GRCh38 using BWA-MEM aligner (v0.7.17) followed by the GATK (v4.2.1.0) best practices pipeline.

The MarkDuplicatesSpark command was used to identify and mark PCR duplicate reads. BaseRecalibrator and ApplyBQSR commands were used along with known polymorphic sites to minimize systematic errors and improve downstream accuracy. Indel and SNP calling was done using Haplotypecaller, with the resulting file separated into SNPs and indels using the SelectVariants command. Variant filtering was performed using VariantFiltration with the following parameters: SNPs: QDLJ<LJ2.0, FSLJ>LJ60.0, SOR > 4.0, MQLJ<LJ40.0, MQRankSum < −12.5, ReadPosRankSum <LJ-8.0; Indels: QDLJ<LJ2.0, FSLJ>LJ200.0, SOR > 10.0, MQLJ<LJ40.0. To ensure pipeline robustness, the site of editing in the Cas9 sample was identified and confirmed for editing using a custom parsing script and IGV(McKenna, Hanna et al., 2010, Robinson, Thorvaldsdottir et al., 2011).

#### Predicted off-target analysis

Off-target events were predicted *in silico* using Cas-OFFinder allowing up to a 4-base mismatch with PAM sequences of either NGG or NAG (Bae et al., 2014). No off-target sites were predicted in this genome that were not found in WT samples.

#### Statistical analysis

All samples used to test CasPlus editing were assayed in duplicate unless otherwise noted in the figure legends. All data were calculated based on HTS results unless otherwise noted in the figure legends. Data were presented as mean and standard error of the mean (SEM). Data were generated and statistical analysis was performed using GraphPad Prism software.

## Acknowledgements

We thank C. Zhao, M. Khurram, and Y. Xiao for the cloning and sequencing of guide RNAs and plasmids; X. Wang, L. Dow, P. Hsu, D. Grimm, and L. Bu for subcloning or lentivirus packaging vector plasmids; H. Stower and R. Barr for manuscript editing; A. Namboodiripad, R. Iyer, M. Huang, and U. Andréo for comments and suggestions. This work was supported by grants from the Departmental Start-Up Grant, NYU Langone Health, and Kids Connect Charitable Fund.

## Author contributions

Q.Y. and C.L. designed, performed, and analyzed the experiments. J.S.A. analyzed the HTS, PacBio sequencing, and WGS data. M.M. helped with vectors cloning and sequencing. S.K performance the microinjection of mouse embryos. Q.Y. and C.L. drafted the original manuscript. Q.Y., C.L., G.R., and O.D. reviewed and edited the manuscripts. C.L. supervised the study. C.L. and Q.Y. conceptualized the study.

## Data and materials availability

All data needed to evaluate the conclusions in the paper are present in the paper and/or the Supplementary information. Additional data related to this paper may be requested from the authors.

## Supplementary information

Appendix Figs. S1-6 and tables EV1-8

## References

(2016) Statistical Genomics. Methods and Protocols. Anticancer Res 36: 3224

(2023) CRISPR CLINICAL TRIALS. CRISPR Medicine News

Abdus Sattar AK, Lin TC, Jones C, Konigsberg WH (1996) Functional consequences and exonuclease kinetic parameters of point mutations in bacteriophage T4 DNA polymerase. Biochemistry 35: 16621–9

Adikusuma F, Piltz S, Corbett MA, Turvey M, McColl SR, Helbig KJ, Beard MR, Hughes J, Pomerantz RT, Thomas PQ (2018) Large deletions induced by Cas9 cleavage. Nature 560: E8–E9

Adorisio R, Mencarelli E, Cantarutti N, Calvieri C, Amato L, Cicenia M, Silvetti M, D’Amico A, Grandinetti M, Drago F, Amodeo A (2020) Duchenne Dilated Cardiomyopathy: Cardiac Management from Prevention to Advanced Cardiovascular Therapies. J Clin Med 9

Allen F, Crepaldi L, Alsinet C, Strong AJ, Kleshchevnikov V, De Angeli P, Palenikova P, Khodak A, Kiselev V, Kosicki M, Bassett AR, Harding H, Galanty Y, Munoz-Martinez F, Metzakopian E, Jackson SP, Parts L (2018) Predicting the mutations generated by repair of Cas9-induced double-strand breaks. Nat Biotechnol

Amit I, Iancu O, Levy-Jurgenson A, Kurgan G, McNeill MS, Rettig GR, Allen D, Breier D, Ben Haim N, Wang Y, Anavy L, Hendel A, Yakhini Z (2021) CRISPECTOR provides accurate estimation of genome editing translocation and off-target activity from comparative NGS data. Nat Commun 12: 3042

Amoasii L, Long C, Li H, Mireault AA, Shelton JM, Sanchez-Ortiz E, McAnally JR, Bhattacharyya S, Schmidt F, Grimm D, Hauschka SD, Bassel-Duby R, Olson EN (2017) Single-cut genome editing restores dystrophin expression in a new mouse model of muscular dystrophy. Sci Transl Med 9

Anzalone AV, Koblan LW, Liu DR (2020) Genome editing with CRISPR-Cas nucleases, base editors, transposases and prime editors. Nat Biotechnol 38: 824-844

Anzalone AV, Randolph PB, Davis JR, Sousa AA, Koblan LW, Levy JM, Chen PJ, Wilson C, Newby GA, Raguram A, Liu DR (2019) Search-and-replace genome editing without double-strand breaks or donor DNA. Nature 576: 149–157

Bae S, Park J, Kim JS (2014) Cas-OFFinder: a fast and versatile algorithm that searches for potential off-target sites of Cas9 RNA-guided endonucleases. Bioinformatics 30: 1473–5

Bebenek K, Joyce CM, Fitzgerald MP, Kunkel TA (1990) The fidelity of DNA synthesis catalyzed by derivatives of Escherichia coli DNA polymerase I. J Biol Chem 265: 13878–87

Bermudez-Cabrera HC, Culbertson S, Barkal S, Holmes B, Shen MW, Zhang S, Gifford DK, Sherwood RI (2021) Small molecule inhibition of ATM kinase increases CRISPR-Cas9 1-bp insertion frequency. Nat Commun 12: 5111

Bothmer A, Gareau KW, Abdulkerim HS, Buquicchio F, Cohen L, Viswanathan R, Zuris JA, Marco E, Fernandez CA, Myer VE, Cotta-Ramusino C (2020) Detection and Modulation of DNA Translocations During Multi-Gene Genome Editing in T Cells. CRISPR J 3: 177–187

Boutin J, Cappellen D, Rosier J, Amintas S, Dabernat S, Bedel A, Moreau-Gaudry F (2022) ON-Target Adverse Events of CRISPR-Cas9 Nuclease: More Chaotic than Expected. CRISPR J 5: 19–30

Capson TL, Peliska JA, Kaboord BF, Frey MW, Lively C, Dahlberg M, Benkovic SJ (1992) Kinetic characterization of the polymerase and exonuclease activities of the gene 43 protein of bacteriophage T4. Biochemistry 31: 10984–94

Cejka P, Symington LS (2021) DNA End Resection: Mechanism and Control. Annu Rev Genet 55: 285–307

Chang HHY, Pannunzio NR, Adachi N, Lieber MR (2017) Non-homologous DNA end joining and alternative pathways to double-strand break repair. Nat Rev Mol Cell Biol 18: 495–506

Choi PS, Meyerson M (2014) Targeted genomic rearrangements using CRISPR/Cas technology. Nat Commun 5: 3728

Cong L, Ran FA, Cox D, Lin S, Barretto R, Habib N, Hsu PD, Wu X, Jiang W, Marraffini LA, Zhang F (2013) Multiplex genome engineering using CRISPR/Cas systems. Science 339: 819–23

De Waard A, Paul AV, Lehman IR (1965) The structural gene for deoxyribonucleic acid polymerase in bacteriophages T4 and T5. Proc Natl Acad Sci U S A 54: 1241-8

Depil S, Duchateau P, Grupp SA, Mufti G, Poirot L (2020) ’Off-the-shelf’ allogeneic CAR T cells: development and challenges. Nat Rev Drug Discov 19: 185–199

Duan D, Goemans N, Takeda S, Mercuri E, Aartsma-Rus A (2021) Duchenne muscular dystrophy. Nat Rev Dis Primers 7: 13

Ferreira da Silva J, Oliveira GP, Arasa-Verge EA, Kagiou C, Moretton A, Timelthaler G, Jiricny J, Loizou JI (2022) Prime editing efficiency and fidelity are enhanced in the absence of mismatch repair. Nat Commun 13: 760

Gaudelli NM, Komor AC, Rees HA, Packer MS, Badran AH, Bryson DI, Liu DR (2017) Programmable base editing of A*T to G*C in genomic DNA without DNA cleavage. Nature 551: 464–471

Geurts MH, de Poel E, Pleguezuelos-Manzano C, Oka R, Carrillo L, Andersson-Rolf A, Boretto M, Brunsveld JE, van Boxtel R, Beekman JM, Clevers H (2021) Evaluating CRISPR-based prime editing for cancer modeling and CFTR repair in organoids. Life Sci Alliance 4

Hogg M, Cooper W, Reha-Krantz L, Wallace SS (2006) Kinetics of error generation in homologous B-family DNA polymerases. Nucleic Acids Res 34: 2528–35

Hori K, Mark DF, Richardson CC (1979) Deoxyribonucleic acid polymerase of bacteriophage T7. Characterization of the exonuclease activities of the gene 5 protein and the reconstituted polymerase. J Biol Chem 254: 11598-604

Jiang F, Taylor DW, Chen JS, Kornfeld JE, Zhou K, Thompson AJ, Nogales E, Doudna JA (2016) Structures of a CRISPR-Cas9 R-loop complex primed for DNA cleavage. Science 351: 867–71

Jinek M, Chylinski K, Fonfara I, Hauer M, Doudna JA, Charpentier E (2012) A programmable dual-RNA-guided DNA endonuclease in adaptive bacterial immunity. Science 337: 816–21

Jinek M, East A, Cheng A, Lin S, Ma E, Doudna J (2013) RNA-programmed genome editing in human cells. Elife 2: e00471

Khan A, Sarkar E (2022) CRISPR/Cas9 encouraged CAR-T cell immunotherapy reporting efficient and safe clinical results towards cancer. Cancer Treat Res Commun 33: 100641

Kim HK, Yu G, Park J, Min S, Lee S, Yoon S, Kim HH (2021) Predicting the efficiency of prime editing guide RNAs in human cells. Nat Biotechnol 39: 198–206

Komor AC, Kim YB, Packer MS, Zuris JA, Liu DR (2016) Programmable editing of a target base in genomic DNA without double-stranded DNA cleavage. Nature 533: 420–4

Konermann S, Brigham MD, Trevino AE, Joung J, Abudayyeh OO, Barcena C, Hsu PD, Habib N, Gootenberg JS, Nishimasu H, Nureki O, Zhang F (2015) Genome-scale transcriptional activation by an engineered CRISPR-Cas9 complex. Nature 517: 583–8

Kosicki M, Allen F, Steward F, Tomberg K, Pan Y, Bradley A (2022) Cas9-induced large deletions and small indels are controlled in a convergent fashion. Nat Commun 13: 3422

Kosicki M, Tomberg K, Bradley A (2018) Repair of double-strand breaks induced by CRISPR-Cas9 leads to large deletions and complex rearrangements. Nat Biotechnol 36: 765–771

Kucera RB, Nichols NM (2008) DNA-dependent DNA polymerases. Curr Protoc Mol Biol Chapter 3: Unit3 5

Labanieh L, Mackall CL (2023) CAR immune cells: design principles, resistance and the next generation. Nature 614: 635–648

Leenay RT, Aghazadeh A, Hiatt J, Tse D, Roth TL, Apathy R, Shifrut E, Hultquist JF, Krogan N, Wu Z, Cirolia G, Canaj H, Leonetti MD, Marson A, May AP, Zou J (2019) Large dataset enables prediction of repair after CRISPR-Cas9 editing in primary T cells. Nat Biotechnol 37: 1034–1037

Leibowitz ML, Papathanasiou S, Doerfler PA, Blaine LJ, Sun L, Yao Y, Zhang CZ, Weiss MJ, Pellman D (2021) Chromothripsis as an on-target consequence of CRISPR-Cas9 genome editing. Nat Genet 53: 895–905

Lemos BR, Kaplan AC, Bae JE, Ferrazzoli AE, Kuo J, Anand RP, Waterman DP, Haber JE (2018) CRISPR/Cas9 cleavages in budding yeast reveal templated insertions and strand-specific insertion/deletion profiles. Proc Natl Acad Sci U S A 115: E2040–E2047

Li H (2018) Minimap2: pairwise alignment for nucleotide sequences. Bioinformatics 34: 3094–3100

Liu Y, Tao W, Wen S, Li Z, Yang A, Deng Z, Sun Y (2015) In Vitro CRISPR/Cas9 System for Efficient Targeted DNA Editing. mBio 6: e01714–15

Long C, Amoasii L, Mireault AA, McAnally JR, Li H, Sanchez-Ortiz E, Bhattacharyya S, Shelton JM, Bassel-Duby R, Olson EN (2016) Postnatal genome editing partially restores dystrophin expression in a mouse model of muscular dystrophy. Science 351: 400–3

Long C, Li H, Tiburcy M, Rodriguez-Caycedo C, Kyrychenko V, Zhou H, Zhang Y, Min YL, Shelton JM, Mammen PPA, Liaw NY, Zimmermann WH, Bassel-Duby R, Schneider JW, Olson EN (2018) Correction of diverse muscular dystrophy mutations in human engineered heart muscle by single-site genome editing. Sci Adv 4: eaap9004

Long C, McAnally JR, Shelton JM, Mireault AA, Bassel-Duby R, Olson EN (2014) Prevention of muscular dystrophy in mice by CRISPR/Cas9-mediated editing of germline DNA. Science 345: 1184–1188

Mali P, Yang L, Esvelt KM, Aach J, Guell M, DiCarlo JE, Norville JE, Church GM (2013) RNA-guided human genome engineering via Cas9. Science 339: 823–6

Mao Z, Bozzella M, Seluanov A, Gorbunova V (2008) Comparison of nonhomologous end joining and homologous recombination in human cells. DNA Repair (Amst) 7: 1765–71

McKenna A, Hanna M, Banks E, Sivachenko A, Cibulskis K, Kernytsky A, Garimella K, Altshuler D, Gabriel S, Daly M, DePristo MA (2010) The Genome Analysis Toolkit: a MapReduce framework for analyzing next-generation DNA sequencing data. Genome Res 20: 1297–303

Muntoni F, Torelli S, Ferlini A (2003) Dystrophin and mutations: one gene, several proteins, multiple phenotypes. Lancet Neurol 2: 731–40

Nahmad AD, Reuveni E, Goldschmidt E, Tenne T, Liberman M, Horovitz-Fried M, Khosravi R, Kobo H, Reinstein E, Madi A, Ben-David U, Barzel A (2022a) Frequent aneuploidy in primary human T cells after CRISPR-Cas9 cleavage. Nat Biotechnol

Nahmad AD, Reuveni E, Goldschmidt E, Tenne T, Liberman M, Horovitz-Fried M, Khosravi R, Kobo H, Reinstein E, Madi A, Ben-David U, Barzel A (2022b) Frequent aneuploidy in primary human T cells after CRISPR-Cas9 cleavage. Nat Biotechnol 40: 1807–1813

Nelson CE, Wu Y, Gemberling MP, Oliver ML, Waller MA, Bohning JD, Robinson-Hamm JN, Bulaklak K, Castellanos Rivera RM, Collier JH, Asokan A, Gersbach CA (2019) Long-term evaluation of AAV-CRISPR genome editing for Duchenne muscular dystrophy. Nat Med 25: 427–432

O’Brien KF, Kunkel LM (2001) Dystrophin and muscular dystrophy: past, present, and future. Mol Genet Metab 74: 75–88

Olson EN (2021) Toward the correction of muscular dystrophy by gene editing. Proc Natl Acad Sci U S A 118

Oshima J, Magner DB, Lee JA, Breman AM, Schmitt ES, White LD, Crowe CA, Merrill M, Jayakar P, Rajadhyaksha A, Eng CM, del Gaudio D (2009) Regional genomic instability predisposes to complex dystrophin gene rearrangements. Hum Genet 126: 411–23

Owens DDG, Caulder A, Frontera V, Harman JR, Allan AJ, Bucakci A, Greder L, Codner GF, Hublitz P, McHugh PJ, Teboul L, de Bruijn M (2019) Microhomologies are prevalent at Cas9-induced larger deletions. Nucleic Acids Res 47: 7402–7417

Papathanasiou S, Markoulaki S, Blaine LJ, Leibowitz ML, Zhang CZ, Jaenisch R, Pellman D (2021) Whole chromosome loss and genomic instability in mouse embryos after CRISPR-Cas9 genome editing. Nat Commun 12: 5855

Pinello L, Canver MC, Hoban MD, Orkin SH, Kohn DB, Bauer DE, Yuan GC (2016) Analyzing CRISPR genome-editing experiments with CRISPResso. Nat Biotechnol 34: 695–7

Poirot L, Philip B, Schiffer-Mannioui C, Le Clerre D, Chion-Sotinel I, Derniame S, Potrel P, Bas C, Lemaire L, Galetto R, Lebuhotel C, Eyquem J, Cheung GW, Duclert A, Gouble A, Arnould S, Peggs K, Pule M, Scharenberg AM, Smith J (2015) Multiplex Genome-Edited T-cell Manufacturing Platform for "Off-the-Shelf" Adoptive T-cell Immunotherapies. Cancer Res 75: 3853–64

Ramsden DA, Asagoshi K (2012) DNA polymerases in nonhomologous end joining: are there any benefits to standing out from the crowd? Environ Mol Mutagen 53: 741–51

Ran FA, Hsu PD, Wright J, Agarwala V, Scott DA, Zhang F (2013) Genome engineering using the CRISPR-Cas9 system. Nat Protoc 8: 2281–2308

Reha-Krantz LJ (1988) Amino acid changes coded by bacteriophage T4 DNA polymerase mutator mutants. Relating structure to function. J Mol Biol 202: 711–24

Reha-Krantz LJ (1998) Regulation of DNA polymerase exonucleolytic proofreading activity: studies of bacteriophage T4 "antimutator" DNA polymerases. Genetics 148: 1551–7

Reha-Krantz LJ, Stocki S, Nonay RL, Dimayuga E, Goodrich LD, Konigsberg WH, Spicer EK (1991) DNA polymerization in the absence of exonucleolytic proofreading: in vivo and in vitro studies. Proc Natl Acad Sci U S A 88: 2417–21

Robinson JT, Thorvaldsdottir H, Winckler W, Guttman M, Lander ES, Getz G, Mesirov JP (2011) Integrative genomics viewer. Nat Biotechnol 29: 24–6

Sfeir A, Symington LS (2015) Microhomology-Mediated End Joining: A Back-up Survival Mechanism or Dedicated Pathway? Trends Biochem Sci 40: 701–714

Shen MW, Arbab M, Hsu JY, Worstell D, Culbertson SJ, Krabbe O, Cassa CA, Liu DR, Gifford DK, Sherwood RI (2018) Predictable and precise template-free CRISPR editing of pathogenic variants. Nature 563: 646–651

Sheridan C (2022a) Off-the-shelf, gene-edited CAR-T cells forge ahead, despite safety scare. nature biotechnology

Sheridan C (2022b) Off-the-shelf, gene-edited CAR-T cells forge ahead, despite safety scare. Nat Biotechnol 40: 5–8

Shi X, Shou J, Mehryar MM, Li J, Wang L, Zhang M, Huang H, Sun X, Wu Q (2019) Cas9 has no exonuclease activity resulting in staggered cleavage with overhangs and predictable di- and tri-nucleotide CRISPR insertions without template donor. Cell Discov 5: 53

Shou J, Li J, Liu Y, Wu Q (2018) Precise and Predictable CRISPR Chromosomal Rearrangements Reveal Principles of Cas9-Mediated Nucleotide Insertion. Mol Cell 71: 498–509 e4

Stadtmauer EA, Fraietta JA, Davis MM, Cohen AD, Weber KL, Lancaster E, Mangan PA, Kulikovskaya I, Gupta M, Chen F, Tian L, Gonzalez VE, Xu J, Jung IY, Melenhorst JJ, Plesa G, Shea J, Matlawski T, Cervini A, Gaymon AL et al. (2020) CRISPR-engineered T cells in patients with refractory cancer. Science 367

Tomomi Aida JJW, Lixin Yang, Yuanyuan Hou, Mengqi Li, Dongdong Xu, Jianbang Lin, Peimin Qi, Zhonghua Lu, Guoping Feng (2020) Prime editing primarily induces undesired outcomes in mice. bioRxiv

Uddin F, Rudin CM, Sen T (2020) CRISPR Gene Therapy: Applications, Limitations, and Implications for the Future. Front Oncol 10: 1387

Xin C, Yin J, Yuan S, Ou L, Liu M, Zhang W, Hu J (2022) Comprehensive assessment of miniature CRISPR-Cas12f nucleases for gene disruption. Nat Commun 13: 5623

Yin J, Lu R, Xin C, Wang Y, Ling X, Li D, Zhang W, Liu M, Xie W, Kong L, Si W, Wei P, Xiao B, Lee HY, Liu T, Hu J (2022) Cas9 exo-endonuclease eliminates chromosomal translocations during genome editing. Nat Commun 13: 1204

Yoo KW, Yadav MK, Song Q, Atala A, Lu B (2022) Targeting DNA polymerase to DNA double-strand breaks reduces DNA deletion size and increases templated insertions generated by CRISPR/Cas9. Nucleic Acids Res 50: 3944–3957

Zetsche B, Gootenberg JS, Abudayyeh OO, Slaymaker IM, Makarova KS, Essletzbichler P, Volz SE, Joung J, van der Oost J, Regev A, Koonin EV, Zhang F (2015) Cpf1 is a single RNA-guided endonuclease of a class 2 CRISPR-Cas system. Cell 163: 759–71

Zhang Y, Li H, Min YL, Sanchez-Ortiz E, Huang J, Mireault AA, Shelton JM, Kim J, Mammen PPA, Bassel-Duby R, Olson EN (2020) Enhanced CRISPR-Cas9 correction of Duchenne muscular dystrophy in mice by a self-complementary AAV delivery system. Sci Adv 6: eaay6812

Zhang Y, Li H, Nishiyama T, McAnally JR, Sanchez-Ortiz E, Huang J, Mammen PPA, Bassel-Duby R, Olson EN (2022) A humanized knockin mouse model of Duchenne muscular dystrophy and its correction by CRISPR-Cas9 therapeutic gene editing. Mol Ther Nucleic Acids 29: 525–537

