## Supplemental materials for "Phage DNA polymerase prevents deleterious on-target DNA damage and enhances precise CRISPR/Cas9 editing"

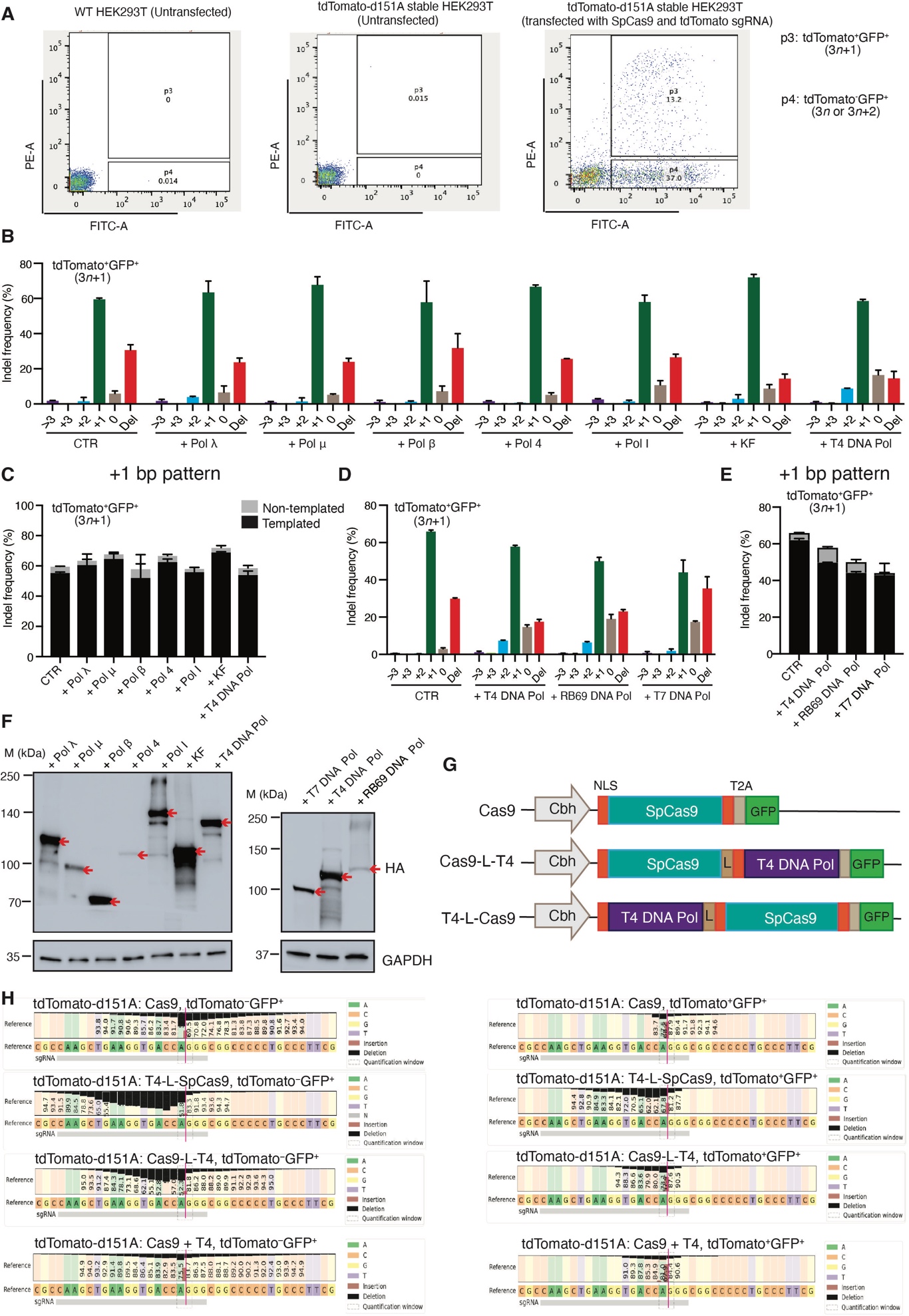

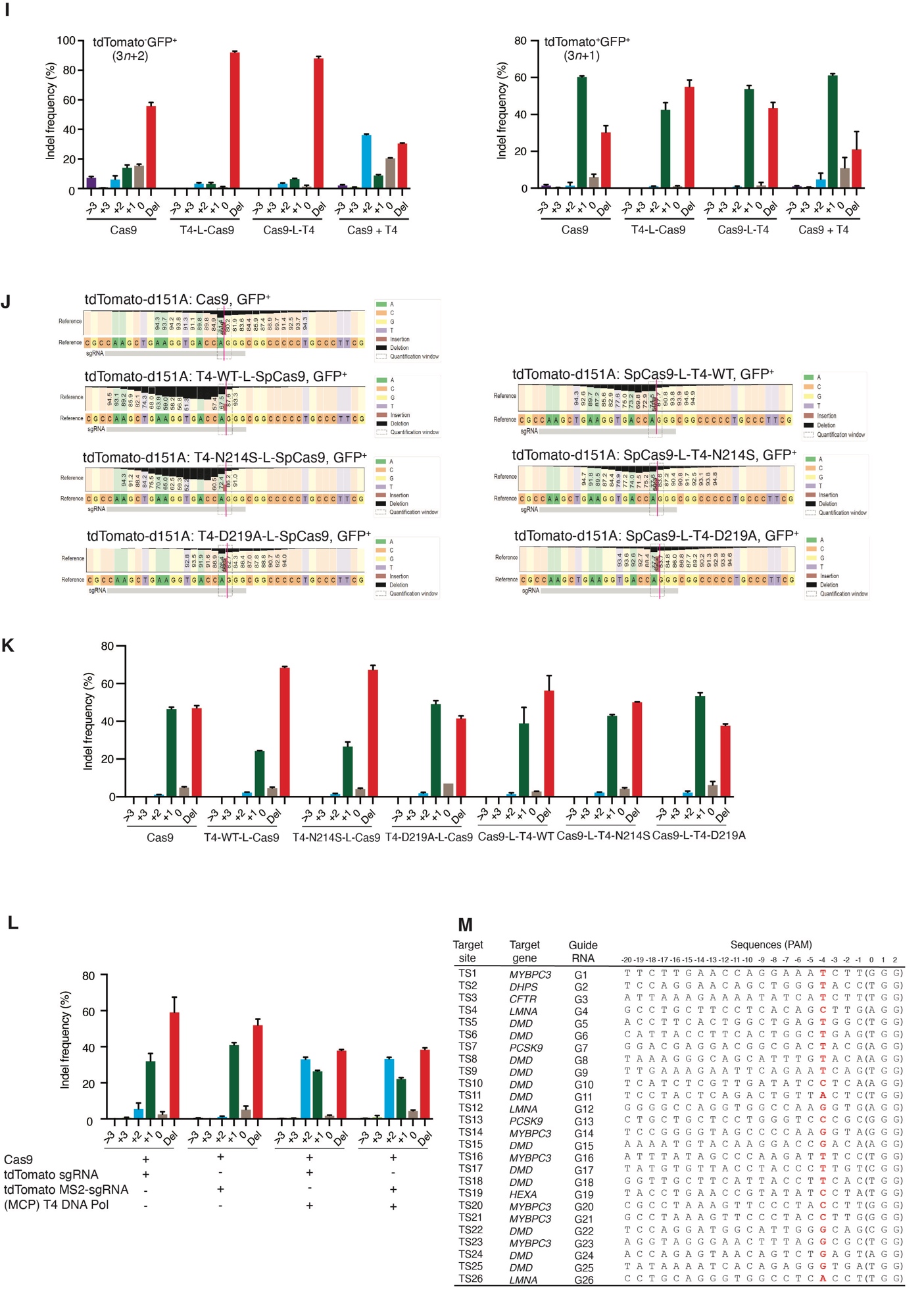

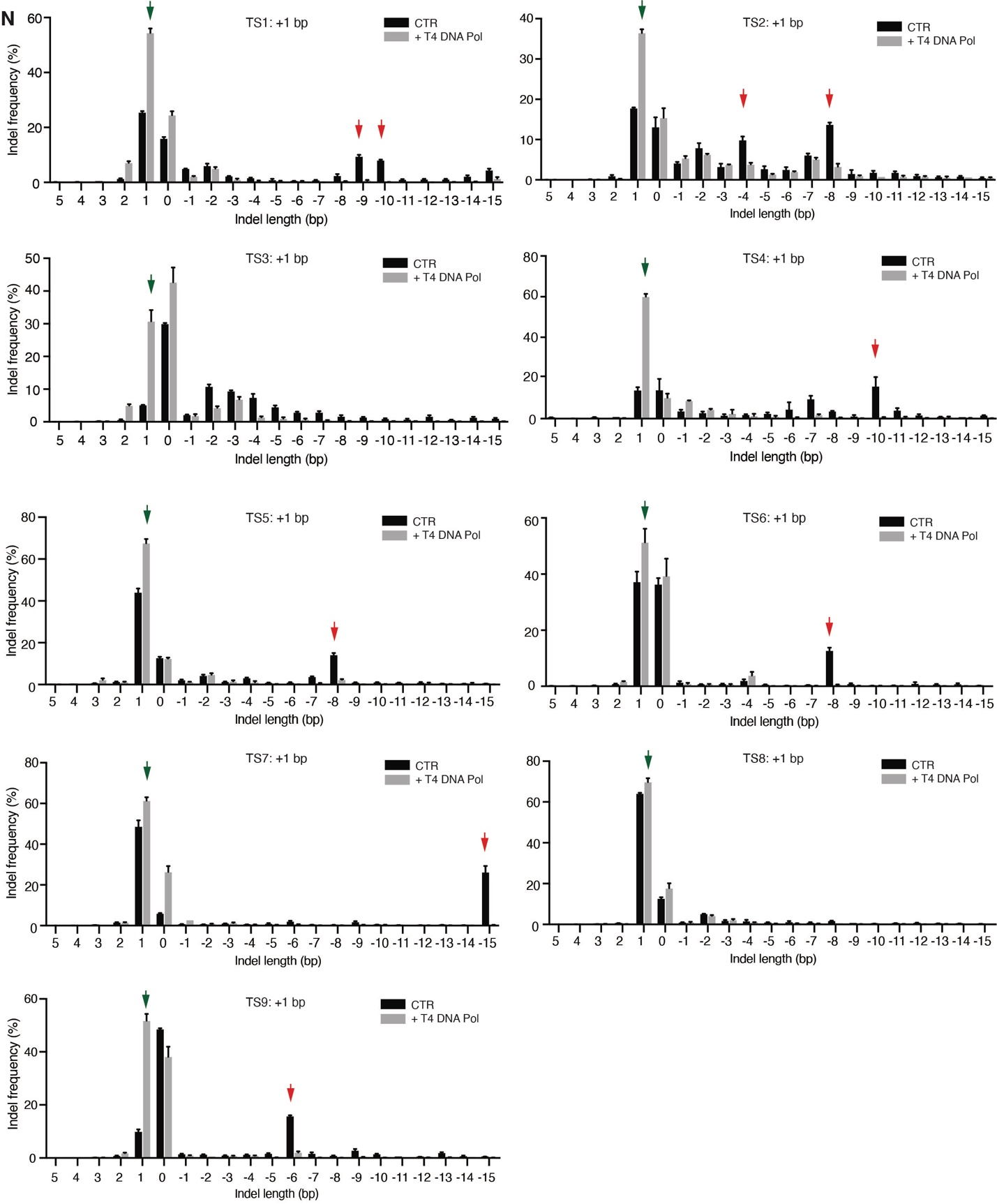

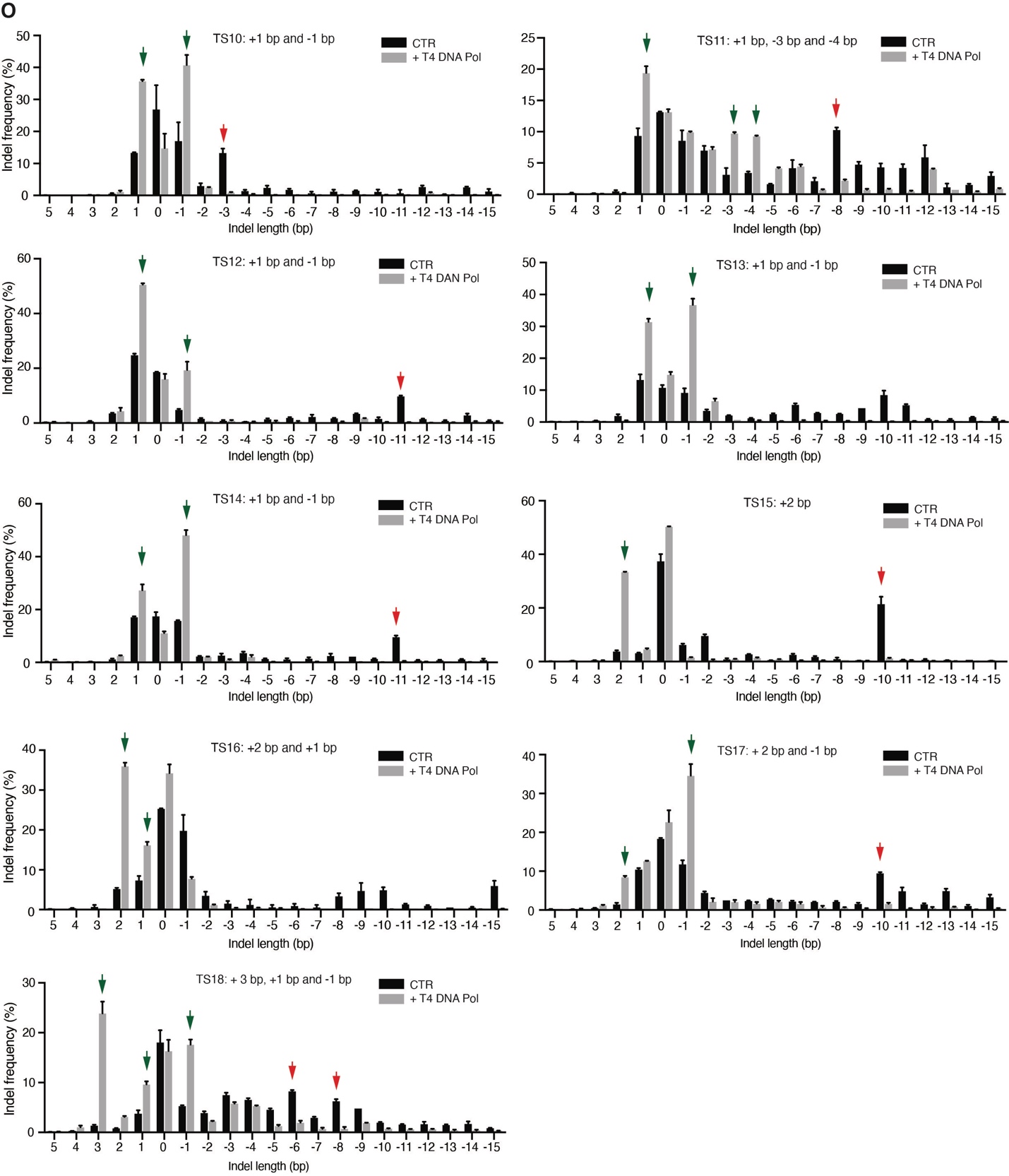

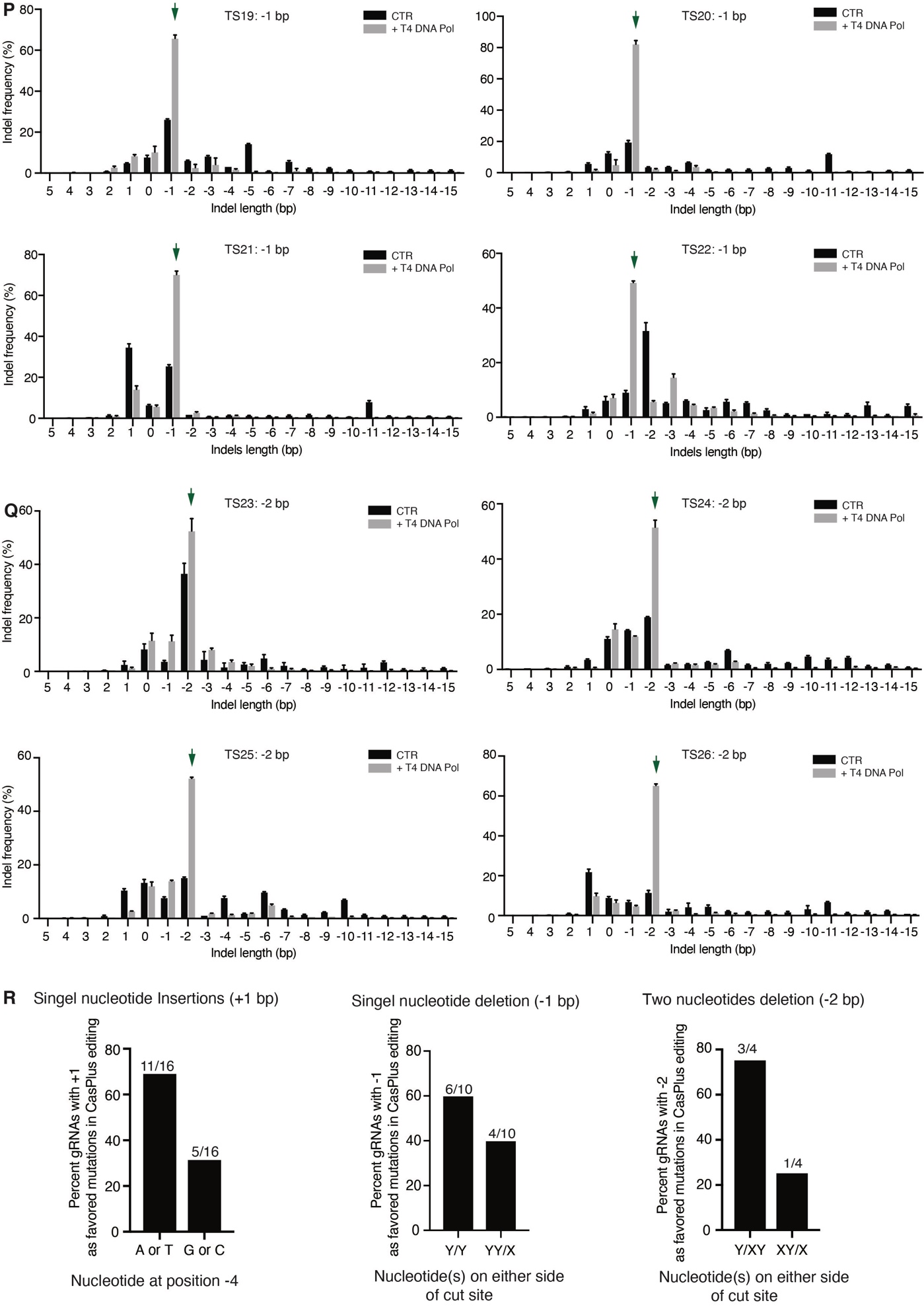

**Appendix Figure 1.** **T4 DNA polymerase favors small insertions over deletions at endogenous genome sites during Cas9 editing**. (**A**) Representive FACS plots demonstrate leakiness of the tdTomato reporter system and the relative proportions of tdTomato+GFP+ and tdTomato-GFP+ populations when transfected with Cas9 and tdTomato sgRNA. (**B**) Frequency of Cas9-induced indels in tdTomato^+^GFP^+^ populations with or without co-expression of a distinct DNA polymerase. (**C**) Pattern of 1-bp insertions in **B**. (**D**) Frequency of Cas9-induced indels in tdTomato^+^GFP^+^ populations with or without co-expression of a phage-derived DNA polymerase. (**E**) Pattern of 1-bp insertions in **D**. (**F**) Western bot analysis showing protein expression in unsorted tdTomato-d151A reporter cells after co-transfection with Cas9 and distinct DNA polymerases. (**G**) Architecture of vectors expressing the fusion proteins of Cas9 and T4 DNA polymerase. (**H**) Sequencing depth around the tdTomato-d151A mutation site. HEK293T tdTomato-d151A reporter cells were transfected with vectors expressing tdTomato-sgRNA and Cas9, T4-L-Cas9, or Cas9-L-T4 or co-transfected with a vector expressing tdTomato-sgRNA and Cas9 combined with a vector expressing T4 DNA polymerase. Transfected cells were then sorted into tdTomato^–^GFP^+^ and tdTomato^+^GFP^–^ populations, and processed for DNA amplification and HTS. Red vertical line, Cas9 cleavage site. (**I**) Frequency of indels observed in samples described in **H**. Cas9 fused with T4 DNA polymerase favors deletions in comparison with Cas9. (**J**) Sequencing depth around the tdTomato-d151A mutation site. HEK293T tdTomato-d151A reporter cells were transfected with vectors expressing tdTomato-sgRNA and Cas9, Cas9 fused with T4-WT (T4-WT-L-Cas9 and Cas9-L-T4-WT), Cas9 fused with DNA polymerase activity-deactivated T4 DNA polymerase (T4-N214S-L-Cas9 and Cas9-L-T4-N214S), or Cas9 fused exonuclease activity-deactivated T4 DNA polymerase (T4-D219A-L-Cas9 and Cas9-L-T4-D219A). Transfected cells (GFP^+^) were then sorted and processed for DNA amplification and HTS. (**K**) Frequency of indels observed in samples described in J. (**L**) Frequency of Cas9-induced indels in transfected cells (GFP^+^) with or without co-expression of T4 DNA polymerase combined with tdTomato-sgRNA or tdTomato MS2-sgRNA. (**M**) Table of target sites and guide RNAs used to test CasPlus editing in HEK293T cells. Nucleotide at position –4 (counting the NGG PAM as nucleotides 0–2) in each guide RNA was highlighted in red. (**N**-**Q**) Cas9-induced indel profiles and frequencies in cells with or without T4 DNA polymerase. Green arrowheads indicate that the frequencies of the insertion(s) or deletion(s) were substantially increased in cells with Cas9 and T4 DNA polymerase compared to Cas9-only (CTR). Red arrowheads indicate the deletions that arise from MMEJ repair in CTR cells. Compared to Cas9-only, Guide RNAs with Cas9 and T4 DNA polymerase in **N** favored 1-bp insertions, in **P** favored 1-bp deletions, in **Q** biased 2-bp deletions and in **O** enhanced the frequencies of both 1-bp insertions and small deletions (<3 bp) or 2–3-bp insertions. (**R**) Guide RNAs containing an A or T at position -4 are more likely to produce 1-bp insertions versus C or G in CasPlus editing. Among 10 guide RNAs that favored 1-bp deletions in CasPlus editing, six contained a repeating nucleotide at positions -4 and -3 (Y/Y), while four contained an identical nucleotide at positions -5 and -4 (YY/X). Among 4 guide RNAs that had a biased for 2-bp deletions in CasPlus editing, three contained a repeating nucleotide at positions -4 and -2 (Y/XY) while one contained a repeating nucleotide on -5 and -3 (XY/X). /, cut site. X and Y represent any nucleotide.

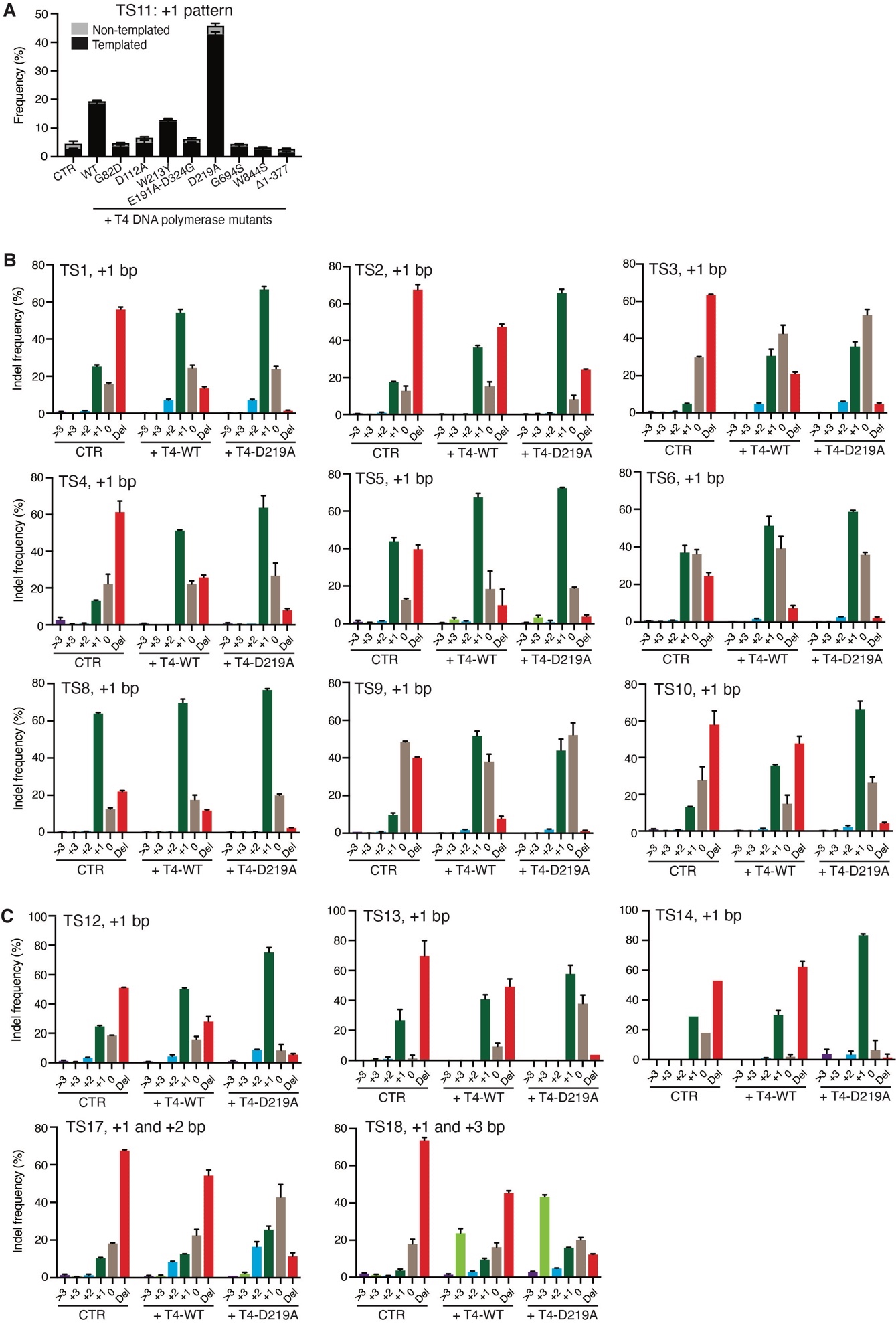

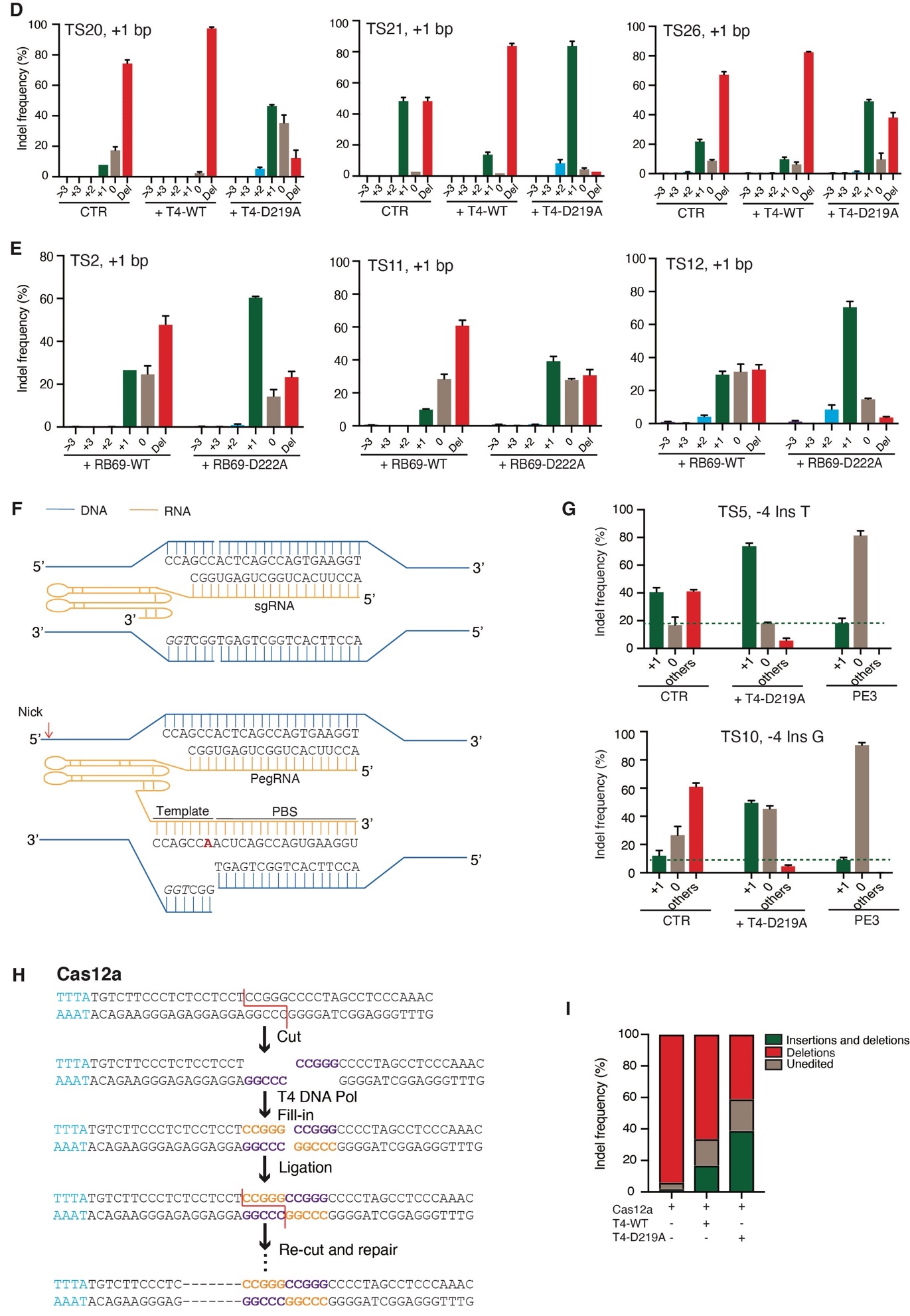

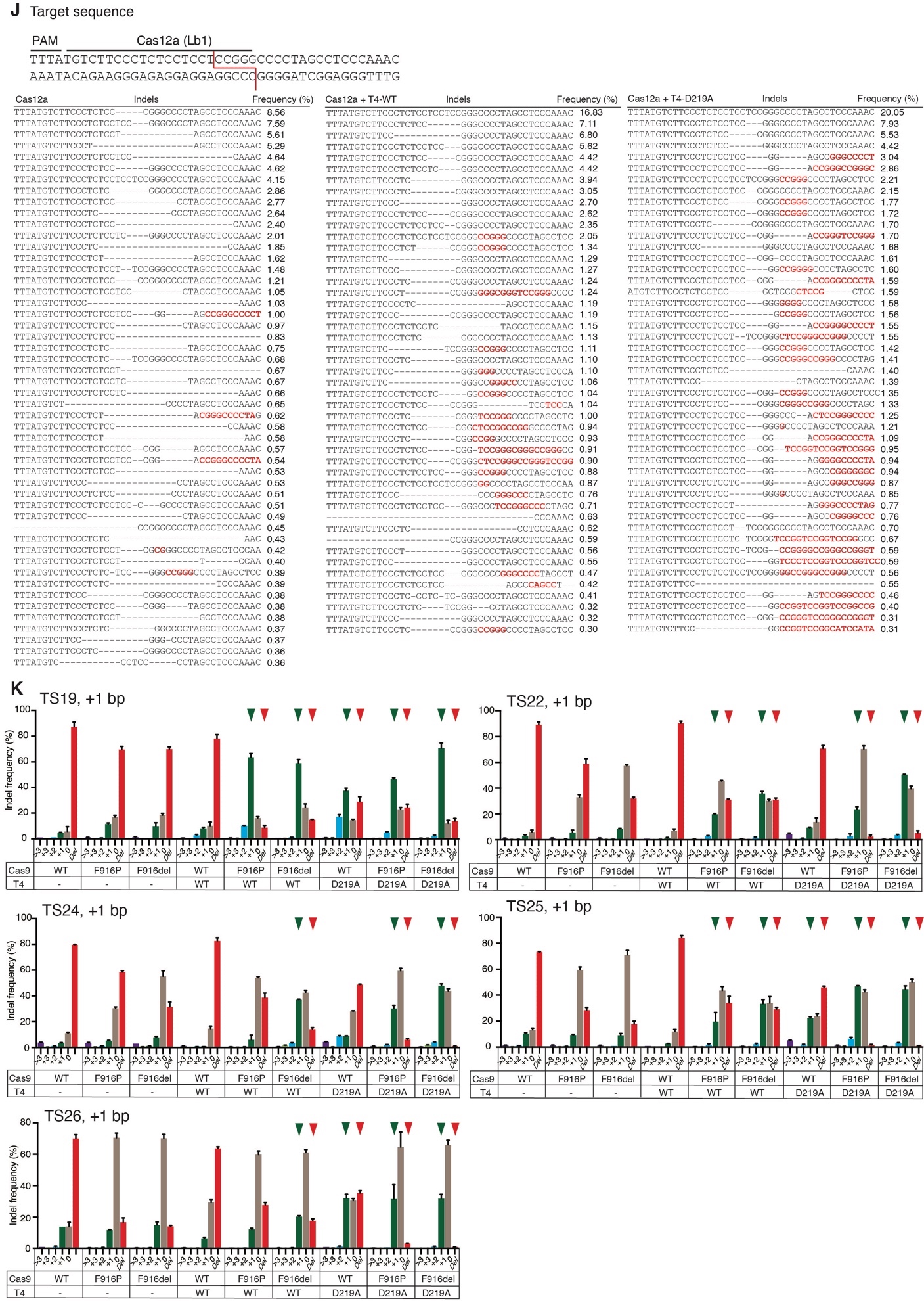

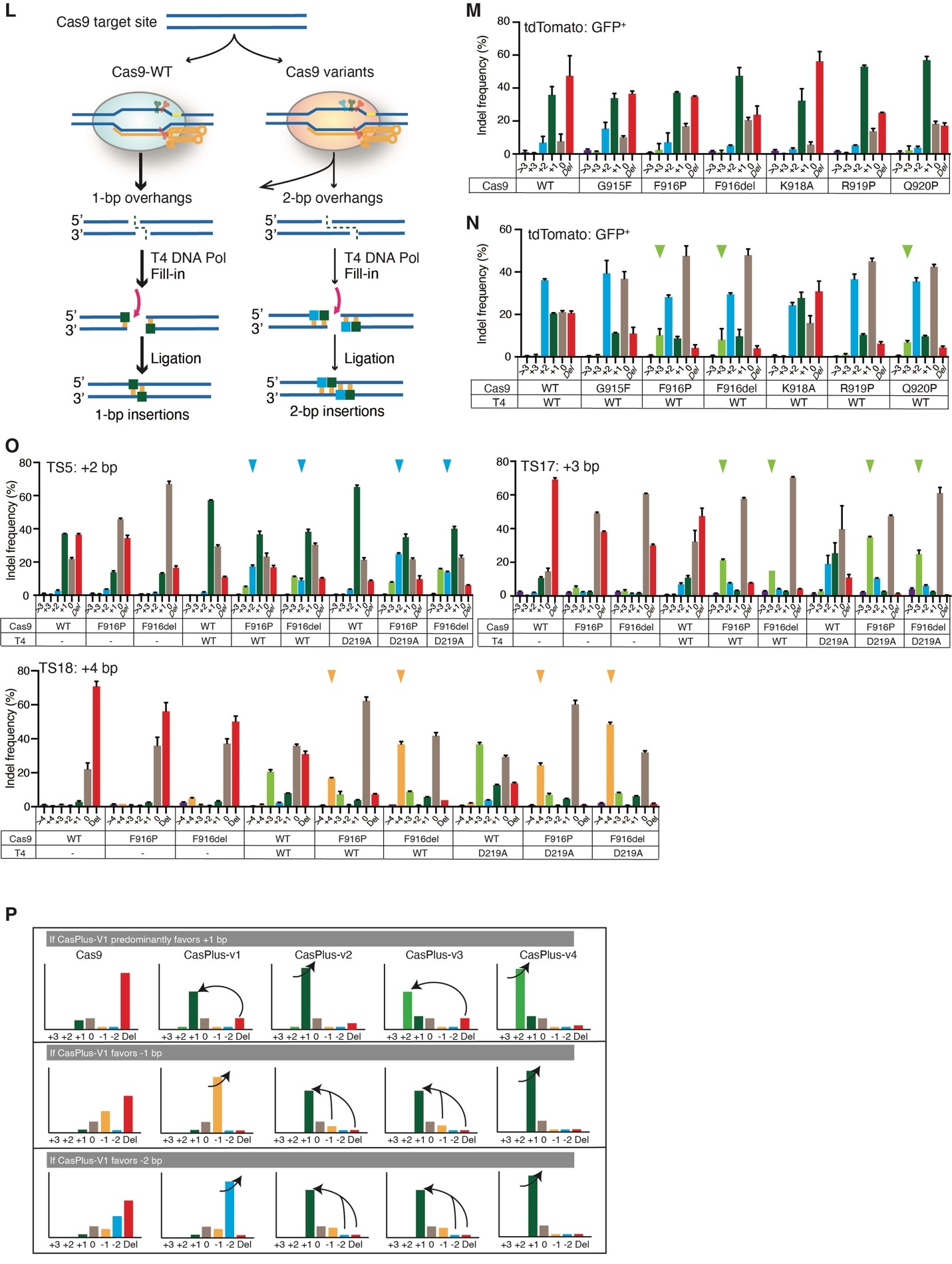
**Appendix Figure 2.** **Combination of** **engineered T4 or RB69 DNA polymerase mutant and Cas9 variants improves CasPlus editing efficiency.** (**A**) Pattern of 1-bp insertions at TS11 in **Fig 2C**. (**B**) Frequencies of Cas9-induced indels at TS1, TS2, TS3, TS4, TS5, TS6, TS8, TS9, and TS10 in cells with or without co-expression of T4-WT or T4-D219A. (**C-D**) Frequencies of Cas9-induced indels at TS12, TS13, TS14, TS17, and TS18 (**C**) or TS20, TS21, and TS26 (**D**) in cells with or without co-expression of T4-WT or T4-D219A. For TS13, TS14, TS20, and TS21, indel frequencies were calculated based on ICE analysis on Sanger sequencing files. (**E**) Frequencies of Cas9-induced indels at TS2, TS11, and TS12 in cells with co-expression of RB69-WT or RB69-D222A. (**F**) Schematics illustrating an example of the sgRNA used for Cas9 and CasPlus editing and the corresponding pegRNA used for prime editing. Red arrowhead, the second nick location. (**G**) Frequencies of targeted 1-bp insertions induced by Cas9 (CTR), CasPlus editing with T4-D219A, and PE3 editing. Others, other indels. (**H**) An example of the editing process that potentially occurs at a Cas12a cut site with the addition of T4 DNA polymerase. (**I**) Frequency of DNA alleles with insertions and deletions or only deletions induced by Cas12a alone or Cas12a combined with T4-WT or T4-D219A. (**J**) Indel profiles and frequencies in cells transfected with Cas12a alone, or co-transfected with Cas12a and T4-WT or T4-D219A. Red line, cleavage sites; Dashed lines, deletions; Red color letters, insertions. Top panel shows the guide RNA Lb1 sequences. Only indels with frequency ≥ 0.30 were shown here. (**K**) Frequencies of Cas9-induced indels at TS19, TS22, TS24, TS25, and TS26 with Cas9-WT or Cas9 variants individually or along with T4-WT or T4-D219A. (**L**) Schematics showing at target sites where Cas9-WT often produces DSB ends with 1-bp overhangs, leading to the production of 1-bp insertions (left), engineered Cas9 variants can facilitate the generation of 2-bp overhangs, enabling the production of 2-bp insertions with the addition of T4 DNA polymerase (right). (**M-N**) Frequencies of Cas9-induced indels at tdTomato-d151A site in GFP^+^ populations in tdTomato-d151A reporter cells transfected with Cas9-WT or Cas9 variants individually (**M**) or along with T4-WT (**N**). (**O**) Frequencies of Cas9-induced indels at TS5, TS17, and TS18 in cells transfected with Cas9-WT or Cas9 variants F916P or F916del individually or in conjunction with T4-WT or T4-D219A. (**P**) General rules following different versions of CasPlus editing.

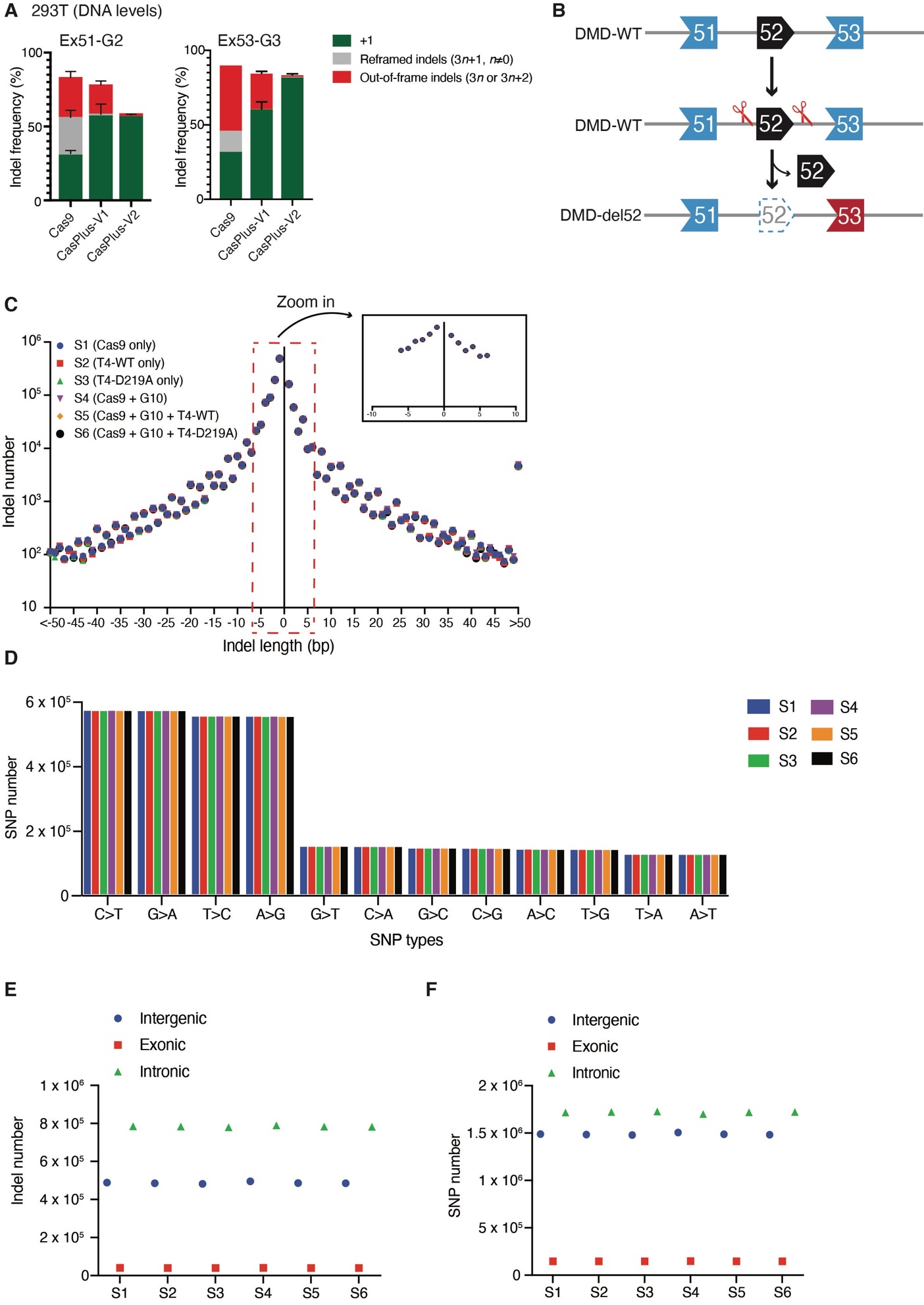

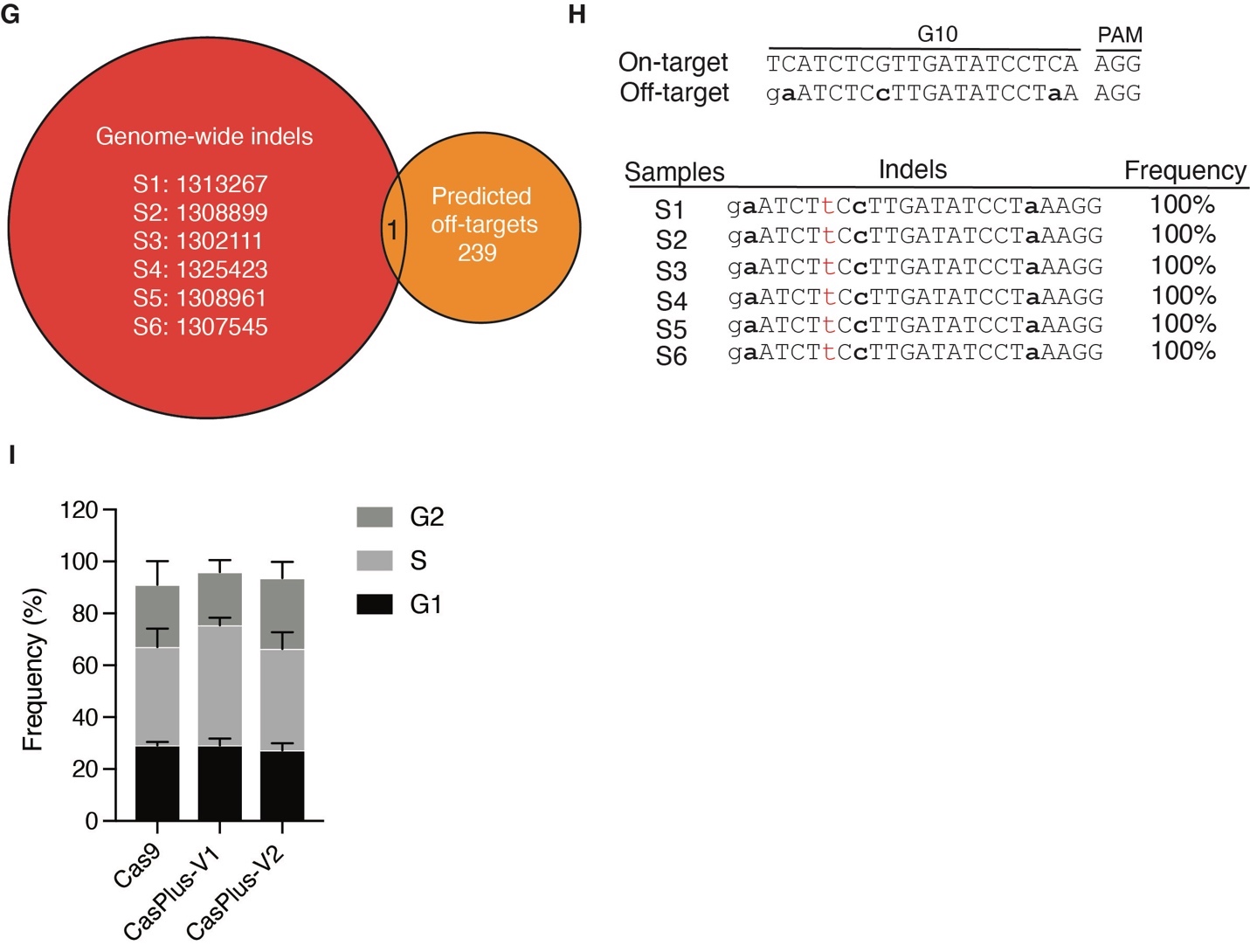

**Appendix Figure 3**. **Evaluation of efficiency and safety of CasPlus editing in correcting *DMD* del52 deletion.** (**A**) Frequencies of in-frame (3*n*+1, n≠0) and out-of-frame indels (3*n* or 3*n*+2) induced by Cas9, CasPlus-V1, and CasPlus-V2 with Ex51-G2 (left) or Ex53-G3 (right) in HEK293T cells. Frequencies were calculated based on ICE analysis on Sanger sequencing files. (**B**) Generation of *DMD* exon 52 deletion in iPSCs via dual guide RNA-mediated removal of exon 52. (**C-D**) Quantity of reads containing indels (**C**) or SNPs **(D**) in HEK293T cells transfected with vectors expressing either only Cas9 (S1), T4-WT (S2), or T4-D219A (S3) proteins; or Cas9 protein and gRNA G10 (S4); or Cas9 protein and gRNA G10 in combination with either T4-WT (S5) or T4-D219A (S6). (**E-F**). Distribution of indels (**E)** and SNPs (**F**). (**G**) Total number of off-target sites detected by WGS and predicted by Cas-OFFinder, and the number of overlapping sites. (**H**) Sequences and editing efficiency detected by WGS in each sample at the off-target site identified in all samples. It was a homozygous insertion of thymidine that was present in all treatment groups. (I) Cell cycle analysis using PI in HEK293T cells transfected with vectors expressing Cas9 protein and gRNA G10 (Cas9) or Cas9 protein and gRNA G10 in combination with either T4-WT (CasPlus-V1) or T4-D219A (CasPlus-V2). Total 10,000 transfected cells each group were analyzed. The percentage of cells in distinct cell cycle phases calculated using Flow Jo software. Values and error bars reflect mean ± SEM of n=3 replicates. G1, S or G2 phase, Cas9 verse CasPlus-V1, p>0.05; Cas9 verse CasPlus-V2, p>0.05.

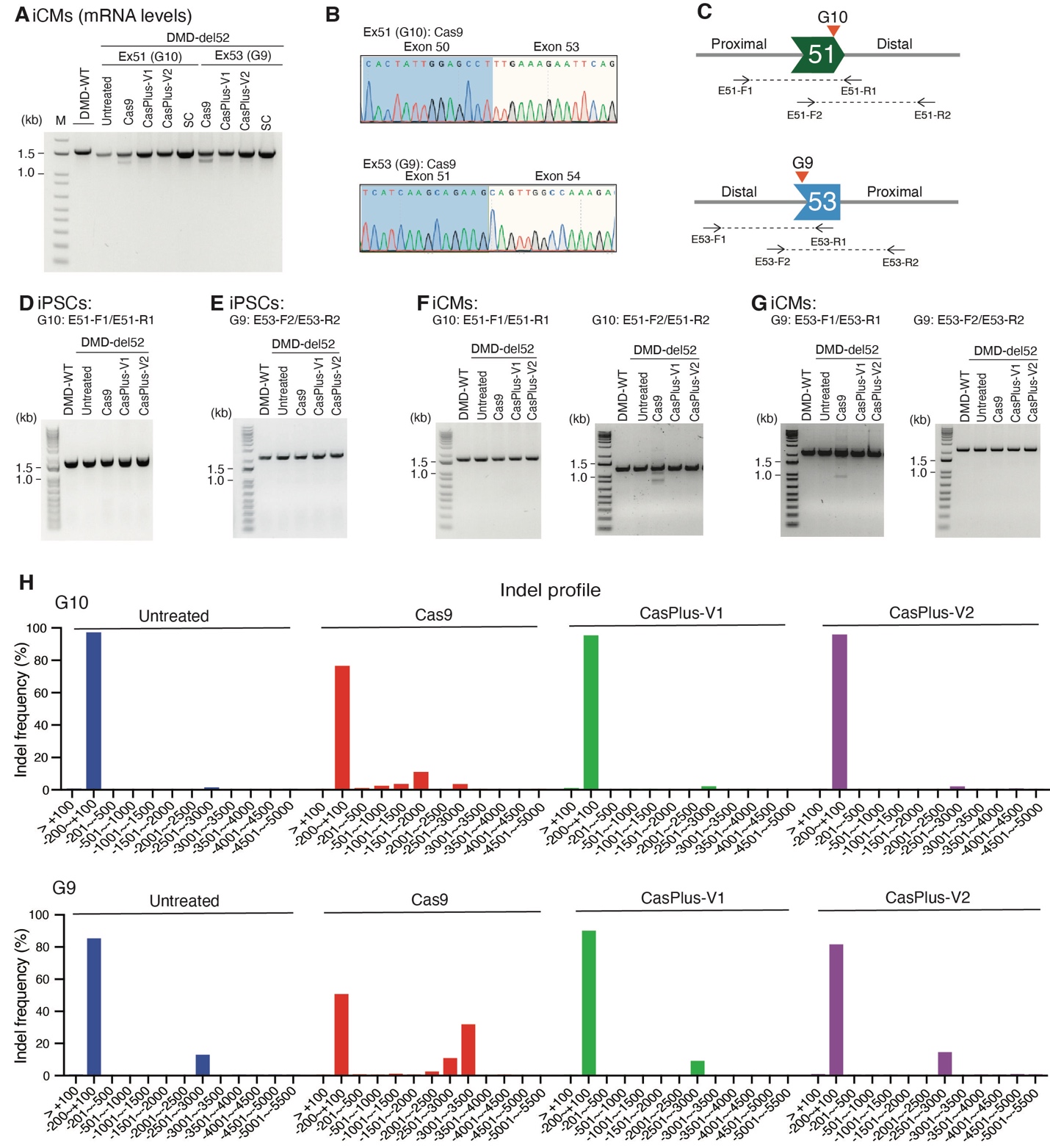

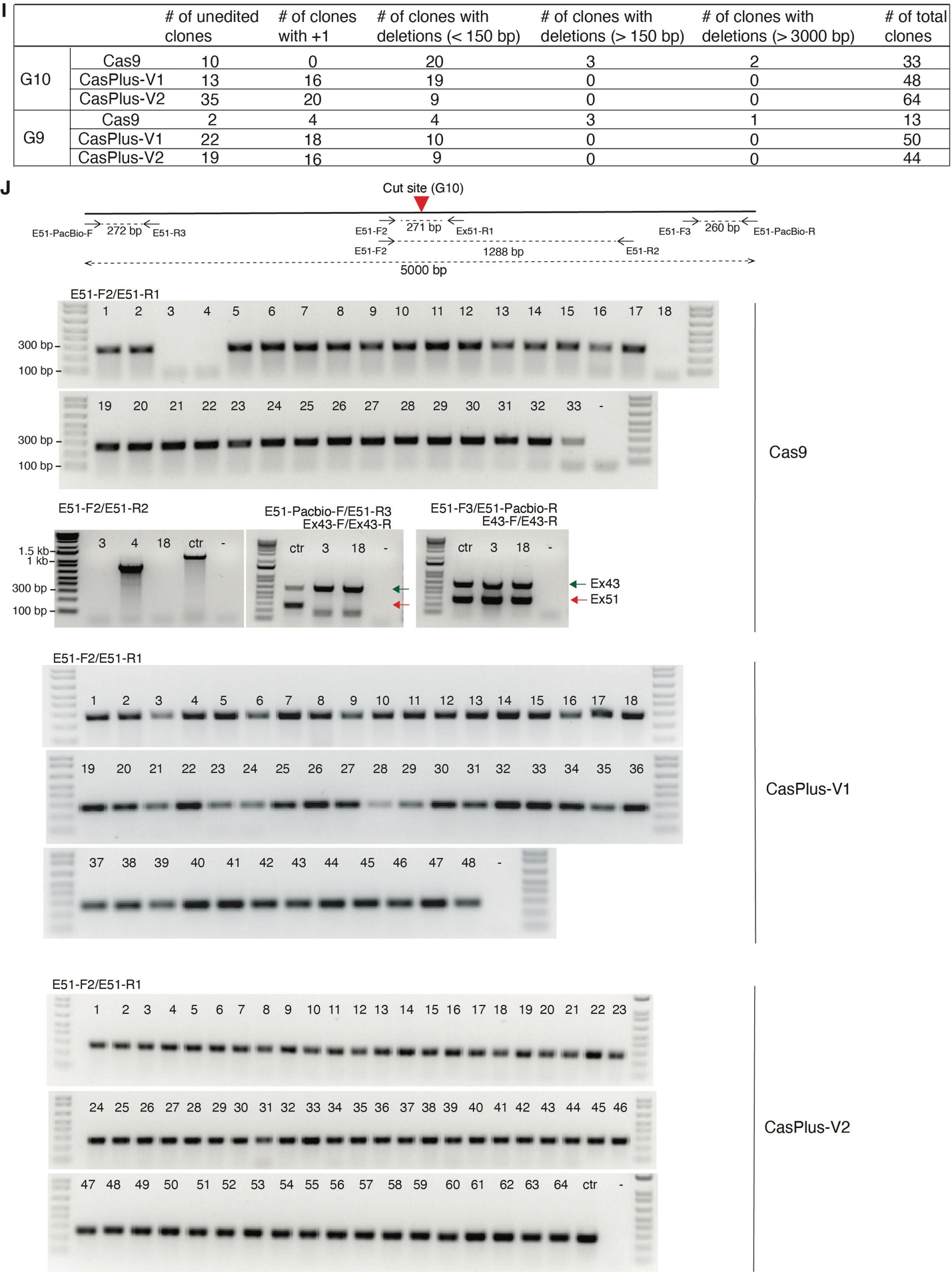

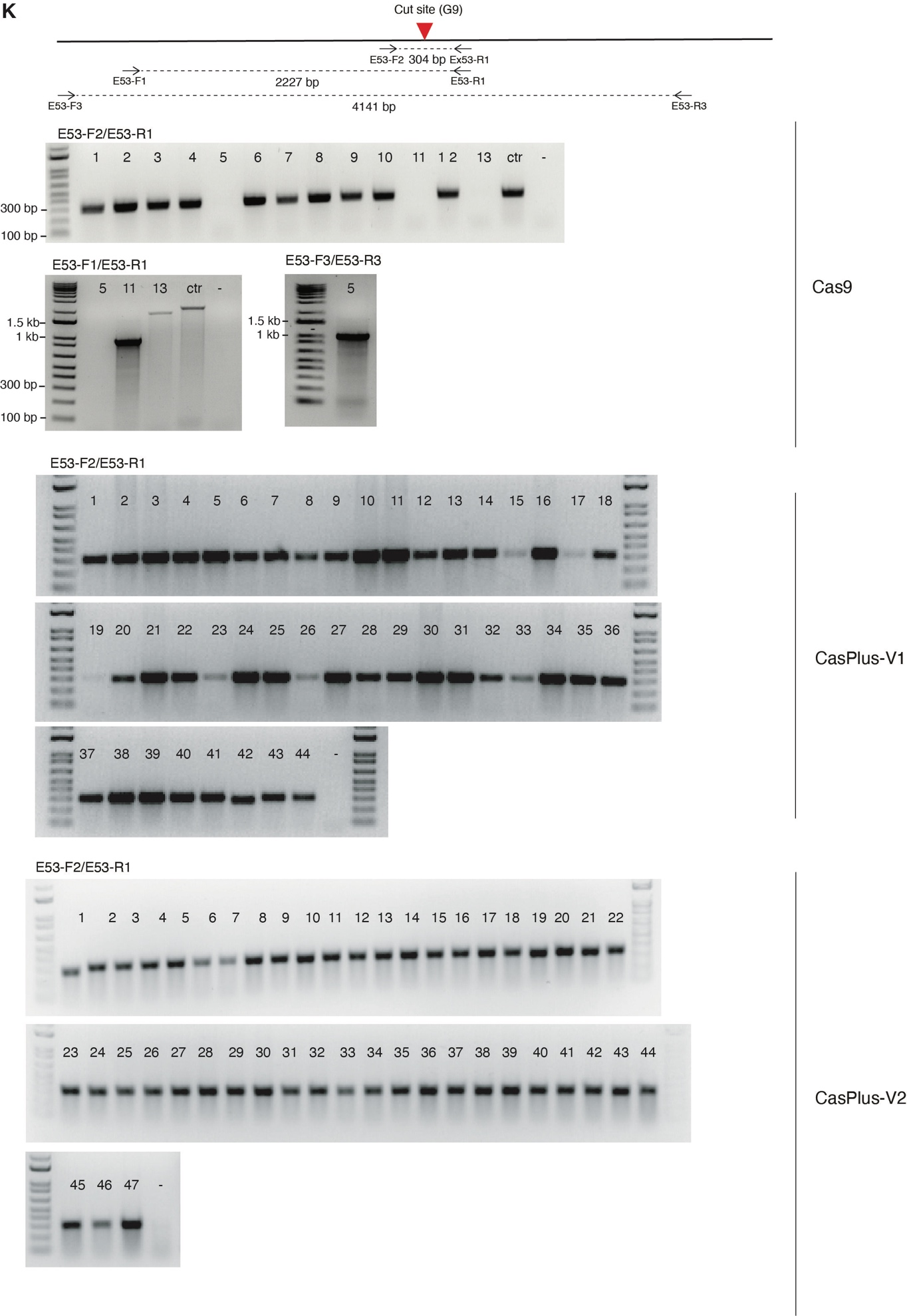

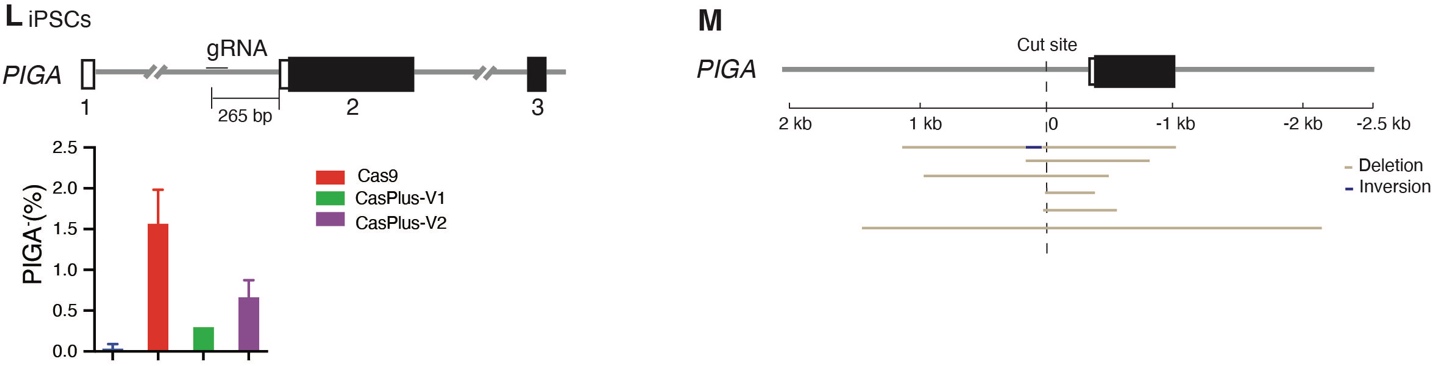
**Appendix Figure 4. CasPlus editing represses large deletions in iPSCs.** (**A**) Representative gel image showing the RT-PCR products amplified from DMD-WT iCMs, untreated or edited DMD-del52 iCMs pools, or DMD-del52 single clone containing 1-bp insertions. Lower bands were purified and sequenced. (**B**) Sanger sequencing results illustrating the skipping of the whole exon 51 using gRNA G10 and exon 53 using gRNA G9 in Cas9-edited DMD-del52 iCMs. (**C**) Schematic illustrating the locations of primer sets used to amplify the target region on *DMD* exons 51 (top) and 53 (bottom). (**D-E**) Representative gel images showing the PCR products amplified from DMD-WT iPSCs, untreated DMD-del52 iPSCs or DMD-del52 iPSCs with edits at *DMD* exon 51 (**D**) or 53 (**E)**. (**F-G**) Representative gel images showing the PCR products amplified from DMD-WT iCMs, untreated DMD-del52 iCMs or DMD-del52 iCMs with edits at *DMD* exon 51 (**F**) or 53 (**G)**. (**H**) Frequencies of indels in DMD-del52 iPSCs edited by Cas9, CasPlus-V1, or -V2 editing with gRNA G10 (top) or G9 (bottom). Frequencies of indels were calculated based on the PacBio sequencing results. (**I**) Raw PCR images of genotyping of single clones isolated from bulk iPSCs edited by Cas9, CasPlus-V1 or CasPlus-V2 with gRNA G10. Top panel indicates the locations of each primer set and the size of each amplicon. All clones were first genotyped using E51-F2/E51-R1 primers and further amplified using E51-F2/E51-R2 if the first round of PCR failed. Clones 3 and 8 which were isolated from the Cas9-edited cell pool were further genotyped using Ex43-F/Ex43-R and either E51-Pacbio-F/E51-R3 or E51-F3/E51-Pacbio-R primers. Failure of the PCR reaction indicates that one or both of the primer binding sites have been lost due to deletions. Primers for exon 43 here were used as a positive control for the PCR reactions. Green arrowheads, expected bands for exon 43; Red arrowheads, expected bands for exon 51. (**J**) Raw PCR images of genotyping of single clones isolated from bulk iPSCs edited by Cas9, CasPlus-V1 or CasPlus-V2 with gRNA G9. Top panel indicates the locations of each primer set and the size of each amplicon. All clones were genotyped first using E53-F2/E53-R1 primers, and then with E53-F1/E53-R1 and E53-F3/E53-R3 if the former round of PCR failed. **(K)** Sequences of clone 4 isolated from Cas9-edited iPSCs using gRNA G10, clone 5, 11 and 13 isolated from Cas9-edited iPSCs using gRNA G9. (**L**) Top schematic demonstrated the location of the gRNA against gene *PIGA*. The graph showed the frequency of PIGA^-^ populations in iPS cells with Cas9, CasPlus-V1 or CasPlus-V2 editing. (**M)** Large deletions detected in single clones isolated from PIGA^-^ population.

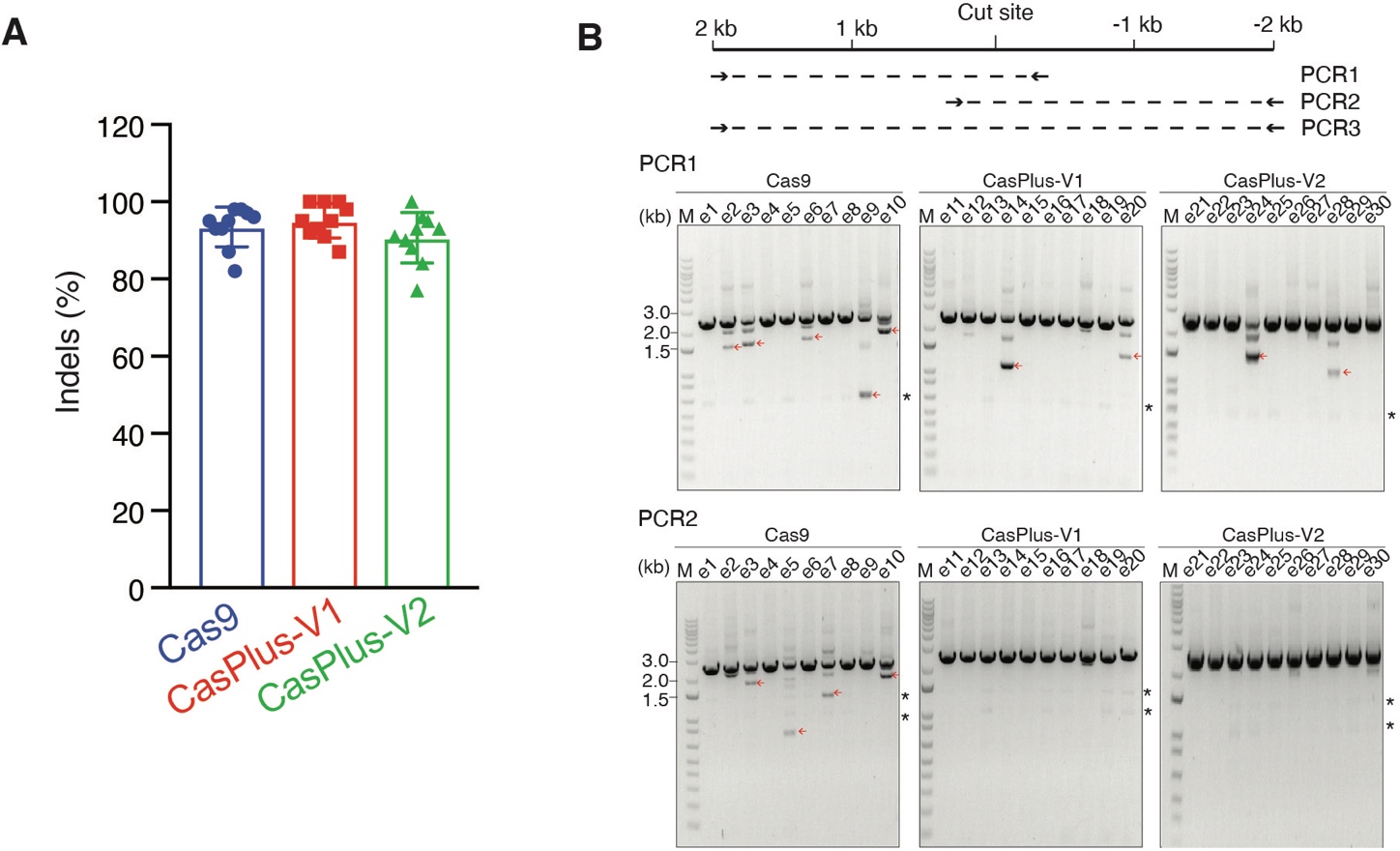

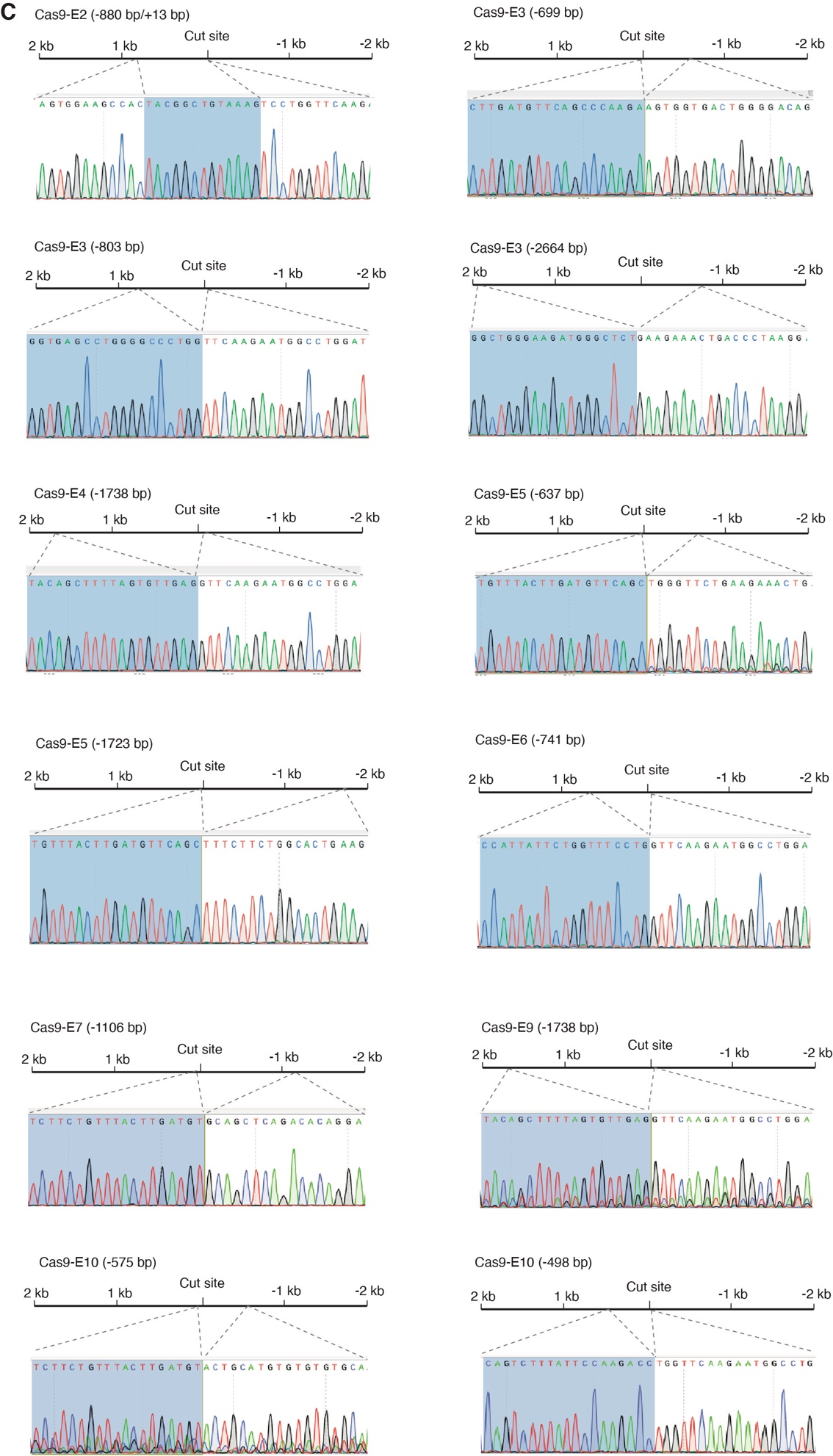

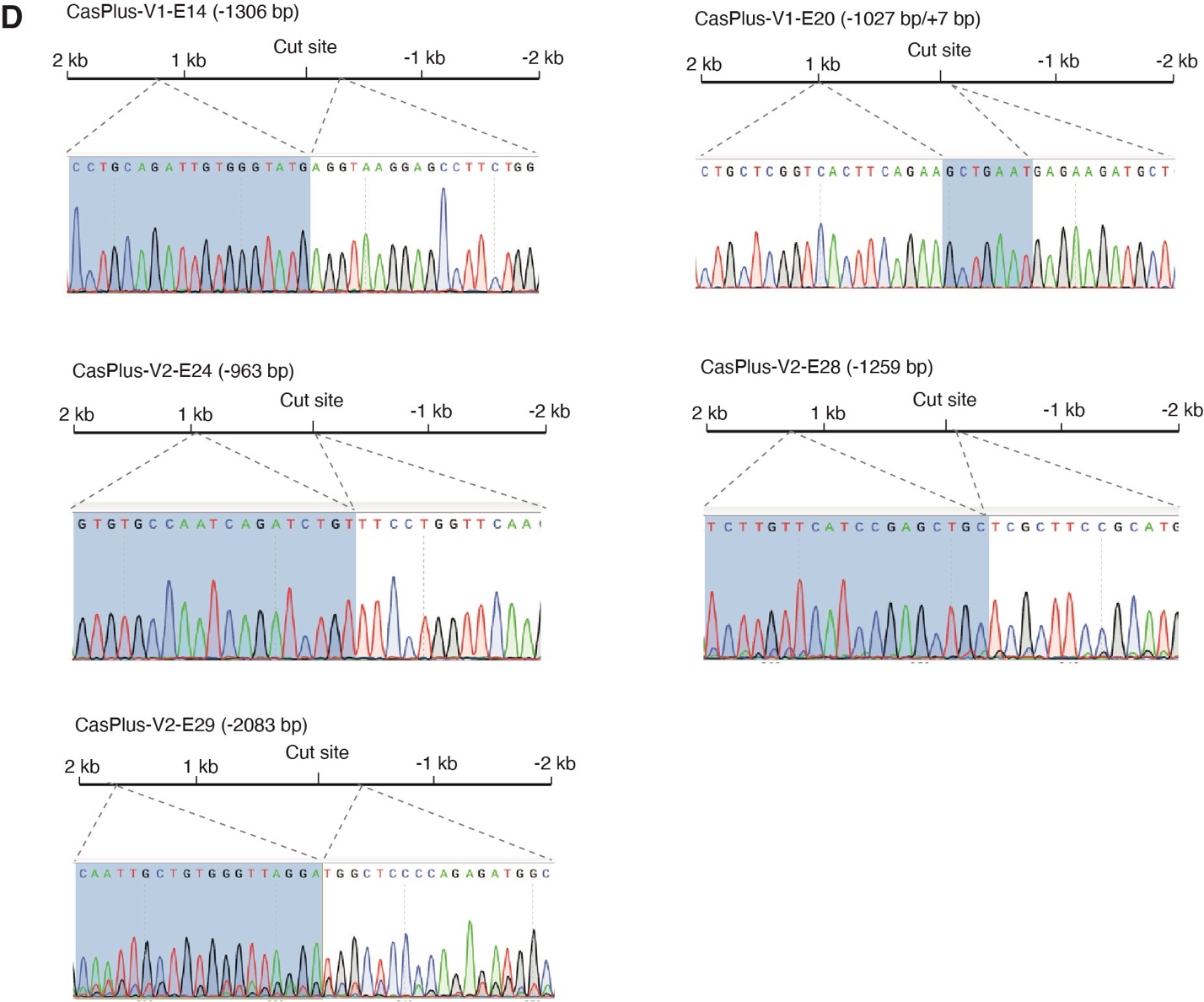

**Appendix Figure 5. CasPlus editing represses large deletions in mouse germline.** (**A**) Summarized editing efficiencies of 10 embryos in each group edited by Cas9, CasPlus-V1 or CasPlus-V2. (**B**) Gel images showing the PCR1 and PCR2 results in mouse embryos. Lower molecular mass bands highlighted with arrowheads were sequenced. Asterisk, non-specific band. (**C-D**) Sanger sequencing files described in Fig **5C**.

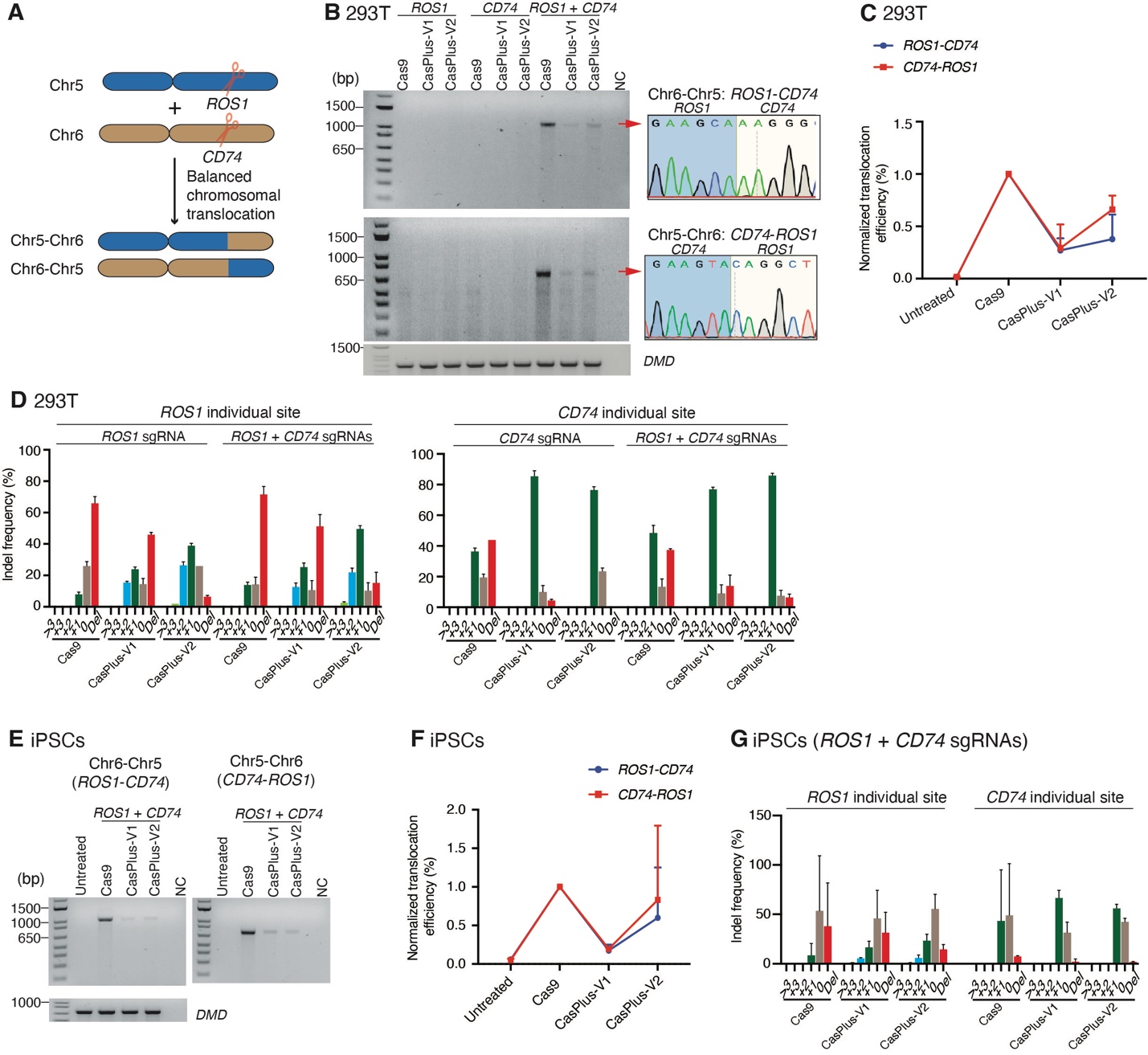

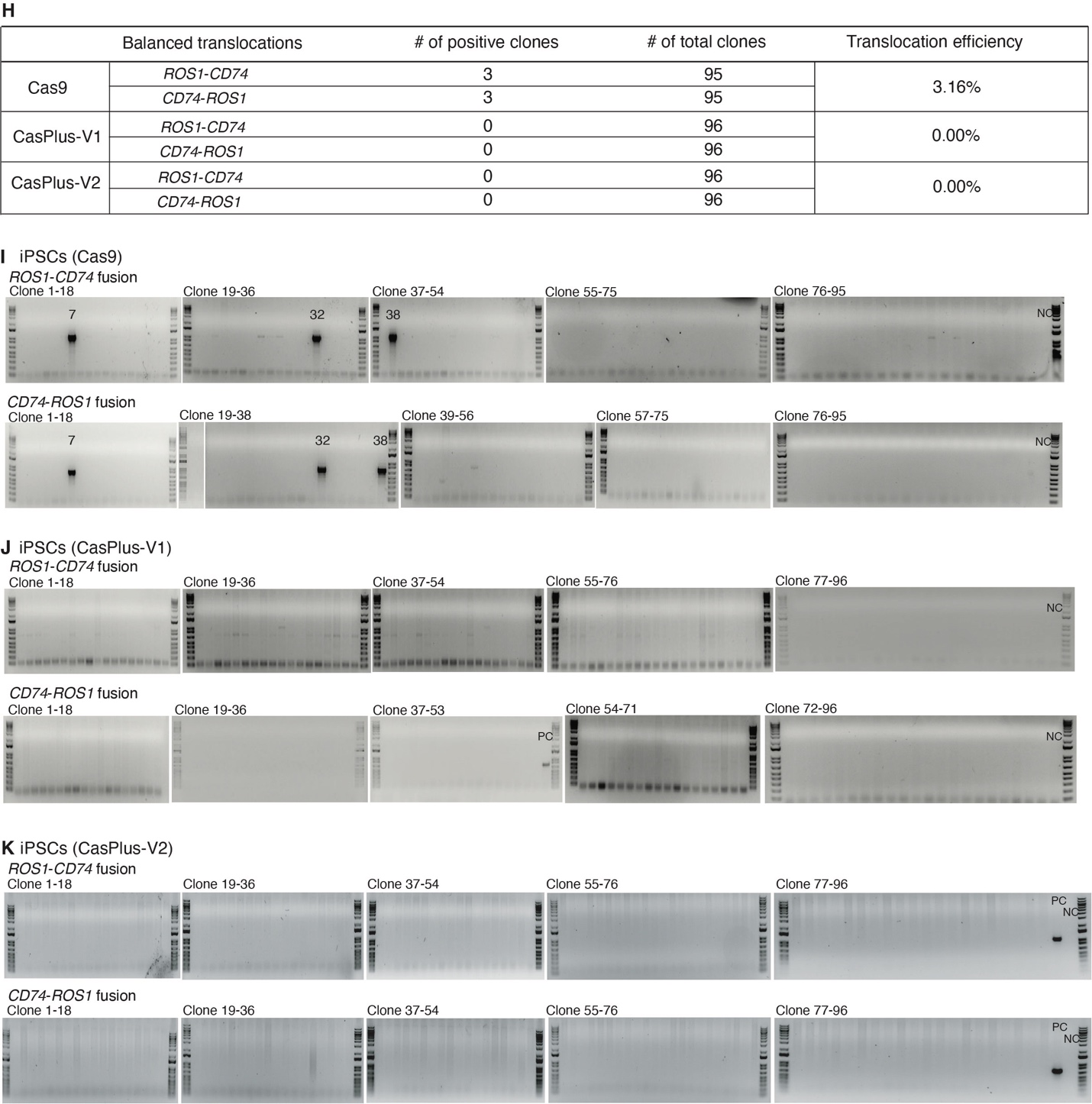

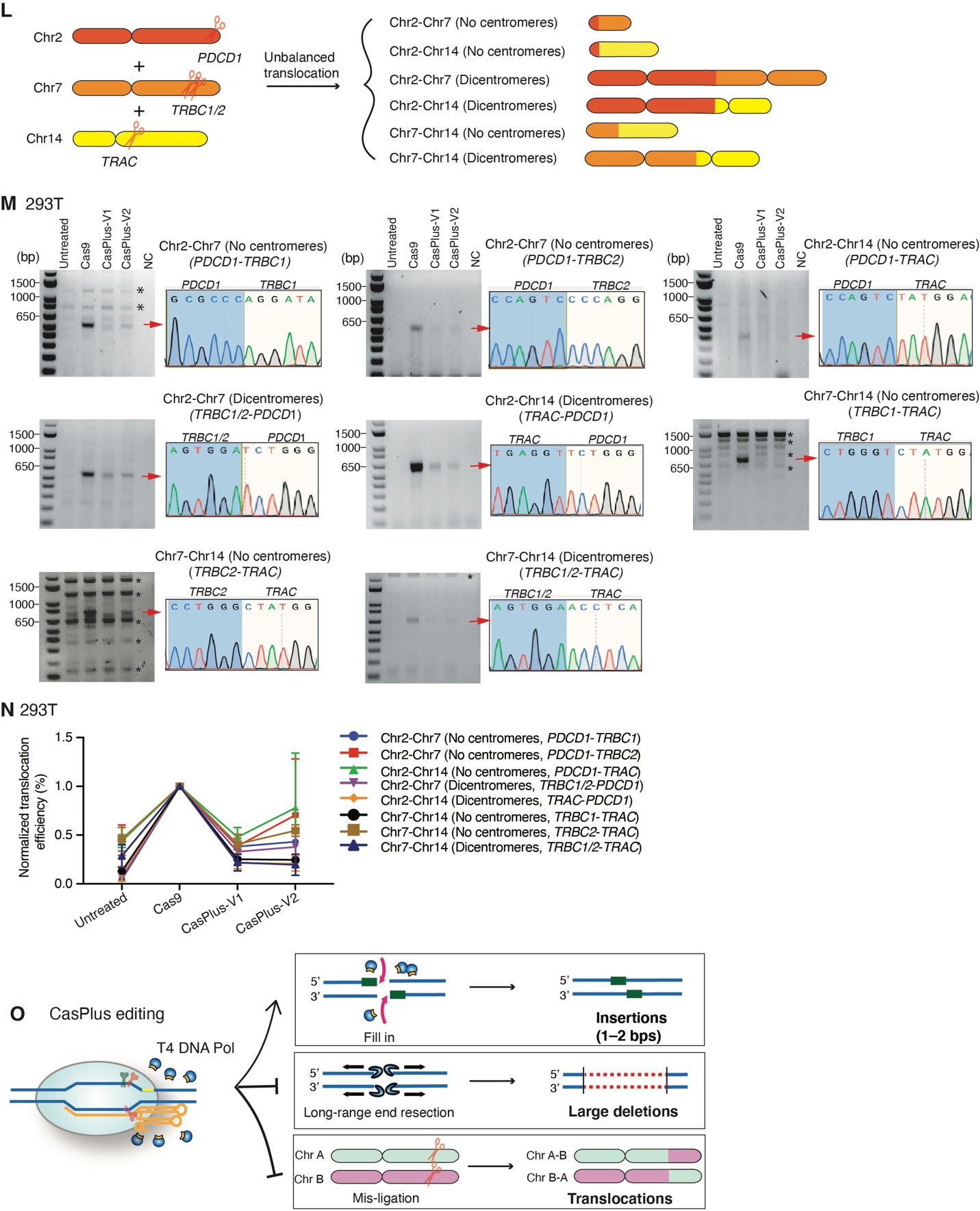

**Appendix Figure 6. CasPlus editing represses chromosomal translocations during multiplex gene editing.** (**A**) Schematic illustrating the formation of balanced chromosomal translocations between genes *ROS1* and *CD74*. (**B**) Representative gel images showing *ROS1-CD74* and *CD74-ROS1* translocations in HEK293T cells during Cas9, CasPlus-V1, or CasPlus-V2 editing (see methods). Bands with expected size (red arrowheads) were purified, TA-cloned and sequenced. One of the representive TA cloning results was shown on the right side of each image. *DMD* is a control for intensity normalization. NC, negative control. (**C**) Normalized quantification of data in **B**. Band intensity obtained from Cas9-edited cells is set as 1. Values and error bars reflect mean ± SEM of n=3 replicates. (**D**) Frequencies of indels at *ROS1* and *CD74* individual sites in HEK293T cells during Cas9, CasPlus-V1, or CasPlus-V2 editing. Frequencies were calculated based on ICE analysis on Sanger sequencing files. Values and error bars reflect mean ± SEM of n=3 replicates. (**E)** Representative gel images demonstrating the *ROS1-CD74* and *CD74-ROS1* translocations in iPSCs during Cas9, CasPlus-V1, or CasPlus-V2 editing. (**F**) Normalized quantification of data in **E**. (**G**) Frequency of indels at *ROS1* and *CD74* individual sites in iPSCs during Cas9, CasPlus-V1, or CasPlus-V2 editing. Frequency was calculated based on ICE analysis on Sanger sequencing files. (**H**) Table of number of total single clones tested and clones containing *ROS1*-*CD74* and *CD74*-*ROS1* translocations in each group. Single clones were isolated from pool of Cas9-, CasPlus-V1- or -V2- edited iPSCs and genotyped using primers specifically binding to the breakpoint junction region of *ROS1*-*CD74* and *CD74*-*ROS1* fused chromosomes. (**I**-**K**) Raw PCR images showing the *ROS1*-*CD74* and *CD74*-*ROS1* translocations occurred in single clones isolated from Cas9- (**I**), CasPlus-V1- (**J**) or -V2- (**K**) edited iPSCs. PC, positive control; NC, negative control. (**L**) Schematic illustrating 6 types of unbalanced inter-chromosomal translocations among genes *PDCD1*, *TRBC1/2*, and *TRAC*. (**M**) Gel images demonstrating the unbalanced inter-chromosomal translocations in HEK293T cells during Cas9, CasPlus-V1, and CasPlus-V2 editing. Expected bands (read arrowheads) were purified, TA-cloned and sequenced. Asterisk, non-specific bands. (**N**) Quantitation of data in **M**. (**O**) Features of CasPlus editing. CasPlus editing utilizes T4 DNA polymerase to fill in the Cas9-created overhangs, thereby biasing insertions over small or large deletions. CasPlus editing can also repress chromosomal translocations that potentially occur between either on-target and off-target site during Cas9-mediated single site editing or different on-target sites during multiplex gene editing.

Supplementary table 1. Information of the pegRNAs used for prime editing

| PegRNA ID | Guide RNA | RTT | PBS | Editing efficiency* |
| --- | --- | --- | --- | --- |
| TS5-R11P13 | ACCTTCACTGGCTGAGTGGC | AAAACCAGCCA | ACTCAGCCAGTGA | 12.5 |
| TS5-R11P17 | ACCTTCACTGGCTGAGTGGC | AAAACCAGCCA | ACTCAGCCAGTGAAGGT | 0 |
| TS5-R18P13 | ACCTTCACTGGCTGAGTGGC | ACAAGGAAAAACCAGCCA | ACTCAGCCAGTGA | 13.5 |
| TS5-R18P17 | ACCTTCACTGGCTGAGTGGC | ACAAGGAAAAACCAGCCA | ACTCAGCCAGTGAAGGT | 0 |
| TS10-R11P13 | TCATCTCGTTGATATCCTCA | GTGACCTTGAG | GGATATCAACGAG | 5.5 |
| TS10-R11P10 | TCATCTCGTTGATATCCTCA | GTGACCTTGAG | GGATATCAAC | 0 |
| TS10-R9P13 | TCATCTCGTTGATATCCTCA | GACCTTGAG | GGATATCAACGAG | 5.5 |
| TS10-R9P10 | TCATCTCGTTGATATCCTCA | GACCTTGAG | GGATATCAAC | 0 |
| TS5-nicking | AGCAGTTCAAGCTAAACAAC | N/A | N/A | N/A |
| TS10-nicking | CTTGGACAGAACTTACCGAC | N/A | N/A | N/A |

*Editing efficiencies were calculated based on ICE analysis on Sanger sequencing files. Date was shown as average of two replicates. N/A, non-available.

Supplemenatary table 2. Information on WGS samples.

| Sample ID | Cas9 (without G10) | Cas9 (containing G10) | T4-WT | T4-D219A |
| --- | --- | --- | --- | --- |
| S1 | + | - | - | - |
| S2 | - | - | + | - |
| S3 | - | - | - | + |
| S4 | - | + | - | - |
| S5 | - | + | + | - |
| S6 | - | + | - | + |

**Supplementary table 3.** Quantity and distribution of total indels and SNPs detected by WGS.

| Sample ID | Total indels | Intergenic | Exonic | Intronic |
| --- | --- | --- | --- | --- |
| S1 | 1313267 | 488466 | 40028 | 784773 |
| S2 | 1308899 | 485670 | 39845 | 783384 |
| S3 | 1302111 | 482497 | 39654 | 779960 |
| S4 | 1325423 | 495575 | 40560 | 789288 |
| S5 | 1308961 | 486255 | 39816 | 782890 |
| S6 | 1307545 | 485655 | 39930 | 781960 |
| Sample ID | Total SNPs | Intergenic | Exonic | Intronic |
| S1 | 3353114 | 1489327 | 146469 | 1717318 |
| S2 | 3353218 | 1484638 | 146152 | 1722428 |
| S3 | 3351872 | 1479051 | 145997 | 1726824 |
| S4 | 3353715 | 1505470 | 147146 | 1701099 |
| S5 | 3352302 | 1486992 | 146222 | 1719088 |
| S6 | 3351200 | 1483156 | 146098 | 1721946 |

**Supplementary table 4**. Summary of predicted off-target sites.

| No. of mismatches | PAM | | Off-targets in WGS samples | | | | | |
| --- | --- | --- | --- | --- | --- | --- | --- | --- |
|  | NGG | NAG | S1 | S2 | S3 | S4 | S5 | S6 |
| 0 | 1 | 0 | 0 | 0 | 0 | 1 | 1 | 1 |
| 1 | 0 | 0 | 0 | 0 | 0 | 0 | 0 | 0 |
| 2 | 0 | 1 | 0 | 0 | 0 | 0 | 0 | 0 |
| 3 | 4 | 9 | 1 | 1 | 1 | 1 | 1 | 1 |
| 4 | 84 | 141 | 0 | 0 | 0 | 0 | 0 | 0 |

Supplementary table 5. Large deletions generated by Cas9 and CasPlus editing using G10 or G9 in male DMD-del52 cells.

| Deletion size (bp) | No. of reads | | | | | | | |
| --- | --- | --- | --- | --- | --- | --- | --- | --- |
|  | G10 | | | | G9 | | | |
|  | Untreated | Cas9 | CasPlus-V1 | CasPlus-V2 | Untreated | Cas9 | CasPlus-V1 | CasPlus-V2 |
| 201–500 | 0 | 19 | 0 | 0 | 0 | 11 | 0 | 2 |
| 501–1000 | 0 | 47 | 4 | 1 | 0 | 5 | 0 | 2 |
| 1001–1500 | 0 | 68 | 4 | 0 | 0 | 22 | 0 | 3 |
| 1501–2000 | 0 | 196 | 0 | 1 | 1 | 6 | 0 | 1 |
| 2001–2500 | 2 | 0 | 0 | 0 | 2 | 49 | 0 | 1 |
| 2501–3000 | 49 | 66 | 41 | 61 | 394 | 197 | 190 | 205 |
| 3001–3500 | 2 | 2 | 1 | 3 | 1 | 568 | 0 | 0 |
| 3501–4000 | 2 | 0 | 3 | 8 | 4 | 0 | 0 | 5 |
| 4001–4500 | 1 | 1 | 1 | 15 | 5 | 5 | 0 | 4 |
| 4501–5000 | 3 | 2 | 1 | 5 | 8 | 1 | 1 | 11 |
| 5001–5500 | NA | NA | NA | NA | 6 | 0 | 1 | 7 |
| Total* | 2902 | 1742 | 1699 | 2700 | 2988 | 1767 | 2029 | 1385 |

*Only those circular consensus sequencing (CCS) reads containing both the 5′ and 3′ index sequences were analyzed.

Supplementary table 6. Summary of the primers used in this paper.

| PCR, RT-PCR primer | Sequences | Purpose of primer |
| --- | --- | --- |
| tdTomato-Seq-F2 | ATGGTGAGCAAGGGCGAGGAGGTC | HTS primers for tdTomato-sgRNA |
| tdTomato-Seq-R2 | TTGAAGCCCTCGGGGAAGGACA |  |
| MYBPC3-Seq-F3 | CTTGGTTCCATGTTTGTTTCC | HTS primers for TS1 and TS16 |
| MYBPC3-Seq-R3 | CGTGCCTCGCCCTGTAAGTT |  |
| DHPS-Seq-F1 | GTGGTTGTCTCCCGTGCCAG | HTS primers for TS2 |
| DHPS-Seq-F2 | ATCTGTTCACCACATGGCCA |  |
| CFTR-Seq-F2 | CCTGAGCGTGATTTGATAATGACCTAATAATGATGGG | HTS primers for TS3 and CFTR-F508del correction |
| CFTR-Seq-R2 | CCCATAGAGGAAACATAAATATATGTAGACTAACCG |  |
| LMNA-Ex3-Seq-F1 | ATGTCCTGACCCCTGCAGGCAT | HTS primers for TS4 |
| LMNA-Ex3-Seq-R1 | GCCCTCTGGCTTGCAAAGCA |  |
| DMD-Ex49-Seq-F1 | GGAGTCTCCAAGGGTATATT | HTS primers for TS5, TS8, TS18, and TS6 |
| DMD-Ex49-Seq-R2 | GTATGGCCAGTATTTCCTTACAAGTTATTTC |  |
| PCSK9-Seq-F2 | TAGGTCTCCTCGCCAGGACAG | HTS primers for TS7 |
| PCSK9-Seq-R2 | AAACAGCACCGCACCGTCCCGGCTGCG |  |
| DMD-Ex53-Seq-F2 | TCCTAGCCATAACACAATG | HTS primers for TS9 (or G9) |
| DMD-Ex53-Seq-R1 | AGGGACCCTCCTTCCATGAC |  |
| DMD-Ex51-Seq-F2 | ATCTCCAAACTAGAAATGCC | HTS primers for TS10 (or G10) and TS25 |
| DMD-Ex51-Seq-R2 | CTGGTGGGAAATGGTCTAGGAG |  |
| DMD-Ex51-Seq-F3 | GTTAGAAAAAGATATATAATGTCATGAATAAGAGTTTGGC | HTS primers for TS11 and TS24 |
| DMD-Ex51-Seq-R1 | CGGTAAGTTCTGTCCAAGCC |  |
| LMNA-Ex2-Seq-F2 | TAGCTGCTCAGGCTCGGCTGAA | HTS primers for TS12 and TS26 |
| LMNA-Ex2-Seq-R2 | ACCTCAGTGGGTAAAATGAC |  |
| PCSK9-Seq-F3 | AAGGCTCAAGGCGCCGCCGGCGTGGA | HTS primers for TS13 |
| PCSK9-Seq-R3 | CCTCCCATCCCTACACCCGCA |  |
| MYBPC3-Seq-F1 | TACCAAGTCCTGTCACCAACAG | HTS primers for TS14, TS20 and TS21 |
| MYBPC3-Seq-R1 | TGAGTACCATGGCCCTGCCCA |  |
| DMD-Ex43-Seq-F1 | GGTCGGATTGACATTATTCATAGC | HTS primers for TS15 and TS17 |
| DMD-Ex43-Seq-R1 | CAATGAGAGTGATACTTCTTTTTCCCTGTC |  |
| HEXA-Seq-F1 | CTGCCATTTGACCTTTTATAACAGATTCAGCC | HTS primers for TS19 |
| HEXA-Seq-R1 | TTCATTTACTAACTGGAAATGTCTGGCCC |  |
| DMD-Ex45-Seq-F1 | GCCAGTACAACTGCATGTGG | HTS primers for TS22 |
| DMD-Ex45-Seq-R1 | CTCTTTTTTCTGTCTGACAGCT |  |
| MYBPC3-Seq-F2 | CCAACAGGCATCACCTATGAG | HTS primers for TS23 |
| MYBPC3-Seq-R2 | CAAACATGGAACCAAGAGTGAG |  |
| LMNA-Ex11-Lb1-Seq-F1 | TAATAGTGCATGCCTGCTGC | HTS primers for Lb1 |
| LMNA-Ex11-Lb1-Seq-R1 | GAGACAAGGCTCAGGCGGGA |  |
| DMD-Ex52-F1 | TTCTTACTCAAGGCATTCAGAC | Primers for genotyping iPSCs with *DMD* exon 52 deletion |
| DMD-Ex52-R1 | GGTCACCACACCCATCAAT |  |
| DMD-Ex51-Seq-F4 | CAAACTAGAAATGCCATCTTC | Primers to detect indels induced by Ex51-G2 |
| DMD-Ex51-Seq-R3 | CCTTGATTATACTTAGGCTG |  |
| DMD-Ex53-F1 | GGGAAATCAGGCTGATGGGT | Primers to detect indels induced by Ex53-G2, -G3, -G4 and -G5 |
| DMD-Ex53-R1 | GTCTACTGTTCATTTCAGC |  |
| DMD-RT-F1 | CTAAGGAAACTGCCATCTCC | RT-PCR primers to detect small indels induced by G10 in iCMs |
| DMD-RT-R1 | TAAGACCTGCTCAGCTTCTTC |  |
| DMD-RT-F2 | GAGGTACCTGCTCTGGCAGA | RT-PCR primers to detect small indels induced by G9 in iCMs |
| DMD-RT-R2 | GTATAGGGACCCTCCTTCCA |  |
| DMD-RT-E49-F3 | GAAACTGAAATAGCAGTTCAAGCTAAACAACCGG | RT-PCR primers to detect *DMD* exon 51 or exon 53 skipping |
| DMD-RT-E54-R3 | TTCTCCAAGAGGCATTGATATTCT |  |
| CFTR-F1 | GCTGTTACTAACCTTTCCCATTC | Primers for genotyping CFTR-F508del HEK293T cells |
| CFTR-R1 | ACCATTGAGGACGTTTGTCT |  |
| E51-F1 | CTGGTTATATTATGCTCTTGAGTCTG | Primers to detect large deletions induced by G10 |
| E51-R1 | CTGGTGGGAAATGGTCTAGGAG |  |
| E51-F2 | ATCTCCAAACTAGAAATGCC | Primers to detect large deletions induced by G10 |
| E51-R2 | CCATATCCATAACTATATTGTGAATGTG |  |
| M13-Pacbio-Ex51-F | GCAGTCGAACATGTAGCTGACTCAGGTCACctgactttcttctttccttcttcctttctgtc | Primers for PacBio sequencing on DMD-Ex51 |
| M13-Pacbio-Ex51-R | TGGATCACTTGTGCAAGCATCACATCGTAGgatggagtctcactctgtcactcagtc |  |
| E53-F1 | CTTTATTCTGGGAGGCTGACC | Primers to detect large deletions induced by G9 |
| E53-R1 | AGGGACCCTCCTTCCATGAC |  |
| E53-F2 | TCCTAGCCATAACACAATG | Primers to detect large deletions induced by G9 |
| E53-R2 | GGAAAATGACCATTGGAGATTCC |  |
| M13-Pacbio-Ex53-F | GCAGTCGAACATGTAGCTGACTCAGGTCACtgtgtgccacttactgagttttaattattctgg | Primers for PacBio sequencing on DMD-Ex53 |
| M13-Pacbio-Ex53-R | TGGATCACTTGTGCAAGCATCACATCGTAGgaaatggatgaaaaaattacagctttggcc |  |
| Mybpc3-F1 | TTCTCTGAGGCCCCAAGCTT | Primers for PCR2 |
| Mybpc3-R3 | gacagttccactctgggaagtg |  |
| Mybpc3-F3 | ccccctcaaaagataacacagggg | Primers for PCR1 |
| Mybpc3-R1 | ccaagatccactgaacatgtcag |  |
| Mybpc3-F3 | ccccctcaaaagataacacagggg | Primers for PCR3 |
| Mybpc3-R3 | gacagttccactctgggaagtg |  |
| ROS1-CD74-F | ATCAGCCACCCCCTTAATTCCT | Primers to detect *ROS1*-*CD74* fused chromosome |
| ROS1-CD74-R | CCATGTGTCCAATTCACCTC |  |
| CD74-ROS1-F | TTGTCTAAGGGTTCCTAAGGGC | Primers to detect *CD74*-*ROS1* fused chromosome |
| CD74-ROS1-R | GACTAAGGAGCATCAGGTAAGC |  |
| ROS1-F | CCAGCCTTATGACCACTCCT | Primers to detect indels at *ROS1* individual site |
| ROS1-R | GCAAACACAGGGCCAAAGAC |  |
| CD74-F | GCTGCAGAGTATGTGGGGTT | Primers to detect indels at *CD74* individual site |
| CD74-R | AGCAATGGGCACCTTGGTAA |  |
| PDCD1-R1 | CAGCACTGCCTCTGTCACTCTC | Primers to detect indels at *PDCD1* individual site |
| PDCD1-F1 | CGAGTGAGGACCAAGGATGC |  |
| TRBC1/2-F | GCAGAGATCTCCCACACCCAAAAG | Primers to detect indels at *TRBC1* individual site |
| TRBC1-R1 | CAGACCAGCTTCCTCTGACAGC |  |
| TRBC1/2-F1 | GCAGAGATCTCCCACACCCAAAAG | Primers to detect indels at *TRBC2* individual site |
| TRBC2-R1 | TCCCTCATGCTGCTCCAGTG |  |
| TRAC-F2 | TTCAGGTTTCCTTGAGTGGC | Primers to detect indels at *TRAC* individual site |
| TRAC-R2 | TGCCAGAACAAGGCTCACTG |  |
| PDCD1-F1 | CGAGTGAGGACCAAGGATGC | Primers to amplify breakpoint region of balanced translocation of *PDCD1-TRAC* |
| TRAC-R1 | CTCCTGCCACCTTCTCTTCA |  |
| PDCD1-F1 | CGAGTGAGGACCAAGGATGC | Primers to amplify breakpoint region of balanced translocation of *PDCD1-TRBC1* |
| TRBC1-R1 | CAGACCAGCTTCCTCTGACAGC |  |
| PDCD1-F1 | CGAGTGAGGACCAAGGATGC | Primers to amplify breakpoint region of balanced translocation of *PDCD1-TRBC2* |
| TRBC2-R1 | TCCCTCATGCTGCTCCAGTG |  |
| TRBC1/2-F | GCAGAGATCTCCCACACCCAAAAG | Primers to amplify breakpoint region of balanced translocation of *TRBC1/2-PDCD1* |
| PDCD1-R1 | CAGCACTGCCTCTGTCACTCTC |  |
| TRBC1/2-F1 | GCAGAGATCTCCCACACCCAAAAG | Primers to amplify breakpoint region of balanced translocation of *TRBC1/2*-*TRAC* |
| TRAC-R2 | TGCCAGAACAAGGCTCACTG |  |
| TRAC-F2 | TTCAGGTTTCCTTGAGTGGC | Primers to amplify breakpoint region of balanced translocation of *TRAC*-*TRBC1* |
| TRBC1-R1 | CAGACCAGCTTCCTCTGACAGC |  |
| TRAC-F2 | TTCAGGTTTCCTTGAGTGGC | Primers to amplify breakpoint region of balanced translocation of *TRAC*-*TRBC2* |
| TRBC2-R1 | TCCCTCATGCTGCTCCAGTG |  |
| PDCD1-R3 | ACAGTTTCCCTTCCGCTCAC | Primers to amplify breakpoint region of unbalanced translocation of *PDCD1*-*TRBC1* (No centromeres) |
| TRBC1-R2 | TTTGCAGTTGGCACAGACTC |  |
| PDCD1-R3 | ACAGTTTCCCTTCCGCTCAC | Primers to amplify breakpoint region of unbalanced translocation of *PDCD1*-*TRBC2* (No centromeres) |
| TRBC2-R2 | AGTCAGCCTCCTTTGATAGC |  |
| PDCD1-R3 | ACAGTTTCCCTTCCGCTCAC | Primers to amplify breakpoint region of unbalanced translocation of *PDCD1*-*TRAC* (No centromeres) |
| TRAC-R1 | CTCCTGCCACCTTCTCTTCA |  |
| TRBC1/2-F1 | GCAGAGATCTCCCACACCCAAAAG | Primers to amplify breakpoint region of unbalanced translocation of *TRBC1/2*-*PDCD1* (Dicentromeres) |
| PDCD1-F2 | AAAGCCACACAGCTCAGGGTAAG |  |
| TRAC-F3 | GTCTGTCTGCCTATTCACCG | Primers to amplify breakpoint region of unbalanced translocation of *TRAC*-*PDCD1* (Dicentromeres) |
| PDCD1-F1 | CGAGTGAGGACCAAGGATGC |  |
| TRBC1-R1 | CAGACCAGCTTCCTCTGACAGC | Primers to amplify breakpoint region of unbalanced translocation of *TRBC1*-*TRAC* (No centromeres) |
| TRAC-R1 | CTCCTGCCACCTTCTCTTCA |  |
| TRBC2-R1 | TCCCTCATGCTGCTCCAGTG | Primers to amplify breakpoint region of unbalanced translocation of *TRBC2*-*TRAC* (No centromeres) |
| TRAC-R1 | CTCCTGCCACCTTCTCTTCA |  |
| TRBC1/2-F1 | GCAGAGATCTCCCACACCCAAAAG | Primers to amplify breakpoint region of unbalanced translocation of *TRBC1/2*-*TRAC* (Dicentromeres) |
| TRAC-F1 | TGAAATCATGGCCTCTTGGC |  |
| PDCD1-qPCR-F1 | GTGAAAGGTCCCTCCAGAC | Primers and probes to detect balanced translocation of *PDCD1*-*TRBC1* |
| TRBC1-qPCR-R1 | GCTGAGAATCTGGGAGAAGAAT |  |
| TRBC1-probe | ACCATGAAGGAGAATTGGGCACCT |  |
| PDCD1-qPCR-F1 | GTGAAAGGTCCCTCCAGAC | Primers and probes to detect balanced translocation of *PDCD1*-*TRBC2* |
| TRBC2-qPCR-R1 | GCTTTGCTGACCCTGTGA |  |
| TRBC2-probe | AGAAAGCCAGAGTGGACAAGGTGG |  |
| PDCD1-qPCR-F1 | GTGAAAGGTCCCTCCAGAC | Primers and probes to detect balanced translocation of *PDCD1*-*TRAC* |
| TRAC-qPCR-R1 | CATTCCTGAAGCAAGGAAACAG |  |
| TRAC-Probe | TGCAAACGCCTTCAACAACAGCAT |  |
| TRBC1/2-qPCR-F1 | GCAGAGATCTCCCACACCCAAAAG | Primers and probes to detect balanced translocation of *TRBC1/2*-*PDCD1* |
| PDCD1-qPCR-R2 | CAGCACTGCCTCTGTCACTCTC |  |
| PDCD1-Probe | AGGTGAGCGGAAGGGAAACTGTC |  |
| TRBC1/2-qPCR-F1 | GCAGAGATCTCCCACACCCAAAAG | Primers and probes to detect balanced translocation of *TRBC1/2*-*TRAC* |
| TRAC-qPCR-R2 | CTCCTGCCACCTTCTCTTCA |  |
| TRAC-Probe | TGCAAACGCCTTCAACAACAGCAT |  |
| TRAC-qPCR-F1 | GGGCAAAGAGGGAAATGAGA | Primers and probes to detect balanced translocation of *TRAC*-*TRBC1* |
| TRBC1-qPCR-R1 | GCTGAGAATCTGGGAGAAGAAT |  |
| TRBC1-probe | ACCATGAAGGAGAATTGGGCACCT |  |
| TRAC-qPCR-F1 | GGGCAAAGAGGGAAATGAGA | Primers and probes to detect balanced translocation of *TRAC*-*TRBC2* |
| TRBC2-qPCR-R1 | GCTTTGCTGACCCTGTGA |  |
| TRBC2-probe | AGAAAGCCAGAGTGGACAAGGTGG |  |
| TRAC-qPCR-F1 | GGGCAAAGAGGGAAATGAGA | Primers and probes to detect balanced translocation of *TRAC*-*PDCD1* |
| PDCD1-qPCR-R1 | CACTGCCTCTGTCACTCTC |  |
| PDCD1-probe | AGGTGAGCGGAAGGGAAACTGTC |  |
| DMD_e52-i52-F1 | GCAATCAAGAGGCTAGAACAATCA | Primers and probes to detect taqman reference gene *DMD* copies. |
| DMD_e52-i52-R1 | CTTTGTGTGTCCCATGCTTGTTA |  |
| DMD_e52-i52-probe | TACGGATCGAAGTAAGTTT |  |

Supplementary table 7. Summary of the guide RNAs used in this paper.

| ID | Target gene | Sequences (5'-3') | PAM | Purpose of the gRNA |
| --- | --- | --- | --- | --- |
| tdTomato-sgRNA | tdTomato-d151A | CAAGCTGAAGGTGACCAGGG | CGG | CasPlus test |
| TS1 | *MYBPC3* | TTCTTGAACCAGGAAATCTT | GGG | CasPlus test |
| TS2 | *DHPS* | TCCAGGAACAGCTGGGTACC | TGG | CasPlus test |
| TS3 | *CFTR* | ATTAAAGAAAATATCATCTT | TGG | CasPlus test; CFTR-F508del knock-in |
| TS4 | *LMNA* | GCCTGCTTCCTCACAGCTTG | AGG | CasPlus test |
| TS5 | *DMD* | ACCTTCACTGGCTGAGTGGC | TGG | CasPlus test |
| TS6 | *DMD* | CATTACCTTCACTGGCTGAG | TGG | CasPlus test |
| TS7 | *PCSK9* | GGACGAGGACGGCGACTACG | AGG | CasPlus test |
| TS8 | *DMD* | TAAAGGGCAGCATTTGTACA | AGG | CasPlus test |
| TS9 (G9) | *DMD* | TTGAAAGAATTCAGAATCAG | TGG | CasPlus test/Correct DMD-delEx52 |
| TS10 (G10) | *DMD* | TCATCTCGTTGATATCCTCA | AGG | CasPlus test/Correct DMD-delEx52 |
| TS11 | *DMD* | TCCTACTCAGACTGTTACTC | TGG | CasPlus test |
| TS12 | *LMNA* | GGGGCCAGGTGGCCAAGGTG | AGG | CasPlus test |
| TS13 | *PCSK9* | CTGCTGCTCCTGGGTCCCGC | GGG | CasPlus test |
| TS14 | *MYBPC3* | TCCGGGGTAGCCCCAAGGTA | GGG | CasPlus test |
| TS15 | *DMD* | AAAATGTACAAGGACCGACA | AGG | CasPlus test |
| TS16 | *MYBPC3* | ATTTATAGCCCAAGATTTCC | TGG | CasPlus test |
| TS17 | *DMD* | TATGTGTTACCTACCCTTGT | CGG | CasPlus test |
| TS18 | *DMD* | GGTTGCTTCATTACCTTCAC | TGG | CasPlus test |
| TS19 | *HEXA* | TACCTGAACCGTATATCCTA | TGG | CasPlus test |
| TS20 | *MYBPC3* | CGCCTAAAGTTCCCTACCTT | GGG | CasPlus test |
| TS21 | *MYBPC3* | GCCTAAAGTTCCCTACCTTG | GGG | CasPlus test |
| TS22 | *DMD* | TCCAGGATGGCATTGGGCAG | CGG | CasPlus test |
| TS23 | *MYBPC3* | AGGTAGGGAACTTTAGGCGC | TGG | CasPlus test |
| TS24 | *DMD* | ACCAGAGTAACAGTCTGAGT | AGG | CasPlus test |
| TS25 | *DMD* | TATAAAATCACAGAGGGTGA | TGG | CasPlus test |
| TS26 | *LMNA* | CCTGCAGGGTGGCCTCACCT | TGG | CasPlus test |
| Lb1 | *LMNA* | TGTCTTCCCTCTCCTCCTCCGGG | TTTA | CasPlus test |
| DMD-Ex52-g1 | *DMD* | TAAGGGATATTTGTTCTTAC | AGG | Generating DMD-del52 cells |
| DMD-Ex52-g2 | *DMD* | AGAGGCTAGAACAATCATTA | CGG | Generating DMD-del52 cells |
| Ex51-G2 | *DMD* | CGAGATGATCATCAAGCAGA | AGG | Correcting DMD-delEx52 |
| Ex53-G2 | *DMD* | TACAAGAACACCTTCAGAAC | CGG | Correcting DMD-delEx52 |
| Ex53-G3 | *DMD* | AAGAACACCTTCAGAACCGG | AGG | Correcting DMD-delEx52 |
| Ex53-G4 | *DMD* | ACTGTTGCCTCCGGTTCTGA | AGG | Correcting DMD-delEx52 |
| Ex53-G5 | *DMD* | TTTCATTCAACTGTTGCCTC | CGG | Correcting DMD-delEx52 |
| PIGA sgRNA | *PIGA* | TGTTACACCACAGCATATCA | TGG |  |
| Mybpc3 sgRNA | *Mybpc3* | TTCTTGAACCAGGAAATCTT | GGG | gRNA for embryo-editing |
| *ROS1* sgRNA | *ROS1* | TTAAATTTAGTTGAAGCAC | AGG | Detection of chromosomal translocations beween genes *ROS1* and *CD74* |
| *CD74*  sgRNA | *CD74* | TCCTGAAGTAGAAGGTCAA | AGG |  |
| *PDCD1* sgRNA | *PDCD1* | GGCGCCCTGGCCAGTCGTCT | GGG | Detection of chromosomal translocations beween genes *PDCD1*, *TRBC1/2,* and *TRAC* |
| *TRBC1/2* sgRNA | *TRBC1/2* | GGAGAATGACGAGTGGACCC | AGG |  |
| *TRAC* sgRNA | *TRAC* | TGTGCTAGACATGAGGTCTA | TGG |  |

**Supplementary table 8**. Summary of the synthetic sequences and vector information in this paper.

**tdTomato-d151A**

atggtgagcaagggcgaggaggtcatcaaagagttcatgcgcttcaaggtgcgcatggagggctccatgaacggccacgagttcgagatcgagggcgagggcgagggccgcccctacgagggcacccagaccgccaagctgaaggtgaccagggcggccccctgcccttcgcctgggacatcctgtccccccagttcatgtacggctccaaggcgtacgtgaagcaccccgccgacatccccgattacaagaagctgtccttccccgagggcttcaagtgggagcgcgtgatgaacttcgaggacggcggtctggtgaccgtgacccaggactcctccctgcaggacggcacgctgatctacaaggtgaagatgcgcggcaccaacttcccccccgacggccccgtaatgcagaagaagaccatgggctgggaggcctccaccgagcgcctgtacccccgcgacggcgtgctgaagggcgagatccaccaggccctgaagctgaaggacggcggccactacctggtggagttcaagaccatctacatggccaagaagcccgtgcaactgcccggctactactacgtggacaccaagctggacatcacctcccacaacgaggactacaccatcgtggaacagtacgagcgctccgagggccgccaccacctgttcctggggcatggcaccggcagcaccggcagcggcagctccggcaccgcctcctccgaggacaacaacatggccgtcatcaaagagttcatgcgcttcaaggtgcgcatggagggctccatgaacggccacgagttcgagatcgagggcgagggcgagggccgcccctacgagggcacccagaccgccaagctgaaggtgaccaagggcggccccctgcccttcgcctgggacatcctgtccccccagttcatgtacggctccaaggcgtacgtgaagcaccccgccgacatccccgattacaagaagctgtccttccccgagggcttcaagtgggagcgcgtgatgaacttcgaggacggcggtctggtgaccgtgacccaggactcctccctgcaggacggcacgctgatctacaaggtgaagatgcgcggcaccaacttcccccccgacggccccgtaatgcagaagaagaccatgggctgggaggcctccaccgagcgcctgtacccccgcgacggcgtgctgaagggcgagatccaccaggccctgaagctgaaggacggcggccactacctggtggagttcaagaccatctacatggccaagaagcccgtgcaactgcccggctactactacgtggacaccaagctggacatcacctcccacaacgaggactacaccatcgtggaacagtacgagcgctccgagggccgccaccacctgttcctg

MS2-Linker-NLS**-**T4 DNA polymerase-NLS-HA-T2A-GFP

atggcttcaaactttactcagttcgtgctcgtggacaatggtgggacaggggatgtgacagtggctccttctaatttcgctaatggggtggcagagtggatcagctccaactcacggagccaggcctacaaggtgacatgcagcgtcaggcagtctagtgcccagaagagaaagtataccatcaaggtggaggtccccaaagtggctacccagacagtgggcggagtcgaactgcctgtcgccgcttggaggtcctacctgaacatggagctcactatcccaattttcgctaccaattctgactgtgaactcatcgtgaaggcaatgcaggggctcctcaaagacggtaatcctatcccttccgccatcgccgctaactcaggtatctacagcgctggaggaggtggaagcggaggaggaggaagcggaggaggaggtagcggacctaagaaaaagaggaaggtgAAGGAATTCTACATCAGCATCGAGACCGTGGGTAACAACATCGTGGAAAGATATATTGACGAAAACGGCAAGGAGAGAACCAGAGAGGTGGAATACCTGCCTACAATGTTCCGGCACTGTAAAGAGGAATCCAAGTACAAGGATATCTACGGCAAAAACTGCGCCCCTCAGAAATTCCCCAGCATGAAAGACGCCAGAGATTGGATGAAGAGAATGGAGGATATCGGACTGGAAGCCCTGGGCATGAACGATTTCAAGCTGGCCTACATCTCCGATACATACGGAAGCGAGATCGTGTATGATAGAAAATTCGTGCGGGTGGCCAATTGTGACATTGAGGTGACCGGCGACAAGTTCCCTGATCCCATGAAAGCTGAATATGAGATCGACGCCATTACCCACTACGACAGCATCGACGACAGATTCTACGTGTTCGACCTGCTGAACTCCATGTACGGCAGCGTGTCCAAGTGGGACGCTAAGCTGGCCGCCAAGCTGGACTGCGAGGGCGGCGACGAGGTTCCACAAGAGATCCTGGACCGGGTCATCTACATGCCCTTCGACAACGAGAGGGACATGCTGATGGAATACATCAACCTGTGGGAGCAGAAGCGCCCCGCCATTTTTACAGGCTGGAACATCGAGGGCTTCGACGTGCCTTATATCATGAATAGAGTGAAAATGATCCTGGGAGAACGGAGCATGAAAAGATTCAGCCCTATCGGCAGAGTGAAGAGCAAGCTGATCCAAAACATGTACGGCTCCAAGGAAATCTATAGCATCGATGGCGTGTCCATCCTGGATTACCTGGACCTGTACAAAAAGTTCGCCTTCACCAACCTGCCATCTTTCTCTCTTGAGAGCGTCGCCCAGCACGAGACAAAGAAGGGCAAGCTGCCGTACGACGGTCCTATCAACAAGCTGAGAGAAACAAATCACCAGAGATACATCAGCTACAACATCATCGATGTGGAAAGCGTTCAGGCCATCGATAAAATCAGAGGCTTCATCGACCTGGTGCTGTCTATGTCTTACTACGCCAAGATGCCTTTTAGCGGAGTGATGAGCCCTATCAAGACCTGGGATGCCATCATCTTCAACAGCCTGAAGGGCGAACACAAGGTGATCCCCCAACAGGGCAGCCACGTGAAGCAGAGCTTCCCAGGCGCTTTTGTGTTCGAGCCCAAGCCCATAGCGCGGAGATACATCATGAGCTTTGATCTGACCAGCCTGTACCCCAGCATCATTCGGCAAGTGAACATTTCTCCAGAAACCATCAGAGGCCAGTTTAAGGTGCACCCTATCCACGAGTATATTGCAGGCACCGCTCCTAAACCTAGCGACGAGTACAGCTGCTCTCCTAACGGCTGGATGTACGACAAGCACCAGGAGGGAATCATCCCTAAGGAAATTGCCAAGGTGTTTTTCCAGCGGAAGGACTGGAAGAAAAAAATGTTCGCCGAGGAAATGAACGCCGAGGCCATCAAGAAGATCATCATGAAGGGCGCCGGCAGCTGCTCCACCAAGCCTGAGGTGGAAAGATACGTGAAGTTCAGCGACGATTTCCTGAATGAGCTCAGCAACTACACCGAGTCTGTCCTGAACTCACTGATTGAGGAATGCGAGAAGGCCGCCACCCTGGCTAATACCAACCAGCTGAACCGGAAGATTCTGATCAACAGCCTGTACGGAGCTCTGGGCAATATTCACTTCAGATACTACGATCTGCGAAACGCCACAGCTATTACAATTTTCGGCCAGGTGGGCATCCAGTGGATCGCCAGAAAGATCAATGAGTACCTGAACAAGGTGTGCGGCACCAACGACGAGGACTTCATCGCCGCTGGCGATACTGATAGCGTGTACGTTTGTGTGGACAAGGTCATCGAGAAGGTTGGCCTGGACAGATTTAAGGAACAGAACGACCTCGTGGAGTTCATGAACCAGTTCGGAAAGAAGAAGATGGAACCCATGATCGATGTGGCTTATAGAGAGCTGTGCGACTACATGAACAACAGAGAGCACCTGATGCACATGGATAGAGAAGCTATTTCTTGCCCTCCTCTGGGCTCTAAGGGAGTGGGCGGATTTTGGAAAGCCAAAAAGAGATACGCCCTGAATGTGTACGACATGGAAGATAAGAGATTCGCCGAGCCTCACCTGAAAATCATGGGCATGGAAACACAGCAGAGCAGCACCCCTAAGGCTGTGCAGGAGGCCCTGGAAGAGTCTATCCGGAGAATCTTGCAGGAGGGCGAGGAAAGCGTGCAGGAGTACTACAAGAACTTCGAGAAAGAATACAGACAGCTGGACTACAAGGTGATCGCGGAGGTGAAGACCGCTAATGATATCGCCAAGTACGACGACAAGGGCTGGCCCGGCTTCAAGTGCCCCTTCCACATCAGAGGCGTGCTCACCTACCGCAGAGCCGTTTCCGGCCTGGGCGTGGCCCCTATCCTGGATGGAAACAAAGTCATGGTGCTGCCTCTGAGAGAGGGCAACCCCTTTGGAGATAAATGCATCGCTTGGCCTAGCGGCACTGAGCTGCCCAAGGAAATCCGCTCCGACGTGCTGAGCTGGATCGATCACAGCACCCTGTTCCAAAAGTCCTTCGTGAAGCCCCTGGCCGGCATGTGCGAGTCCGCCGGCATGGACTACGAGGAAAAGGCCAGCCTGGATTTCCTGTTCGGCGGATCCggacctaagaaaaagaggaaggtggcggccgct TACCCATACGATGTTCCAGATTACGCT gctagcGGCAGTGGAGAGGGCAGAGGAAGTCTGCTAACATGCGGTGACGTCGAGGAGAATCCTGGCCCAGTGAGCAAGGGCGAGGAGCTGTTCACCGGGGTGGTGCCCATCCTGGTCGAGCTGGACGGCGACGTAAACGGCCACAAGTTCAGCGTGTCCGGCGAGGGCGAGGGCGATGCCACCTACGGCAAGCTGACCCTGAAGTTCATCTGCACCACCGGCAAGCTGCCCGTGCCCTGGCCCACCCTCGTGACCACCCTGACCTACGGCGTGCAGTGCTTCAGCCGCTACCCCGACCACATGAAGCAGCACGACTTCTTCAAGTCCGCCATGCCCGAAGGCTACGTCCAGGAGCGCACCATCTTCTTCAAGGACGACGGCAACTACAAGACCCGCGCCGAGGTGAAGTTCGAGGGCGACACCCTGGTGAACCGCATCGAGCTGAAGGGCATCGACTTCAAGGAGGACGGCAACATCCTGGGGCACAAGCTGGAGTACAACTACAACAGCCACAACGTCTATATCATGGCCGACAAGCAGAAGAACGGCATCAAGGTGAACTTCAAGATCCGCCACAACATCGAGGACGGCAGCGTGCAGCTCGCCGACCACTACCAGCAGAACACCCCCATCGGCGACGGCCCCGTGCTGCTGCCCGACAACCACTACCTGAGCACCCAGTCCGCCCTGAGCAAAGACCCCAACGAGAAGCGCGATCACATGGTCCTGCTGGAGTTCGTGACCGCCGCCGGGATCACTCTCGGCATGGACGAGCTGTACAAGTAA

MS2-Linker-NLS-**RB69** DNA polymerase-NLS-HA-T2A-GFP

atggcttcaaactttactcagttcgtgctcgtggacaatggtgggacaggggatgtgacagtggctccttctaatttcgctaatggggtggcagagtggatcagctccaactcacggagccaggcctacaaggtgacatgcagcgtcaggcagtctagtgcccagaagagaaagtataccatcaaggtggaggtccccaaagtggctacccagacagtgggcggagtcgaactgcctgtcgccgcttggaggtcctacctgaacatggagctcactatcccaattttcgctaccaattctgactgtgaactcatcgtgaaggcaatgcaggggctcctcaaagacggtaatcctatcccttccgccatcgccgctaactcaggtatctacagcgctggaggaggtggaagcggaggaggaggaagcggaggaggaggtagcggacctaagaaaaagaggaaggtgAAGGAGTTCTACCTGACCGTGGAGCAGATCGGCGACAGCATCTTCGAGAGATACATCGACAGCAACGGCAGAGAGAGAACAAGAGAGGTGGAGTACAAGCCAAGCCTGTTTGCGCACTGCCCCGAGAGCCAAGCCACCAAGTACTTCGACATCTACGGCAAGCCCTGCACAAGAAAGCTGTTCGCCAACATGAGAGACGCTTCTCAGTGGATCAAGAGAATGGAGGACATCGGCCTGGAGGCCCTGGGCATGGACGACTTCAAGCTGGCCTACCTGAGCGACACCTACAACTACGAGATCAAGTACGACCACACCAAGATCAGAGTGGCCAACTTCGACATCGAGGTGACAAGCCCCGACGGCTTCCCCGAGCCTAGCCAAGCCAAGCACCCCATCGACGCCATCACCCACTACGACAGCATCGACGACAGATTCTACGTGTTCGACCTGCTGAACAGCCCCTACGGCAACGTGGAGGAGTGGAGCATCGAGATCGCCGCGAAGCTGCAAGAGCAAGGCGGCGACGAGGTGCCTAGCGAAATCATCGATAAGATAATCTACATGCCCTTCGACAACGAGAAGGAGCTGCTGATGGAATATCTGAACTTTTGGCAGCAGAAGACCCCCGTGATCCTGACCGGCTGGAACGTGGAGAGCTTCGACATCCCCTACGTGTACAACAGAATCAAGAACATCTTCGGCGAGAGCACCGCCAAGAGACTGAGCCCCCACAGAAAGACAAGAGTGAAGGTGATCGAGAACATGTACGGCAGCAGAGAGATCATCACCCTGTTCGGCATCAGCGTGCTGGACTACATAGACCTGTACAAGAAGTTCAGCTTCACCAATCAGCCTAGCTACAGCCTGGACTACATCAGTGAATTCGAGCTGAACGTGGGCAAGCTGAAGTACGACGGCCCCATCAGCAAGCTGAGAGAGAGCAACCATCAGAGATACATCAGCTACAACATCATCGATGTGTACCGCGTACTGCAGATCGACGCCAAGAGACAGTTCATCAACCTGAGCCTGGACATGGGCTACTACGCCAAGATTCAGATTCAGAGCGTGTTCAGCCCCATCAAGACCTGGGACGCCATCATCTTCAACAGCCTGAAGGAGCAGAACAAGGTGATCCCCCAAGGCAGAAGCCACCCCGTGCAGCCCTACCCCGGCGCCTTCGTGAAGGAGCCCATCCCCAACAGATACAAGTACGTGATGAGCTTCGACCTGACAAGCCTGTACCCTAGCATCATCAGACAAGTGAACATCAGCCCCGAGACCATCGCCGGCACCTTCAAGGTGGCCCCCCTGCACGACTATATCAATGCCGTGGCCGAGAGACCTAGCGATGTCTACAGCTGCAGCCCCAACGGCATGATGTACTACAAGGACAGAGACGGCGTGGTGCCCACCGAAATCACGAAAGTGTTCAATCAGAGAAAGGAGCACAAGGGCTACATGCTGGCCGCTCAGAGAAACGGCGAGATCATCAAGGAGGCCCTGCACAACCCCAACCTGAGCGTGGACGAGCCCCTGGACGTGGACTACAGATTCGACTTCAGCGACGAGATCAAGGAGAAGATCAAGAAGCTGAGCGCCAAGAGCCTGAACGAGATGCTGTTCAGAGCTCAGAGAACCGAGGTGGCCGGCATGACCGCTCAGATCAACAGAAAGCTGCTGATCAACAGCCTGTACGGCGCCCTGGGCAACGTGTGGTTCAGATACTACGACCTGAGAAACGCCACCGCCATCACCACCTTCGGGCAGATGGCCCTGCAGTGGATCGAGAGAAAGGTGAACGAATACCTGAACGAGGTATGCGGCACCGAGGGCGAGGCCTTCGTGCTGTACGGCGACACCGACAGCATCTACGTGAGCGCCGACAAGATAATAGACAAGGTGGGTGAGAGCAAGTTCAGAGACACCAACCACTGGGTGGACTTCCTGGACAAGTTCGCTAGAGAGAGAATGGAGCCCGCCATCGACAGAGGCTTCAGAGAGATGTGCGAGTACATGAACAACAAGCAGCACCTGATGTTCATGGACAGAGAGGCCATCGCCGGCCCCCCCCTGGGCAGCAAGGGCATCGGCGGCTTCTGGACCGGCAAGAAGAGATACGCCCTGAACGTGTGGGACATGGAGGGCACAAGATACGCCGAGCCCAAGCTGAAGATCATGGGCCTGGAGACACAGAAGAGCAGCACCCCCAAGGCCGTGCAGAAGGCCCTGAAGGAGTGCATCAGAAGAATGCTGCAAGAGGGCGAGGAGAGCCTGCAAGAGTACTTCAAGGAGTTCGAGAAGGAGTTCAGACAGCTGAATTACATCAGCATCGCAAGCGTGAGCAGCGCCAACAACATCGCCAAGTACGACGTGGGCGGCTTCCCCGGCCCCAAGTGCCCCTTCCACATCAGAGGCATCCTGACCTACAACAGAGCCATCAAGGGCAACATCGACGCCCCCCAAGTGGTGGAGGGCGAGAAGGTGTACGTGCTGCCCCTGAGAGAGGGCAACCCCTTCGGCGACAAGTGCATCGCCTGGCCTAGCGGCACCGAGATCACCGACCTGATCAAGGACGACGTGCTGCACTGGATGGACTACACCGTGCTGCTGGAGAAGACCTTCATCAAGCCCCTGGAGGGCTTCACAAGCGCCGCCAAGCTGGATTACGAGAAGAAGGCTAGCCTCTTTGACATGTTCGACTTCGGATCCggacctaagaaaaagaggaaggtggcggccgctTACCCATACGATGTTCCAGATTACGCTgctagcGGCAGTGGAGAGGGCAGAGGAAGTCTGCTAACATGCGGTGACGTCGAGGAGAATCCTGGCCCAGTGAGCAAGGGCGAGGAGCTGTTCACCGGGGTGGTGCCCATCCTGGTCGAGCTGGACGGCGACGTAAACGGCCACAAGTTCAGCGTGTCCGGCGAGGGCGAGGGCGATGCCACCTACGGCAAGCTGACCCTGAAGTTCATCTGCACCACCGGCAAGCTGCCCGTGCCCTGGCCCACCCTCGTGACCACCCTGACCTACGGCGTGCAGTGCTTCAGCCGCTACCCCGACCACATGAAGCAGCACGACTTCTTCAAGTCCGCCATGCCCGAAGGCTACGTCCAGGAGCGCACCATCTTCTTCAAGGACGACGGCAACTACAAGACCCGCGCCGAGGTGAAGTTCGAGGGCGACACCCTGGTGAACCGCATCGAGCTGAAGGGCATCGACTTCAAGGAGGACGGCAACATCCTGGGGCACAAGCTGGAGTACAACTACAACAGCCACAACGTCTATATCATGGCCGACAAGCAGAAGAACGGCATCAAGGTGAACTTCAAGATCCGCCACAACATCGAGGACGGCAGCGTGCAGCTCGCCGACCACTACCAGCAGAACACCCCCATCGGCGACGGCCCCGTGCTGCTGCCCGACAACCACTACCTGAGCACCCAGTCCGCCCTGAGCAAAGACCCCAACGAGAAGCGCGATCACATGGTCCTGCTGGAGTTCGTGACCGCCGCCGGGATCACTCTCGGCATGGACGAGCTGTACAAGTAA

MS2-Linker-NLS-T7 DNA polymerase-NLS-HA-T2A-GFP

atggcttcaaactttactcagttcgtgctcgtggacaatggtgggacaggggatgtgacagtggctccttctaatttcgctaatggggtggcagagtggatcagctccaactcacggagccaggcctacaaggtgacatgcagcgtcaggcagtctagtgcccagaagagaaagtataccatcaaggtggaggtccccaaagtggctacccagacagtgggcggagtcgaactgcctgtcgccgcttggaggtcctacctgaacatggagctcactatcccaattttcgctaccaattctgactgtgaactcatcgtgaaggcaatgcaggggctcctcaaagacggtaatcctatcccttccgccatcgccgctaactcaggtatctacagcgctggaggaggtggaagcggaggaggaggaagcggaggaggaggtagcggacctaagaaaaagaggaaggtgATCGTGAGCGACATCGAGGCCAACGCCCTGCTGGAGAGCGTTACCAAGTTCCACTGCGGCGTGATCTACGACTACAGCACCGCCGAGTACGTGAGCTACAGACCTAGCGACTTCGGCGCCTACCTGGATGCTCTGGAGGCCGAAGTCGCTAGAGGCGGCCTGATCGTGTTCCACAACGGCCACAAGTACGACGTGCCCGCCCTGACCAAGCTGGCCAAGCTGCAGCTGAACAGAGAGTTCCACCTGCCTAGAGAGAACTGCATCGACACCCTGGTGCTGAGCAGACTGATCCACAGCAACCTGAAGGACACCGACATGGGCCTGCTGAGAAGCGGCAAGCTGCCCGGCAAGAGATTCGGCAGCCACGCCCTAGAGGCCTGGGGTTACAGACTGGGCGAGATGAAGGGCGAGTACAAGGACGACTTCAAGAGAATGCTGGAGGAGCAAGGCGAGGAGTACGTGGACGGCATGGAGTGGTGGAACTTCAACGAGGAGATGATGGACTACAACGTGCAAGACGTGGTGGTGACCAAGGCACTGCTGGAGAAATTGCTGAGCGACAAGCACTACTTCCCCCCCGAGATCGACTTCACCGACGTGGGCTACACCACCTTCTGGAGCGAGAGCCTGGAGGCCGTGGACATCGAGCACAGAGCCGCCTGGCTGCTGGCCAAGCAAGAGAGAAACGGCTTCCCCTTCGACACCAAGGCCATCGAGGAGCTGTACGTGGAGCTGGCCGCTAGAAGAAGCGAGCTGCTGAGAAAGCTGACCGAGACCTTCGGCAGCTGGTATCAGCCCAAGGGCGGCACCGAGATGTTCTGCCACCCTAGAACCGGAAAGCCGCTCCCCAAGTACCCTAGAATCAAGACCCCCAAGGTGGGCGGCATCTTCAAGAAGCCCAAGAACAAGGCTCAGAGAGAGGGCAGAGAGCCCTGCGAGCTGGACACAAGAGAGTACGTGGCCGGCGCCCCCTACACCCCCGTGGAGCACGTGGTGTTCAACCCTAGCAGCAGAGACCACATTCAGAAGAAGCTGCAAGAGGCCGGCTGGGTGCCCACCAAGTACACCGACAAGGGCGCCCCCGTGGTTGACGACGAAGTACTGGAGGGCGTGAGAGTGGACGACCCCGAGAAGCAAGCCGCCATCGACCTGATCAAGGAGTACCTGATGATTCAGAAGAGAATCGGGCAGAGCGCCGAGGGCGACAAGGCCTGGCTGAGATACGTGGCCGAGGACGGCAAAATTCACGGCTCGGTCAACCCCAACGGCGCCGTAACCGGCAGAGCAACCCACGCCTTCCCCAACCTGGCTCAGATCCCCGGCGTAAGAAGTCCTTATGGGGAACAGTGCCGCGCGGCGTTTGGCGCCGAGCACCATCTGGATGGCATAACCGGCAAGCCCTGGGTGCAAGCCGGCATCGATGCTAGCGGCCTGGAGCTGAGATGCCTGGCCCACTTCATGGCTAGATTCGACAACGGCGAGTACGCCCACGAGATCCTGAACGGCGACATCCACACCAAGAATCAGATCGCCGCCGAGCTGCCCACAAGAGACAACGCCAAGACCTTCATCTACGGCTTCCTGTACGGCGCCGGCGACGAGAAGATCGGGCAGATCGTGGGCGCCGGCAAGGAGAGAGGCAAGGAGCTGAAGAAGAAGTTCCTGGAGAACACCCCCGCCATCGCCGCCCTGAGAGAGAGCATTCAGCAGACCCTGGTGGAGAGCTCTCAGTGGGTGGCCGGCGAACAGCAAGTCAAGTGGAAGAGAAGATGGATCAAGGGCCTGGACGGCAGAAAGGTGCACGTGAGAAGTCCGCACGCCGCCCTGAACACCCTGCTGCAGAGCGCCGGCGCCCTGATCTGCAAGCTGTGGATCATCAAGACCGAGGAGATGCTGGTGGAGAAGGGCCTGAAGCACGGCTGGGACGGCGACTTCGCCTACATGGCCTGGGTGCACGACGAGATCCAAGTGGGCTGCAGAACCGAGGAGATCGCCCAAGTGGTGATCGAGACCGCCCAAGAGGCCATGAGATGGGTGGGCGACCACTGGAACTTCAGATGCCTGCTGGACACCGAGGGCAAGATGGGCCCCAACTGGGCCATCTGCCACGGATCCggacctaagaaaaagaggaaggtggcggccgctTACCCATACGATGTTCCAGATTACGCTgctagcGGCAGTGGAGAGGGCAGAGGAAGTCTGCTAACATGCGGTGACGTCGAGGAGAATCCTGGCCCAGTGAGCAAGGGCGAGGAGCTGTTCACCGGGGTGGTGCCCATCCTGGTCGAGCTGGACGGCGACGTAAACGGCCACAAGTTCAGCGTGTCCGGCGAGGGCGAGGGCGATGCCACCTACGGCAAGCTGACCCTGAAGTTCATCTGCACCACCGGCAAGCTGCCCGTGCCCTGGCCCACCCTCGTGACCACCCTGACCTACGGCGTGCAGTGCTTCAGCCGCTACCCCGACCACATGAAGCAGCACGACTTCTTCAAGTCCGCCATGCCCGAAGGCTACGTCCAGGAGCGCACCATCTTCTTCAAGGACGACGGCAACTACAAGACCCGCGCCGAGGTGAAGTTCGAGGGCGACACCCTGGTGAACCGCATCGAGCTGAAGGGCATCGACTTCAAGGAGGACGGCAACATCCTGGGGCACAAGCTGGAGTACAACTACAACAGCCACAACGTCTATATCATGGCCGACAAGCAGAAGAACGGCATCAAGGTGAACTTCAAGATCCGCCACAACATCGAGGACGGCAGCGTGCAGCTCGCCGACCACTACCAGCAGAACACCCCCATCGGCGACGGCCCCGTGCTGCTGCCCGACAACCACTACCTGAGCACCCAGTCCGCCCTGAGCAAAGACCCCAACGAGAAGCGCGATCACATGGTCCTGCTGGAGTTCGTGACCGCCGCCGGGATCACTCTCGGCATGGACGAGCTGTACAAGTAA

MS2-Linker-NLS-DNA polymerase 4-NLS-HA-T2A-GFP

AtggcttcaaactttactcagttcgtgctcgtggacaatggtgggacaggggatgtgacagtggctccttctaatttcgctaatggggtggcagagtggatcagctccaactcacggagccaggcctacaaggtgacatgcagcgtcaggcagtctagtgcccagaagagaaagtataccatcaaggtggaggtccccaaagtggctacccagacagtgggcggagtcgaactgcctgtcgccgcttggaggtcctacctgaacatggagctcactatcccaattttcgctaccaattctgactgtgaactcatcgtgaaggcaatgcaggggctcctcaaagacggtaatcctatcccttccgccatcgccgctaactcaggtatctacagcgctggaggaggtggaagcggaggaggaggaagcggaggaggaggtagcggacctaagaaaaagaggaaggtgtctctaaagggtaaatttttcgcctttttacctaatcctaacacatcttccaataagttctttaagagtatattggagaaaaagggcgccacaattgtgtcaagtattcaaaattgtcttcaatctagccgtaaggaagtggtcattttgattgaggactcctttgttgattctgatatgcatttgactcagaaagatattttccaaagggaagcaggcttaaatgatgtcgatgaatttcttggtaagattgaacagtcaggcattcaatgtgtgaaaaccagttgcatcacaaagtgggtccagaatgataaatttgcgtttcaaaaagatgatttgattaaatttcaaccatccattatcgttatatcagataacgctgatgacggacaaagttctactgataaagagagtgagatttcaactgacgtagaaagtgaaaggaatgatgacagcaacaataaagatatgatacaagcttcaaaacctcttaagcgactcttacagggggataaaggaagagcttcccttgttactgacaaaacgaagtacaaaaacaatgaattgattatcggagcgttgaaaaggttaacaaaaaaatatgagatcgaaggtgagaaatttcgtgcaagaagttatagactggctaaacagtcgatggaaaattgcgatttcaatgttcgttccggtgaagaagcacatactaaattaaggaatatcgggcctagtattgccaaaaaaatacaagttatattagatacgggagttttaccaggtttaaatgattcagtgggattagaagacaagttgaaatacttcaaaaattgttacggcattgggtcggaaattgctaaacgctggaatcttctaaattttgaaagcttttgtgttgcagctaagaaggacccagaggagtttgtatcagattggacaattttatttggttggtcatattacgacgattggttatgcaagatgtctcggaatgaatgtttcacacatttaaagaaggttcaaaaagcgctgcgtggcattgatcctgaatgccaagtcgaattacagggaagttataataggggctattccaagtgtggtgacattgatcttttatttttcaagccgttttgtaatgacacgaccgagttggcaaaaatcatggaaacgctttgtattaagttgtacaaggatggctatatccattgttttttacagctaacgccaaacttggaaaagctattcttaaaaagaatagtggagagatttcgtacagcgaagattgttgggtatggagaaagaaagaggtggtattcttctgagataatcaagaaatttttcatgggagtcaaattgtctccaagagaattagaagaactgaaagaaatgaaaaatgatgaaggcacattgttaattgaagaagaagaagaagaagaaacaaaattaaaaccgattgaccaatatatgtctctgaatgccaaggatggaaattattgcagaagattagactttttttgttgcaagtgggatgagcttggagcaggaagaatacactatactggatctaaagagtacaatagatggataagaatattggcagcgcaaaaaggcttcaagcttacacaacacggtttatttcgaaataatatccttctcgaaagctttaacgaacgcagaattttcgagttattaaacttaaaatacgctgaacccgaacatagaaatatcgaatgggaaaaaaaaactgcaGGATCCggacctaagaaaaagaggaaggtggcggccgctTACCCATACGATGTTCCAGATTACGCTgctagcGGCAGTGGAGAGGGCAGAGGAAGTCTGCTAACATGCGGTGACGTCGAGGAGAATCCTGGCCCAGTGAGCAAGGGCGAGGAGCTGTTCACCGGGGTGGTGCCCATCCTGGTCGAGCTGGACGGCGACGTAAACGGCCACAAGTTCAGCGTGTCCGGCGAGGGCGAGGGCGATGCCACCTACGGCAAGCTGACCCTGAAGTTCATCTGCACCACCGGCAAGCTGCCCGTGCCCTGGCCCACCCTCGTGACCACCCTGACCTACGGCGTGCAGTGCTTCAGCCGCTACCCCGACCACATGAAGCAGCACGACTTCTTCAAGTCCGCCATGCCCGAAGGCTACGTCCAGGAGCGCACCATCTTCTTCAAGGACGACGGCAACTACAAGACCCGCGCCGAGGTGAAGTTCGAGGGCGACACCCTGGTGAACCGCATCGAGCTGAAGGGCATCGACTTCAAGGAGGACGGCAACATCCTGGGGCACAAGCTGGAGTACAACTACAACAGCCACAACGTCTATATCATGGCCGACAAGCAGAAGAACGGCATCAAGGTGAACTTCAAGATCCGCCACAACATCGAGGACGGCAGCGTGCAGCTCGCCGACCACTACCAGCAGAACACCCCCATCGGCGACGGCCCCGTGCTGCTGCCCGACAACCACTACCTGAGCACCCAGTCCGCCCTGAGCAAAGACCCCAACGAGAAGCGCGATCACATGGTCCTGCTGGAGTTCGTGACCGCCGCCGGGATCACTCTCGGCATGGACGAGCTGTACAAGTAA

MS2-Linker-NLS-DNA polymerase I-NLS-HA-T2A-GFP

AtggcttcaaactttactcagttcgtgctcgtggacaatggtgggacaggggatgtgacagtggctccttctaatttcgctaatggggtggcagagtggatcagctccaactcacggagccaggcctacaaggtgacatgcagcgtcaggcagtctagtgcccagaagagaaagtataccatcaaggtggaggtccccaaagtggctacccagacagtgggcggagtcgaactgcctgtcgccgcttggaggtcctacctgaacatggagctcactatcccaattttcgctaccaattctgactgtgaactcatcgtgaaggcaatgcaggggctcctcaaagacggtaatcctatcccttccgccatcgccgctaactcaggtatctacagcgctggaggaggtggaagcggaggaggaggaagcggaggaggaggtagcggacctaagaaaaagaggaaggtgGTGCAGATCCCTCAGAACCCCCTGATCCTGGTGGACGGCAGCAGCTACCTGTACAGAGCCTATCACGCCTTCCCACCCTTAACTAACAGCGCCGGCGAGCCCACCGGCGCCATGTACGGCGTGCTGAACATGCTGAGAAGCCTGATCATGCAGTACAAGCCCACCCACGCCGCCGTGGTGTTCGACGCCAAGGGCAAGACCTTCAGAGACGAGCTGTTCGAGCACTACAAGAGCCACAGACCCCCCATGCCCGACGACCTGAGAGCTCAGATCGAGCCCCTGCACGCCATGGTTAAGGCCATGGGCTTACCTCTATTGGCGGTAAGCGGCGTGGAGGCCGATGACGTGATCGGCACCCTGGCTAGAGAGGCCGAGAAGGCCGGCAGACCCGTGCTGATCAGCACCGGCGACAAGGACATGGCTCAGCTGGTGACCCCCAACATCACCCTGATCAACACCATGACCAACACCATCCTGGGCCCCGAAGAGGTGGTGAACAAGTACGGCGTGCCCCCCGAGCTGATCATCGACTTCCTGGCCCTGATGGGCGACAGCAGCGATAACATACCCGGCGTACCCGGCGTGGGCGAAAAGACTGCCCAAGCCCTGCTCCAAGGCTTGGGCGGCCTGGACACCCTGTACGCCGAGCCCGAGAAGATCGCCGGACTGAGCTTCCGTGGGGCCAAGACCATGGCCGCCAAGTTAGAACAGAACAAAGAGGTGGCCTACCTGAGCTATCAGCTGGCCACCATCAAGACCGACGTGGAGCTGGAGCTGACCTGCGAGCAGCTGGAGGTGCAGCAGCCCGCGGCCGAAGAACTTCTGGGCCTGTTCAAGAAGTACGAGTTCAAGCGTTGGACCGCTGACGTGGAGGCCGGCAAATGGCTCCAAGCGAAAGGGGCAAAACCCGCTGCCAAGCCCCAAGAGACAAGCGTAGCCGACGAGGCCCCCGAGGTAACCGCGACAGTGATCAGCTACGACAACTACGTGACCATCCTGGACGAGGAGACCCTGAAGGCCTGGATCGCCAAACTGGAGAAAGCCCCCGTGTTCGCCTTCGACACGGAAACAGACAGCCTGGACAACATCAGCGCCAACTTAGTGGGGCTGAGCTTTGCCATCGAACCCGGCGTGGCTGCCTACATTCCTGTGGCCCACGACTACCTGGACGCCCCCGATCAGATCAGCAGAGAGAGAGCCCTGGAGCTGCTGAAGCCCCTGCTGGAGGACGAGAAGGCCCTGAAGGTGGGGCAGAACCTGAAGTACGACAGAGGCATCCTGGCCAACTACGGCATAGAGTTGAGAGGCATTGCCTTCGACACCATGCTCGAGAGCTACATCCTGAACAGCGTGGCCGGCAGACACGACATGGACAGCCTGGCCGAGAGATGGCTGAAGCACAAGACCATCACCTTCGAGGAGATCGCCGGCAAGGGCAAGAATCAGCTGACCTTCAATCAGATCGCCCTGGAGGAGGCCGGCAGATACGCGGCTGAGGACGCCGACGTGACCCTGCAGCTGCACCTGAAGATGTGGCCCGACCTGCAGAAGCACAAGGGCCCCCTGAACGTGTTCGAGAACATCGAGATGCCCCTGGTGCCCGTGCTGAGCAGAATCGAGAGAAACGGCGTGAAGATCGACCCCAAGGTGCTGCACAACCACAGCGAGGAGCTGACCCTGAGACTGGCCGAGCTGGAGAAGAAGGCCCACGAAATCGCCGGCGAAGAGTTCAACCTGTCGAGCACCAAACAGCTGCAGACCATCCTGTTCGAGAAACAAGGCATCAAGCCGCTAAAAAAAACCCCCGGCGGCGCCCCTAGCACTAGCGAAGAGGTGCTGGAGGAGCTGGCCCTGGACTACCCCCTGCCCAAGGTGATCCTGGAGTACAGAGGCCTGGCCAAGCTGAAGAGCACCTACACCGACAAGCTGCCCCTGATGATCAACCCCAAGACCGGCAGAGTGCACACAAGCTACCACCAAGCCGTGACCGCTACGGGCAGACTGAGCAGCACCGATCCAAACCTGCAGAACATCCCCGTGAGAAACGAGGAGGGCAGAAGAATCAGACAAGCCTTCATCGCCCCCGAGGACTACGTGATCGTGAGCGCCGACTACTCTCAGATCGAGCTGAGAATCATGGCCCACCTGAGCAGAGACAAGGGCCTGCTGACCGCCTTCGCCGAGGGCAAGGACATCCACAGAGCCACCGCCGCCGAGGTCTTTGGCCTGCCCCTGGAGACCGTGACAAGCGAGCAGAGAAGAAGCGCCAAGGCCATCAACTTCGGCCTGATCTACGGCATGAGCGCCTTCGGCCTGGCTAGACAGCTGAACATCCCTAGAAAGGAGGCTCAGAAGTACATGGACCTGTACTTCGAGAGATACCCCGGCGTGCTGGAGTACATGGAGAGAACAAGAGCCCAAGCCAAGGAGCAAGGCTACGTGGAGACCCTGGACGGCAGAAGACTGTACCTGCCCGACATCAAGAGCAGCAATGGAGCTAGACGAGCCGCCGCCGAGAGAGCCGCCATCAACGCCCCCATGCAAGGCACCGCCGCAGACATCATCAAGAGAGCCATGATCGCCGTGGACGCCTGGCTGCAAGCCGAGCAGCCTAGAGTGAGAATGATCATGCAAGTGCACGACGAGCTGGTGTTCGAGGTGCACAAGGACGACGTGGACGCCGTGGCCAAGCAGATCCATCAGCTGATGGAGAACTGCACAAGACTGGACGTCCCCCTCCTTGTGGAAGTGGGCAGCGGCGAGAACTGGGACCAAGCCCACGGATCCggacctaagaaaaagaggaaggtggcggccgctTACCCATACGATGTTCCAGATTACGCTgctagcGGCAGTGGAGAGGGCAGAGGAAGTCTGCTAACATGCGGTGACGTCGAGGAGAATCCTGGCCCAGTGAGCAAGGGCGAGGAGCTGTTCACCGGGGTGGTGCCCATCCTGGTCGAGCTGGACGGCGACGTAAACGGCCACAAGTTCAGCGTGTCCGGCGAGGGCGAGGGCGATGCCACCTACGGCAAGCTGACCCTGAAGTTCATCTGCACCACCGGCAAGCTGCCCGTGCCCTGGCCCACCCTCGTGACCACCCTGACCTACGGCGTGCAGTGCTTCAGCCGCTACCCCGACCACATGAAGCAGCACGACTTCTTCAAGTCCGCCATGCCCGAAGGCTACGTCCAGGAGCGCACCATCTTCTTCAAGGACGACGGCAACTACAAGACCCGCGCCGAGGTGAAGTTCGAGGGCGACACCCTGGTGAACCGCATCGAGCTGAAGGGCATCGACTTCAAGGAGGACGGCAACATCCTGGGGCACAAGCTGGAGTACAACTACAACAGCCACAACGTCTATATCATGGCCGACAAGCAGAAGAACGGCATCAAGGTGAACTTCAAGATCCGCCACAACATCGAGGACGGCAGCGTGCAGCTCGCCGACCACTACCAGCAGAACACCCCCATCGGCGACGGCCCCGTGCTGCTGCCCGACAACCACTACCTGAGCACCCAGTCCGCCCTGAGCAAAGACCCCAACGAGAAGCGCGATCACATGGTCCTGCTGGAGTTCGTGACCGCCGCCGGGATCACTCTCGGCATGGACGAGCTGTACAAGTAA

MS2-Linker-NLS-Klenow fragment-NLS-HA-T2A-GFP

AtggcttcaaactttactcagttcgtgctcgtggacaatggtgggacaggggatgtgacagtggctccttctaatttcgctaatggggtggcagagtggatcagctccaactcacggagccaggcctacaaggtgacatgcagcgtcaggcagtctagtgcccagaagagaaagtataccatcaaggtggaggtccccaaagtggctacccagacagtgggcggagtcgaactgcctgtcgccgcttggaggtcctacctgaacatggagctcactatcccaattttcgctaccaattctgactgtgaactcatcgtgaaggcaatgcaggggctcctcaaagacggtaatcctatcccttccgccatcgccgctaactcaggtatctacagcgctggaggaggtggaagcggaggaggaggaagcggaggaggaggtagcggacctaagaaaaagaggaaggtgGTGATCAGCTACGACAACTACGTGACCATCCTGGACGAGGAGACCCTGAAGGCCTGGATCGCCAAACTGGAGAAAGCCCCCGTGTTCGCCTTCGACACGGAAACAGACAGCCTGGACAACATCAGCGCCAACTTAGTGGGGCTGAGCTTTGCCATCGAACCCGGCGTGGCTGCCTACATTCCTGTGGCCCACGACTACCTGGACGCCCCCGATCAGATCAGCAGAGAGAGAGCCCTGGAGCTGCTGAAGCCCCTGCTGGAGGACGAGAAGGCCCTGAAGGTGGGGCAGAACCTGAAGTACGACAGAGGCATCCTGGCCAACTACGGCATAGAGTTGAGAGGCATTGCCTTCGACACCATGCTCGAGAGCTACATCCTGAACAGCGTGGCCGGCAGACACGACATGGACAGCCTGGCCGAGAGATGGCTGAAGCACAAGACCATCACCTTCGAGGAGATCGCCGGCAAGGGCAAGAATCAGCTGACCTTCAATCAGATCGCCCTGGAGGAGGCCGGCAGATACGCGGCTGAGGACGCCGACGTGACCCTGCAGCTGCACCTGAAGATGTGGCCCGACCTGCAGAAGCACAAGGGCCCCCTGAACGTGTTCGAGAACATCGAGATGCCCCTGGTGCCCGTGCTGAGCAGAATCGAGAGAAACGGCGTGAAGATCGACCCCAAGGTGCTGCACAACCACAGCGAGGAGCTGACCCTGAGACTGGCCGAGCTGGAGAAGAAGGCCCACGAAATCGCCGGCGAAGAGTTCAACCTGTCGAGCACCAAACAGCTGCAGACCATCCTGTTCGAGAAACAAGGCATCAAGCCGCTAAAAAAAACCCCCGGCGGCGCCCCTAGCACTAGCGAAGAGGTGCTGGAGGAGCTGGCCCTGGACTACCCCCTGCCCAAGGTGATCCTGGAGTACAGAGGCCTGGCCAAGCTGAAGAGCACCTACACCGACAAGCTGCCCCTGATGATCAACCCCAAGACCGGCAGAGTGCACACAAGCTACCACCAAGCCGTGACCGCTACGGGCAGACTGAGCAGCACCGATCCAAACCTGCAGAACATCCCCGTGAGAAACGAGGAGGGCAGAAGAATCAGACAAGCCTTCATCGCCCCCGAGGACTACGTGATCGTGAGCGCCGACTACTCTCAGATCGAGCTGAGAATCATGGCCCACCTGAGCAGAGACAAGGGCCTGCTGACCGCCTTCGCCGAGGGCAAGGACATCCACAGAGCCACCGCCGCCGAGGTCTTTGGCCTGCCCCTGGAGACCGTGACAAGCGAGCAGAGAAGAAGCGCCAAGGCCATCAACTTCGGCCTGATCTACGGCATGAGCGCCTTCGGCCTGGCTAGACAGCTGAACATCCCTAGAAAGGAGGCTCAGAAGTACATGGACCTGTACTTCGAGAGATACCCCGGCGTGCTGGAGTACATGGAGAGAACAAGAGCCCAAGCCAAGGAGCAAGGCTACGTGGAGACCCTGGACGGCAGAAGACTGTACCTGCCCGACATCAAGAGCAGCAATGGAGCTAGACGAGCCGCCGCCGAGAGAGCCGCCATCAACGCCCCCATGCAAGGCACCGCCGCAGACATCATCAAGAGAGCCATGATCGCCGTGGACGCCTGGCTGCAAGCCGAGCAGCCTAGAGTGAGAATGATCATGCAAGTGCACGACGAGCTGGTGTTCGAGGTGCACAAGGACGACGTGGACGCCGTGGCCAAGCAGATCCATCAGCTGATGGAGAACTGCACAAGACTGGACGTCCCCCTCCTTGTGGAAGTGGGCAGCGGCGAGAACTGGGACCAAGCCCACGGATCCggacctaagaaaaagaggaaggtggcggccgctTACCCATACGATGTTCCAGATTACGCTgctagcGGCAGTGGAGAGGGCAGAGGAAGTCTGCTAACATGCGGTGACGTCGAGGAGAATCCTGGCCCAGTGAGCAAGGGCGAGGAGCTGTTCACCGGGGTGGTGCCCATCCTGGTCGAGCTGGACGGCGACGTAAACGGCCACAAGTTCAGCGTGTCCGGCGAGGGCGAGGGCGATGCCACCTACGGCAAGCTGACCCTGAAGTTCATCTGCACCACCGGCAAGCTGCCCGTGCCCTGGCCCACCCTCGTGACCACCCTGACCTACGGCGTGCAGTGCTTCAGCCGCTACCCCGACCACATGAAGCAGCACGACTTCTTCAAGTCCGCCATGCCCGAAGGCTACGTCCAGGAGCGCACCATCTTCTTCAAGGACGACGGCAACTACAAGACCCGCGCCGAGGTGAAGTTCGAGGGCGACACCCTGGTGAACCGCATCGAGCTGAAGGGCATCGACTTCAAGGAGGACGGCAACATCCTGGGGCACAAGCTGGAGTACAACTACAACAGCCACAACGTCTATATCATGGCCGACAAGCAGAAGAACGGCATCAAGGTGAACTTCAAGATCCGCCACAACATCGAGGACGGCAGCGTGCAGCTCGCCGACCACTACCAGCAGAACACCCCCATCGGCGACGGCCCCGTGCTGCTGCCCGACAACCACTACCTGAGCACCCAGTCCGCCCTGAGCAAAGACCCCAACGAGAAGCGCGATCACATGGTCCTGCTGGAGTTCGTGACCGCCGCCGGGATCACTCTCGGCATGGACGAGCTGTACAAGTAA

MS2-Linker-NLS-DNA polymerase lambda-NLS-HA-T2A-GFP

AtggcttcaaactttactcagttcgtgctcgtggacaatggtgggacaggggatgtgacagtggctccttctaatttcgctaatggggtggcagagtggatcagctccaactcacggagccaggcctacaaggtgacatgcagcgtcaggcagtctagtgcccagaagagaaagtataccatcaaggtggaggtccccaaagtggctacccagacagtgggcggagtcgaactgcctgtcgccgcttggaggtcctacctgaacatggagctcactatcccaattttcgctaccaattctgactgtgaactcatcgtgaaggcaatgcaggggctcctcaaagacggtaatcctatcccttccgccatcgccgctaactcaggtatctacagcgctggaggaggtggaagcggaggaggaggaagcggaggaggaggtagcggacctaagaaaaagaggaaggtggatcccaggggtatcttgaaggcatttcccaagcggcagaaaattcatgctgatgcatcatcaaaagtacttgcaaagattcctaggagggaagagggagaagaagcagaagagtggctgagctcccttcgggcccatgttgtgcgcactggcattggacgagcccgggcagaactctttgagaagcagattgttcagcatggcggccagctatgccctgcccagggcccaggtgtcactcacattgtggtggatgaaggcatggactatgagcgagccctccgccttctcagactaccccagctgcccccgggtgctcagctggtgaagtcagcctggctgagcttgtgccttcaggagaggaggctggtggatgtagctggattcagcatcttcatccccagtaggtacttggaccatccacagcccagcaaggcagagcaggatgcttctattcctcctggcacccatgaggccctgcttcagacagccctttctcctcctcctcctcccaccaggcctgtgtctcctccccaaaaggcaaaagaggcaccaaacacccaagcccagcccatctctgatgatgaagccagtgatggggaagaaacccaggttagtgcagctgatctggaagccctcatcagtggccactaccccacctcccttgagggagattgtgagcctagcccagcccctgctgtcctggataagtgggtctgtgcacagccctcaagccagaaggcgaccaatcacaacctccatatcacagagaagctggaagttctggccaaagcctacagtgttcagggagacaagtggagggccctgggctatgccaaggccatcaatgccctcaagagcttccataagcctgtcacctcgtaccaggaggcctgcagtatccctgggattgggaagcggatggctgagaaaatcatagagatcctggagagcgggcatttgcggaagctggaccatatcagtgagagcgtgcctgtcttggagctcttctccaacatctggggagctgggaccaagactgcccagatgtggtaccaacagggcttccgaagtctggaagacatccgcagccaggcctccctgacaacccagcaggccatcggcctgaagcattacagtgacttcctggaacgtatgcccagggaggaggctacagagattgagcagacagtccagaaagcagcccaggcctttaactctgggctgctgtgtgtggcatgtggttcataccgacggggaaaggcgacctgtggtgatgtcgacgtgctcatcactcacccagatggccggtcccaccggggtatcttcagccgcctccttgacagtcttcggcaggaagggttcctcacagatgacttggtgagccaagaggagaatggtcagcaacagaagtacttgggggtgtgccggctcccagggccagggcggcggcaccggcgcctggacatcatcgtggtgccctatagcgagtttgcctgtgccctgctctacttcaccggctctgcacacttcaaccgctccatgcgagccctggccaaaaccaagggcatgagtctgtcagaacatgccctcagcactgctgtggtccggaacacccatggctgcaaggtggggcctggccgagtgctgcccactcccactgagaaggatgtcttcaggctcttaggcctcccctaccgagaacctgctgagcgggactggGGATCCggacctaagaaaaagaggaaggtggcggccgctTACCCATACGATGTTCCAGATTACGCTgctagcGGCAGTGGAGAGGGCAGAGGAAGTCTGCTAACATGCGGTGACGTCGAGGAGAATCCTGGCCCAGTGAGCAAGGGCGAGGAGCTGTTCACCGGGGTGGTGCCCATCCTGGTCGAGCTGGACGGCGACGTAAACGGCCACAAGTTCAGCGTGTCCGGCGAGGGCGAGGGCGATGCCACCTACGGCAAGCTGACCCTGAAGTTCATCTGCACCACCGGCAAGCTGCCCGTGCCCTGGCCCACCCTCGTGACCACCCTGACCTACGGCGTGCAGTGCTTCAGCCGCTACCCCGACCACATGAAGCAGCACGACTTCTTCAAGTCCGCCATGCCCGAAGGCTACGTCCAGGAGCGCACCATCTTCTTCAAGGACGACGGCAACTACAAGACCCGCGCCGAGGTGAAGTTCGAGGGCGACACCCTGGTGAACCGCATCGAGCTGAAGGGCATCGACTTCAAGGAGGACGGCAACATCCTGGGGCACAAGCTGGAGTACAACTACAACAGCCACAACGTCTATATCATGGCCGACAAGCAGAAGAACGGCATCAAGGTGAACTTCAAGATCCGCCACAACATCGAGGACGGCAGCGTGCAGCTCGCCGACCACTACCAGCAGAACACCCCCATCGGCGACGGCCCCGTGCTGCTGCCCGACAACCACTACCTGAGCACCCAGTCCGCCCTGAGCAAAGACCCCAACGAGAAGCGCGATCACATGGTCCTGCTGGAGTTCGTGACCGCCGCCGGGATCACTCTCGGCATGGACGAGCTGTACAAGTAA

MS2-Linker-NLS-DNA polymerase mu-NLS-HA-T2A-GFP

AtggcttcaaactttactcagttcgtgctcgtggacaatggtgggacaggggatgtgacagtggctccttctaatttcgctaatggggtggcagagtggatcagctccaactcacggagccaggcctacaaggtgacatgcagcgtcaggcagtctagtgcccagaagagaaagtataccatcaaggtggaggtccccaaagtggctacccagacagtgggcggagtcgaactgcctgtcgccgcttggaggtcctacctgaacatggagctcactatcccaattttcgctaccaattctgactgtgaactcatcgtgaaggcaatgcaggggctcctcaaagacggtaatcctatcccttccgccatcgccgctaactcaggtatctacagcgctggaggaggtggaagcggaggaggaggaagcggaggaggaggtagcggacctaagaaaaagaggaaggtgctccccaaacggcggcgagcgcgggtcgggtcccctagcggcgatgccgcttcctccacgccgccctcgacgcgcttcccgggagtcgccatctacctggtcgagcctcgcatgggtcgcagccgccgggccttcctcacaggcctggcgcgctccaaaggcttccgcgtccttgacgcctgcagctccgaagcgacacatgttgtgatggaagagacctcagcagaggaggccgtcagctggcaggagcgcaggatggcagctgctcccccgggttgcacccccccagctctgctggacataagctggttaacagagagcctgggagctgggcagcctgtacctgtggagtgccggcaccgcctggaggtggctgggccaaggaaggggcctctgagcccagcatggatgcctgcctatgcctgccagcgccctacgcccctcacacaccacaacactggcctctccgaggctctggagatactggccgaggcagcaggctttgaaggcagtgagggccgcctcctcaccttctgcagagcagcctcggtgctcaaggcccttcccagccctgtcacaaccctgagccagctgcaggggcttccccactttggagaacactcctctagggttgtccaggagctgctggagcatggagtgtgtgaggaggtggagagagttcggcgctcagagaggtaccagaccatgaagctcttcacccagatcttcggggtcggtgtgaagactgctgaccggtggtaccgggaaggactgcgaaccttagatgacctccgagagcagccccagaaactaacccaacagcagaaagcggggctccagcaccaccaggacctgagcaccccagtcctgcggtccgatgtagatgccctgcagcaggtggtggaggaagctgtggggcaggccctgcctggggccaccgtcacgctgaccggcggcttccgcagggggaagttgcagggccatgacgtggacttcctcatcacccaccccaaggagggtcaggaggcggggctgctgcctagagtgatgtgccgcctgcaggaccagggcctcatcctgtaccaccagcaccagcacagctgctgtgagtcccctacccgcctggcccaacagagccacatggacgcttttgagagaagtttctgcattttccgcctaccacaacctccaggggctgctgtggggggatccacgaggccctgcccatcctggaaggccgtgagagtggacttggtagttgcacccgtcagccagttccctttcgccctgctcggttggactggctccaagcttttccagcgggagctgcgccgcttcagccggaaggagaagggcctgtggctgaacagccatgggctgtttgacccggagcagaagacatttttccaagcggcttcagaggaagacatcttcagacacctgggccttgagtaccttcctccagagcagagaaacgccGGATCCggacctaagaaaaagaggaaggtggcggccgctTACCCATACGATGTTCCAGATTACGCTgctagcGGCAGTGGAGAGGGCAGAGGAAGTCTGCTAACATGCGGTGACGTCGAGGAGAATCCTGGCCCAGTGAGCAAGGGCGAGGAGCTGTTCACCGGGGTGGTGCCCATCCTGGTCGAGCTGGACGGCGACGTAAACGGCCACAAGTTCAGCGTGTCCGGCGAGGGCGAGGGCGATGCCACCTACGGCAAGCTGACCCTGAAGTTCATCTGCACCACCGGCAAGCTGCCCGTGCCCTGGCCCACCCTCGTGACCACCCTGACCTACGGCGTGCAGTGCTTCAGCCGCTACCCCGACCACATGAAGCAGCACGACTTCTTCAAGTCCGCCATGCCCGAAGGCTACGTCCAGGAGCGCACCATCTTCTTCAAGGACGACGGCAACTACAAGACCCGCGCCGAGGTGAAGTTCGAGGGCGACACCCTGGTGAACCGCATCGAGCTGAAGGGCATCGACTTCAAGGAGGACGGCAACATCCTGGGGCACAAGCTGGAGTACAACTACAACAGCCACAACGTCTATATCATGGCCGACAAGCAGAAGAACGGCATCAAGGTGAACTTCAAGATCCGCCACAACATCGAGGACGGCAGCGTGCAGCTCGCCGACCACTACCAGCAGAACACCCCCATCGGCGACGGCCCCGTGCTGCTGCCCGACAACCACTACCTGAGCACCCAGTCCGCCCTGAGCAAAGACCCCAACGAGAAGCGCGATCACATGGTCCTGCTGGAGTTCGTGACCGCCGCCGGGATCACTCTCGGCATGGACGAGCTGTACAAGTAA

MS2-Linker-NLS-DNA polymerase beta-NLS-HA-T2A-GFP

AtggcttcaaactttactcagttcgtgctcgtggacaatggtgggacaggggatgtgacagtggctccttctaatttcgctaatggggtggcagagtggatcagctccaactcacggagccaggcctacaaggtgacatgcagcgtcaggcagtctagtgcccagaagagaaagtataccatcaaggtggaggtccccaaagtggctacccagacagtgggcggagtcgaactgcctgtcgccgcttggaggtcctacctgaacatggagctcactatcccaattttcgctaccaattctgactgtgaactcatcgtgaaggcaatgcaggggctcctcaaagacggtaatcctatcccttccgccatcgccgctaactcaggtatctacagcgctggaggaggtggaagcggaggaggaggaagcggaggaggaggtagcggacctaagaaaaagaggaaggtgagcaaacggaaggcgccgcaggagactctcaacgggggaatcaccgacatgctcacagaactcgcaaactttgagaagaacgtgagccaagctatccacaagtacaatgcttacagaaaagcagcatctgttatagcaaaatacccacacaaaataaagagtggagctgaagctaagaaattgcctggagtaggaacaaaaattgctgaaaagattgatgagtttttagcaactggaaaattacgtaaactggaaaagattcggcaggatgatacgagttcatccatcaatttcctgactcgagttagtggcattggtccatctgctgcaaggaagtttgtagatgaaggaattaaaacactagaagatctcagaaaaaatgaagataaattgaaccatcatcagcgaattgggctgaaatattttggggactttgaaaaaagaattcctcgtgaagagatgttacaaatgcaagatattgtactaaatgaagttaaaaaagtggattctgaatacattgctacagtctgtggcagtttcagaagaggtgcagagtccagtggtgacatggatgttctcctgacccatcccagcttcacttcagaatcaaccaaacagccaaaactgttacatcaggttgtggagcagttacaaaaggttcattttatcacagataccctgtcaaagggtgagacaaagttcatgggtgtttgccagcttcccagtaaaaatgatgaaaaagaatatccacacagaagaattgatatcaggttgatacccaaagatcagtattactgtggtgttctctatttcactgggagtgatattttcaataagaatatgagggctcatgccctagaaaagggtttcacaatcaatgagtacaccatccgtcccttgggagtcactggagttgcaggagaacccctgccagtggatagtgaaaaagacatctttgattacatccagtggaaataccgggaacccaaggaccggagcgaaGGATCCggacctaagaaaaagaggaaggtggcggccgctTACCCATACGATGTTCCAGATTACGCTgctagcGGCAGTGGAGAGGGCAGAGGAAGTCTGCTAACATGCGGTGACGTCGAGGAGAATCCTGGCCCAGTGAGCAAGGGCGAGGAGCTGTTCACCGGGGTGGTGCCCATCCTGGTCGAGCTGGACGGCGACGTAAACGGCCACAAGTTCAGCGTGTCCGGCGAGGGCGAGGGCGATGCCACCTACGGCAAGCTGACCCTGAAGTTCATCTGCACCACCGGCAAGCTGCCCGTGCCCTGGCCCACCCTCGTGACCACCCTGACCTACGGCGTGCAGTGCTTCAGCCGCTACCCCGACCACATGAAGCAGCACGACTTCTTCAAGTCCGCCATGCCCGAAGGCTACGTCCAGGAGCGCACCATCTTCTTCAAGGACGACGGCAACTACAAGACCCGCGCCGAGGTGAAGTTCGAGGGCGACACCCTGGTGAACCGCATCGAGCTGAAGGGCATCGACTTCAAGGAGGACGGCAACATCCTGGGGCACAAGCTGGAGTACAACTACAACAGCCACAACGTCTATATCATGGCCGACAAGCAGAAGAACGGCATCAAGGTGAACTTCAAGATCCGCCACAACATCGAGGACGGCAGCGTGCAGCTCGCCGACCACTACCAGCAGAACACCCCCATCGGCGACGGCCCCGTGCTGCTGCCCGACAACCACTACCTGAGCACCCAGTCCGCCCTGAGCAAAGACCCCAACGAGAAGCGCGATCACATGGTCCTGCTGGAGTTCGTGACCGCCGCCGGGATCACTCTCGGCATGGACGAGCTGTACAAGTAA

Cas9-L-T4 DNA polymerase

atggactataaggaccacgacggagactacaaggatcatgatattgattacaaagacgatgacgataagatggccccaaagaagaagcggaaggtcggtatccacggagtcccagcagccgacaagaagtacagcatcggcctggacatcggcaccaactctgtgggctgggccgtgatcaccgacgagtacaaggtgcccagcaagaaattcaaggtgctgggcaacaccgaccggcacagcatcaagaagaacctgatcggagccctgctgttcgacagcggcgaaacagccgaggccacccggctgaagagaaccgccagaagaagatacaccagacggaagaaccggatctgctatctgcaagagatcttcagcaacgagatggccaaggtggacgacagcttcttccacagactggaagagtccttcctggtggaagaggataagaagcacgagcggcaccccatcttcggcaacatcgtggacgaggtggcctaccacgagaagtaccccaccatctaccacctgagaaagaaactggtggacagcaccgacaaggccgacctgcggctgatctatctggccctggcccacatgatcaagttccggggccacttcctgatcgagggcgacctgaaccccgacaacagcgacgtggacaagctgttcatccagctggtgcagacctacaaccagctgttcgaggaaaaccccatcaacgccagcggcgtggacgccaaggccatcctgtctgccagactgagcaagagcagacggctggaaaatctgatcgcccagctgcccggcgagaagaagaatggcctgttcggaaacctgattgccctgagcctgggcctgacccccaacttcaagagcaacttcgacctggccgaggatgccaaactgcagctgagcaaggacacctacgacgacgacctggacaacctgctggcccagatcggcgaccagtacgccgacctgtttctggccgccaagaacctgtccgacgccatcctgctgagcgacatcctgagagtgaacaccgagatcaccaaggcccccctgagcgcctctatgatcaagagatacgacgagcaccaccaggacctgaccctgctgaaagctctcgtgcggcagcagctgcctgagaagtacaaagagattttcttcgaccagagcaagaacggctacgccggctacattgacggcggagccagccaggaagagttctacaagttcatcaagcccatcctggaaaagatggacggcaccgaggaactgctcgtgaagctgaacagagaggacctgctgcggaagcagcggaccttcgacaacggcagcatcccccaccagatccacctgggagagctgcacgccattctgcggcggcaggaagatttttacccattcctgaaggacaaccgggaaaagatcgagaagatcctgaccttccgcatcccctactacgtgggccctctggccaggggaaacagcagattcgcctggatgaccagaaagagcgaggaaaccatcaccccctggaacttcgaggaagtggtggacaagggcgcttccgcccagagcttcatcgagcggatgaccaacttcgataagaacctgcccaacgagaaggtgctgcccaagcacagcctgctgtacgagtacttcaccgtgtataacgagctgaccaaagtgaaatacgtgaccgagggaatgagaaagcccgccttcctgagcggcgagcagaaaaaggccatcgtggacctgctgttcaagaccaaccggaaagtgaccgtgaagcagctgaaagaggactacttcaagaaaatcgagtgcttcgactccgtggaaatctccggcgtggaagatcggttcaacgcctccctgggcacataccacgatctgctgaaaattatcaaggacaaggacttcctggacaatgaggaaaacgaggacattctggaagatatcgtgctgaccctgacactgtttgaggacagagagatgatcgaggaacggctgaaaacctatgcccacctgttcgacgacaaagtgatgaagcagctgaagcggcggagatacaccggctggggcaggctgagccggaagctgatcaacggcatccgggacaagcagtccggcaagacaatcctggatttcctgaagtccgacggcttcgccaacagaaacttcatgcagctgatccacgacgacagcctgacctttaaagaggacatccagaaagcccaggtgtccggccagggcgatagcctgcacgagcacattgccaatctggccggcagccccgccattaagaagggcatcctgcagacagtgaaggtggtggacgagctcgtgaaagtgatgggccggcacaagcccgagaacatcgtgatcgaaatggccagagagaaccagaccacccagaagggacagaagaacagccgcgagagaatgaagcggatcgaagagggcatcaaagagctgggcagccagatcctgaaagaacaccccgtggaaaacacccagctgcagaacgagaagctgtacctgtactacctgcagaatgggcgggatatgtacgtggaccaggaactggacatcaaccggctgtccgactacgatgtggaccatatcgtgcctcagagctttctgaaggacgactccatcgacaacaaggtgctgaccagaagcgacaagaaccggggcaagagcgacaacgtgccctccgaagaggtcgtgaagaagatgaagaactactggcggcagctgctgaacgccaagctgattacccagagaaagttcgacaatctgaccaaggccgagagaggcggcctgagcgaactggataaggccggcttcatcaagagacagctggtggaaacccggcagatcacaaagcacgtggcacagatcctggactcccggatgaacactaagtacgacgagaatgacaagctgatccgggaagtgaaagtgatcaccctgaagtccaagctggtgtccgatttccggaaggatttccagttttacaaagtgcgcgagatcaacaactaccaccacgcccacgacgcctacctgaacgccgtcgtgggaaccgccctgatcaaaaagtaccctaagctggaaagcgagttcgtgtacggcgactacaaggtgtacgacgtgcggaagatgatcgccaagagcgagcaggaaatcggcaaggctaccgccaagtacttcttctacagcaacatcatgaactttttcaagaccgagattaccctggccaacggcgagatccggaagcggcctctgatcgagacaaacggcgaaaccggggagatcgtgtgggataagggccgggattttgccaccgtgcggaaagtgctgagcatgccccaagtgaatatcgtgaaaaagaccgaggtgcagacaggcggcttcagcaaagagtctatcctgcccaagaggaacagcgataagctgatcgccagaaagaaggactgggaccctaagaagtacggcggcttcgacagccccaccgtggcctattctgtgctggtggtggccaaagtggaaaagggcaagtccaagaaactgaagagtgtgaaagagctgctggggatcaccatcatggaaagaagcagcttcgagaagaatcccatcgactttctggaagccaagggctacaaagaagtgaaaaaggacctgatcatcaagctgcctaagtactccctgttcgagctggaaaacggccggaagagaatgctggcctctgccggcgaactgcagaagggaaacgaactggccctgccctccaaatatgtgaacttcctgtacctggccagccactatgagaagctgaagggctcccccgaggataatgagcagaaacagctgtttgtggaacagcacaagcactacctggacgagatcatcgagcagatcagcgagttctccaagagagtgatcctggccgacgctaatctggacaaagtgctgtccgcctacaacaagcaccgggataagcccatcagagagcaggccgagaatatcatccacctgtttaccctgaccaatctgggagcccctgccgccttcaagtactttgacaccaccatcgaccggaagaggtacaccagcaccaaagaggtgctggacgccaccctgatccaccagagcatcaccggcctgtacgagacacggatcgacctgtctcagctgggaggcgacagcgctggaggaggtggaagcggaggaggaggaagcggaggaggaggtagcggacctaagaaaaagaggaaggtggcggccgctggatccggacgggctAAGGAATTCTACATCAGCATCGAGACCGTGGGTAACAACATCGTGGAAAGATATATTGACGAAAACGGCAAGGAGAGAACCAGAGAGGTGGAATACCTGCCTACAATGTTCCGGCACTGTAAAGAGGAATCCAAGTACAAGGATATCTACGGCAAAAACTGCGCCCCTCAGAAATTCCCCAGCATGAAAGACGCCAGAGATTGGATGAAGAGAATGGAGGATATCGGACTGGAAGCCCTGGGCATGAACGATTTCAAGCTGGCCTACATCTCCGATACATACGGAAGCGAGATCGTGTATGATAGAAAATTCGTGCGGGTGGCCAATTGTGACATTGAGGTGACCGGCGACAAGTTCCCTGATCCCATGAAAGCTGAATATGAGATCGACGCCATTACCCACTACGACAGCATCGACGACAGATTCTACGTGTTCGACCTGCTGAACTCCATGTACGGCAGCGTGTCCAAGTGGGACGCTAAGCTGGCCGCCAAGCTGGACTGCGAGGGCGGCGACGAGGTTCCACAAGAGATCCTGGACCGGGTCATCTACATGCCCTTCGACAACGAGAGGGACATGCTGATGGAATACATCAACCTGTGGGAGCAGAAGCGCCCCGCCATTTTTACAGGCTGGAACATCGAGGGCTTCGACGTGCCTTATATCATGAATAGAGTGAAAATGATCCTGGGAGAACGGAGCATGAAAAGATTCAGCCCTATCGGCAGAGTGAAGAGCAAGCTGATCCAAAACATGTACGGCTCCAAGGAAATCTATAGCATCGATGGCGTGTCCATCCTGGATTACCTGGACCTGTACAAAAAGTTCGCCTTCACCAACCTGCCATCTTTCTCTCTTGAGAGCGTCGCCCAGCACGAGACAAAGAAGGGCAAGCTGCCGTACGACGGTCCTATCAACAAGCTGAGAGAAACAAATCACCAGAGATACATCAGCTACAACATCATCGATGTGGAAAGCGTTCAGGCCATCGATAAAATCAGAGGCTTCATCGACCTGGTGCTGTCTATGTCTTACTACGCCAAGATGCCTTTTAGCGGAGTGATGAGCCCTATCAAGACCTGGGATGCCATCATCTTCAACAGCCTGAAGGGCGAACACAAGGTGATCCCCCAACAGGGCAGCCACGTGAAGCAGAGCTTCCCAGGCGCTTTTGTGTTCGAGCCCAAGCCCATAGCGCGGAGATACATCATGAGCTTTGATCTGACCAGCCTGTACCCCAGCATCATTCGGCAAGTGAACATTTCTCCAGAAACCATCAGAGGCCAGTTTAAGGTGCACCCTATCCACGAGTATATTGCAGGCACCGCTCCTAAACCTAGCGACGAGTACAGCTGCTCTCCTAACGGCTGGATGTACGACAAGCACCAGGAGGGAATCATCCCTAAGGAAATTGCCAAGGTGTTTTTCCAGCGGAAGGACTGGAAGAAAAAAATGTTCGCCGAGGAAATGAACGCCGAGGCCATCAAGAAGATCATCATGAAGGGCGCCGGCAGCTGCTCCACCAAGCCTGAGGTGGAAAGATACGTGAAGTTCAGCGACGATTTCCTGAATGAGCTCAGCAACTACACCGAGTCTGTCCTGAACTCACTGATTGAGGAATGCGAGAAGGCCGCCACCCTGGCTAATACCAACCAGCTGAACCGGAAGATTCTGATCAACAGCCTGTACGGAGCTCTGGGCAATATTCACTTCAGATACTACGATCTGCGAAACGCCACAGCTATTACAATTTTCGGCCAGGTGGGCATCCAGTGGATCGCCAGAAAGATCAATGAGTACCTGAACAAGGTGTGCGGCACCAACGACGAGGACTTCATCGCCGCTGGCGATACTGATAGCGTGTACGTTTGTGTGGACAAGGTCATCGAGAAGGTTGGCCTGGACAGATTTAAGGAACAGAACGACCTCGTGGAGTTCATGAACCAGTTCGGAAAGAAGAAGATGGAACCCATGATCGATGTGGCTTATAGAGAGCTGTGCGACTACATGAACAACAGAGAGCACCTGATGCACATGGATAGAGAAGCTATTTCTTGCCCTCCTCTGGGCTCTAAGGGAGTGGGCGGATTTTGGAAAGCCAAAAAGAGATACGCCCTGAATGTGTACGACATGGAAGATAAGAGATTCGCCGAGCCTCACCTGAAAATCATGGGCATGGAAACACAGCAGAGCAGCACCCCTAAGGCTGTGCAGGAGGCCCTGGAAGAGTCTATCCGGAGAATCTTGCAGGAGGGCGAGGAAAGCGTGCAGGAGTACTACAAGAACTTCGAGAAAGAATACAGACAGCTGGACTACAAGGTGATCGCGGAGGTGAAGACCGCTAATGATATCGCCAAGTACGACGACAAGGGCTGGCCCGGCTTCAAGTGCCCCTTCCACATCAGAGGCGTGCTCACCTACCGCAGAGCCGTTTCCGGCCTGGGCGTGGCCCCTATCCTGGATGGAAACAAAGTCATGGTGCTGCCTCTGAGAGAGGGCAACCCCTTTGGAGATAAATGCATCGCTTGGCCTAGCGGCACTGAGCTGCCCAAGGAAATCCGCTCCGACGTGCTGAGCTGGATCGATCACAGCACCCTGTTCCAAAAGTCCTTCGTGAAGCCCCTGGCCGGCATGTGCGAGTCCGCCGGCATGGACTACGAGGAAAAGGCCAGCCTGGATTTCCTGTTCGGC

T4 DNA polymerase-L-Cas9

atgccaaagaagaagcggaaggtcAAGGAATTCTACATCAGCATCGAGACCGTGGGTAACAACATCGTGGAAAGATATATTGACGAAAACGGCAAGGAGAGAACCAGAGAGGTGGAATACCTGCCTACAATGTTCCGGCACTGTAAAGAGGAATCCAAGTACAAGGATATCTACGGCAAAAACTGCGCCCCTCAGAAATTCCCCAGCATGAAAGACGCCAGAGATTGGATGAAGAGAATGGAGGATATCGGACTGGAAGCCCTGGGCATGAACGATTTCAAGCTGGCCTACATCTCCGATACATACGGAAGCGAGATCGTGTATGATAGAAAATTCGTGCGGGTGGCCAATTGTGACATTGAGGTGACCGGCGACAAGTTCCCTGATCCCATGAAAGCTGAATATGAGATCGACGCCATTACCCACTACGACAGCATCGACGACAGATTCTACGTGTTCGACCTGCTGAACTCCATGTACGGCAGCGTGTCCAAGTGGGACGCTAAGCTGGCCGCCAAGCTGGACTGCGAGGGCGGCGACGAGGTTCCACAAGAGATCCTGGACCGGGTCATCTACATGCCCTTCGACAACGAGAGGGACATGCTGATGGAATACATCAACCTGTGGGAGCAGAAGCGCCCCGCCATTTTTACAGGCTGGAACATCGAGGGCTTCGACGTGCCTTATATCATGAATAGAGTGAAAATGATCCTGGGAGAACGGAGCATGAAAAGATTCAGCCCTATCGGCAGAGTGAAGAGCAAGCTGATCCAAAACATGTACGGCTCCAAGGAAATCTATAGCATCGATGGCGTGTCCATCCTGGATTACCTGGACCTGTACAAAAAGTTCGCCTTCACCAACCTGCCATCTTTCTCTCTTGAGAGCGTCGCCCAGCACGAGACAAAGAAGGGCAAGCTGCCGTACGACGGTCCTATCAACAAGCTGAGAGAAACAAATCACCAGAGATACATCAGCTACAACATCATCGATGTGGAAAGCGTTCAGGCCATCGATAAAATCAGAGGCTTCATCGACCTGGTGCTGTCTATGTCTTACTACGCCAAGATGCCTTTTAGCGGAGTGATGAGCCCTATCAAGACCTGGGATGCCATCATCTTCAACAGCCTGAAGGGCGAACACAAGGTGATCCCCCAACAGGGCAGCCACGTGAAGCAGAGCTTCCCAGGCGCTTTTGTGTTCGAGCCCAAGCCCATAGCGCGGAGATACATCATGAGCTTTGATCTGACCAGCCTGTACCCCAGCATCATTCGGCAAGTGAACATTTCTCCAGAAACCATCAGAGGCCAGTTTAAGGTGCACCCTATCCACGAGTATATTGCAGGCACCGCTCCTAAACCTAGCGACGAGTACAGCTGCTCTCCTAACGGCTGGATGTACGACAAGCACCAGGAGGGAATCATCCCTAAGGAAATTGCCAAGGTGTTTTTCCAGCGGAAGGACTGGAAGAAAAAAATGTTCGCCGAGGAAATGAACGCCGAGGCCATCAAGAAGATCATCATGAAGGGCGCCGGCAGCTGCTCCACCAAGCCTGAGGTGGAAAGATACGTGAAGTTCAGCGACGATTTCCTGAATGAGCTCAGCAACTACACCGAGTCTGTCCTGAACTCACTGATTGAGGAATGCGAGAAGGCCGCCACCCTGGCTAATACCAACCAGCTGAACCGGAAGATTCTGATCAACAGCCTGTACGGAGCTCTGGGCAATATTCACTTCAGATACTACGATCTGCGAAACGCCACAGCTATTACAATTTTCGGCCAGGTGGGCATCCAGTGGATCGCCAGAAAGATCAATGAGTACCTGAACAAGGTGTGCGGCACCAACGACGAGGACTTCATCGCCGCTGGCGATACTGATAGCGTGTACGTTTGTGTGGACAAGGTCATCGAGAAGGTTGGCCTGGACAGATTTAAGGAACAGAACGACCTCGTGGAGTTCATGAACCAGTTCGGAAAGAAGAAGATGGAACCCATGATCGATGTGGCTTATAGAGAGCTGTGCGACTACATGAACAACAGAGAGCACCTGATGCACATGGATAGAGAAGCTATTTCTTGCCCTCCTCTGGGCTCTAAGGGAGTGGGCGGATTTTGGAAAGCCAAAAAGAGATACGCCCTGAATGTGTACGACATGGAAGATAAGAGATTCGCCGAGCCTCACCTGAAAATCATGGGCATGGAAACACAGCAGAGCAGCACCCCTAAGGCTGTGCAGGAGGCCCTGGAAGAGTCTATCCGGAGAATCTTGCAGGAGGGCGAGGAAAGCGTGCAGGAGTACTACAAGAACTTCGAGAAAGAATACAGACAGCTGGACTACAAGGTGATCGCGGAGGTGAAGACCGCTAATGATATCGCCAAGTACGACGACAAGGGCTGGCCCGGCTTCAAGTGCCCCTTCCACATCAGAGGCGTGCTCACCTACCGCAGAGCCGTTTCCGGCCTGGGCGTGGCCCCTATCCTGGATGGAAACAAAGTCATGGTGCTGCCTCTGAGAGAGGGCAACCCCTTTGGAGATAAATGCATCGCTTGGCCTAGCGGCACTGAGCTGCCCAAGGAAATCCGCTCCGACGTGCTGAGCTGGATCGATCACAGCACCCTGTTCCAAAAGTCCTTCGTGAAGCCCCTGGCCGGCATGTGCGAGTCCGCCGGCATGGACTACGAGGAAAAGGCCAGCCTGGATTTCCTGTTCGGCtctggaggatctagcggaggatcctctggcagcgagacaccaggaacaagcgagtcagcaacaccagagagcagtggcggcagcagcggcggcagcccaaagaagaagcggaaggtcggtatccacggagtcccagcagccgacaagaagtacagcatcggcctggacatcggcaccaactctgtgggctgggccgtgatcaccgacgagtacaaggtgcccagcaagaaattcaaggtgctgggcaacaccgaccggcacagcatcaagaagaacctgatcggagccctgctgttcgacagcggcgaaacagccgaggccacccggctgaagagaaccgccagaagaagatacaccagacggaagaaccggatctgctatctgcaagagatcttcagcaacgagatggccaaggtggacgacagcttcttccacagactggaagagtccttcctggtggaagaggataagaagcacgagcggcaccccatcttcggcaacatcgtggacgaggtggcctaccacgagaagtaccccaccatctaccacctgagaaagaaactggtggacagcaccgacaaggccgacctgcggctgatctatctggccctggcccacatgatcaagttccggggccacttcctgatcgagggcgacctgaaccccgacaacagcgacgtggacaagctgttcatccagctggtgcagacctacaaccagctgttcgaggaaaaccccatcaacgccagcggcgtggacgccaaggccatcctgtctgccagactgagcaagagcagacggctggaaaatctgatcgcccagctgcccggcgagaagaagaatggcctgttcggaaacctgattgccctgagcctgggcctgacccccaacttcaagagcaacttcgacctggccgaggatgccaaactgcagctgagcaaggacacctacgacgacgacctggacaacctgctggcccagatcggcgaccagtacgccgacctgtttctggccgccaagaacctgtccgacgccatcctgctgagcgacatcctgagagtgaacaccgagatcaccaaggcccccctgagcgcctctatgatcaagagatacgacgagcaccaccaggacctgaccctgctgaaagctctcgtgcggcagcagctgcctgagaagtacaaagagattttcttcgaccagagcaagaacggctacgccggctacattgacggcggagccagccaggaagagttctacaagttcatcaagcccatcctggaaaagatggacggcaccgaggaactgctcgtgaagctgaacagagaggacctgctgcggaagcagcggaccttcgacaacggcagcatcccccaccagatccacctgggagagctgcacgccattctgcggcggcaggaagatttttacccattcctgaaggacaaccgggaaaagatcgagaagatcctgaccttccgcatcccctactacgtgggccctctggccaggggaaacagcagattcgcctggatgaccagaaagagcgaggaaaccatcaccccctggaacttcgaggaagtggtggacaagggcgcttccgcccagagcttcatcgagcggatgaccaacttcgataagaacctgcccaacgagaaggtgctgcccaagcacagcctgctgtacgagtacttcaccgtgtataacgagctgaccaaagtgaaatacgtgaccgagggaatgagaaagcccgccttcctgagcggcgagcagaaaaaggccatcgtggacctgctgttcaagaccaaccggaaagtgaccgtgaagcagctgaaagaggactacttcaagaaaatcgagtgcttcgactccgtggaaatctccggcgtggaagatcggttcaacgcctccctgggcacataccacgatctgctgaaaattatcaaggacaaggacttcctggacaatgaggaaaacgaggacattctggaagatatcgtgctgaccctgacactgtttgaggacagagagatgatcgaggaacggctgaaaacctatgcccacctgttcgacgacaaagtgatgaagcagctgaagcggcggagatacaccggctggggcaggctgagccggaagctgatcaacggcatccgggacaagcagtccggcaagacaatcctggatttcctgaagtccgacggcttcgccaacagaaacttcatgcagctgatccacgacgacagcctgacctttaaagaggacatccagaaagcccaggtgtccggccagggcgatagcctgcacgagcacattgccaatctggccggcagccccgccattaagaagggcatcctgcagacagtgaaggtggtggacgagctcgtgaaagtgatgggccggcacaagcccgagaacatcgtgatcgaaatggccagagagaaccagaccacccagaagggacagaagaacagccgcgagagaatgaagcggatcgaagagggcatcaaagagctgggcagccagatcctgaaagaacaccccgtggaaaacacccagctgcagaacgagaagctgtacctgtactacctgcagaatgggcgggatatgtacgtggaccaggaactggacatcaaccggctgtccgactacgatgtggaccatatcgtgcctcagagctttctgaaggacgactccatcgacaacaaggtgctgaccagaagcgacaagaaccggggcaagagcgacaacgtgccctccgaagaggtcgtgaagaagatgaagaactactggcggcagctgctgaacgccaagctgattacccagagaaagttcgacaatctgaccaaggccgagagaggcggcctgagcgaactggataaggccggcttcatcaagagacagctggtggaaacccggcagatcacaaagcacgtggcacagatcctggactcccggatgaacactaagtacgacgagaatgacaagctgatccgggaagtgaaagtgatcaccctgaagtccaagctggtgtccgatttccggaaggatttccagttttacaaagtgcgcgagatcaacaactaccaccacgcccacgacgcctacctgaacgccgtcgtgggaaccgccctgatcaaaaagtaccctaagctggaaagcgagttcgtgtacggcgactacaaggtgtacgacgtgcggaagatgatcgccaagagcgagcaggaaatcggcaaggctaccgccaagtacttcttctacagcaacatcatgaactttttcaagaccgagattaccctggccaacggcgagatccggaagcggcctctgatcgagacaaacggcgaaaccggggagatcgtgtgggataagggccgggattttgccaccgtgcggaaagtgctgagcatgccccaagtgaatatcgtgaaaaagaccgaggtgcagacaggcggcttcagcaaagagtctatcctgcccaagaggaacagcgataagctgatcgccagaaagaaggactgggaccctaagaagtacggcggcttcgacagccccaccgtggcctattctgtgctggtggtggccaaagtggaaaagggcaagtccaagaaactgaagagtgtgaaagagctgctggggatcaccatcatggaaagaagcagcttcgagaagaatcccatcgactttctggaagccaagggctacaaagaagtgaaaaaggacctgatcatcaagctgcctaagtactccctgttcgagctggaaaacggccggaagagaatgctggcctctgccggcgaactgcagaagggaaacgaactggccctgccctccaaatatgtgaacttcctgtacctggccagccactatgagaagctgaagggctcccccgaggataatgagcagaaacagctgtttgtggaacagcacaagcactacctggacgagatcatcgagcagatcagcgagttctccaagagagtgatcctggccgacgctaatctggacaaagtgctgtccgcctacaacaagcaccgggataagcccatcagagagcaggccgagaatatcatccacctgtttaccctgaccaatctgggagcccctgccgccttcaagtactttgacaccaccatcgaccggaagaggtacaccagcaccaaagaggtgctggacgccaccctgatccaccagagcatcaccggcctgtacgagacacggatcgacctgtctcagctgggaggcgacaaaaggccggcggccacgaaaaaggccggccaggcaaaaaagaaaaag
